# Abscisic acid-mediated water stress regulation can mechanistically explain oscillations and water stress memory in stomatal conductance

**DOI:** 10.64898/2025.12.26.696581

**Authors:** Sahil Desai, Abraham Stroock

## Abstract

Stomatal pores, formed by guard cells, govern the critical trade-off between carbon assimilation and water loss in plants. Their dynamic responses to environmental stresses, such as stomatal oscillations and drought “stress memory” (hysteresis), have lacked a unified mechanistic explanation. While abscisic acid (ABA) is believed to play key roles in water stress responses, no model has linked its core regulatory kinetics to these complex stomatal behaviors. Here, we introduce a coupled hydropassive-hydroactive (HP-HA) model that integrates leaf hydraulics with the biokinetics of guard cell-autonomous ABA regulation and plasma membrane-mediated osmoregulation. We demonstrate that this framework predicts accurate, genotype-specific stomatal regulation across wildtype, ABA-insensitive mutant (*ost1-3*), and ABA-synthesis mutant (*aao3-2*) in *Arabidopsis thaliana* (*At*) and that non-linear feedbacks in ABA autoregulation can drive both stomatal oscillations and hysteresis. This work unifies genetic, signaling, and membrane processes with leaf-scale physiological dynamics, providing a new predictive foundation for understanding and modulating plant management of water use and water stress.

## Introduction

Stomata are pores on the surface of a leaf formed by guard cells (Fig. 1a). Changes in the apertures of these pores change their diffusive resistance (1/*g*_*s*_ [s. m^2^/mmol], where *g*_*s*_ [mmol/m^2^/s] is the stomatal conductance) to the exchange of gases between the leaf and the atmosphere. The pores mediate both the uptake of carbon dioxide (CO_2_) and the loss of water vapor (H_2_O) without strong selectivity. At the leaf level, the stomatal regulation of water flux controls water-use and leaf temperature via transpiration rate (E) and the regulation of CO_2_ flux influences the rate of carbon assimilation by photosynthesis (A)^1^. The coupled regulation of the fluxes of water vapor and carbon dioxide by stomates impacts instantaneous water-use efficiency (A/E), growth, productivity, and heat and drought tolerance in natural and agricultural contexts^2,3^. The stomatal resistance varies with changes in water availability (xylem water potential, *ψ*_*xyl*_ [MPa]), water demand (VPD [kPa]), concentrations of CO_2_, intensity of photosynthetically active light, and with time of day and season^4^.

**Figure 1.**
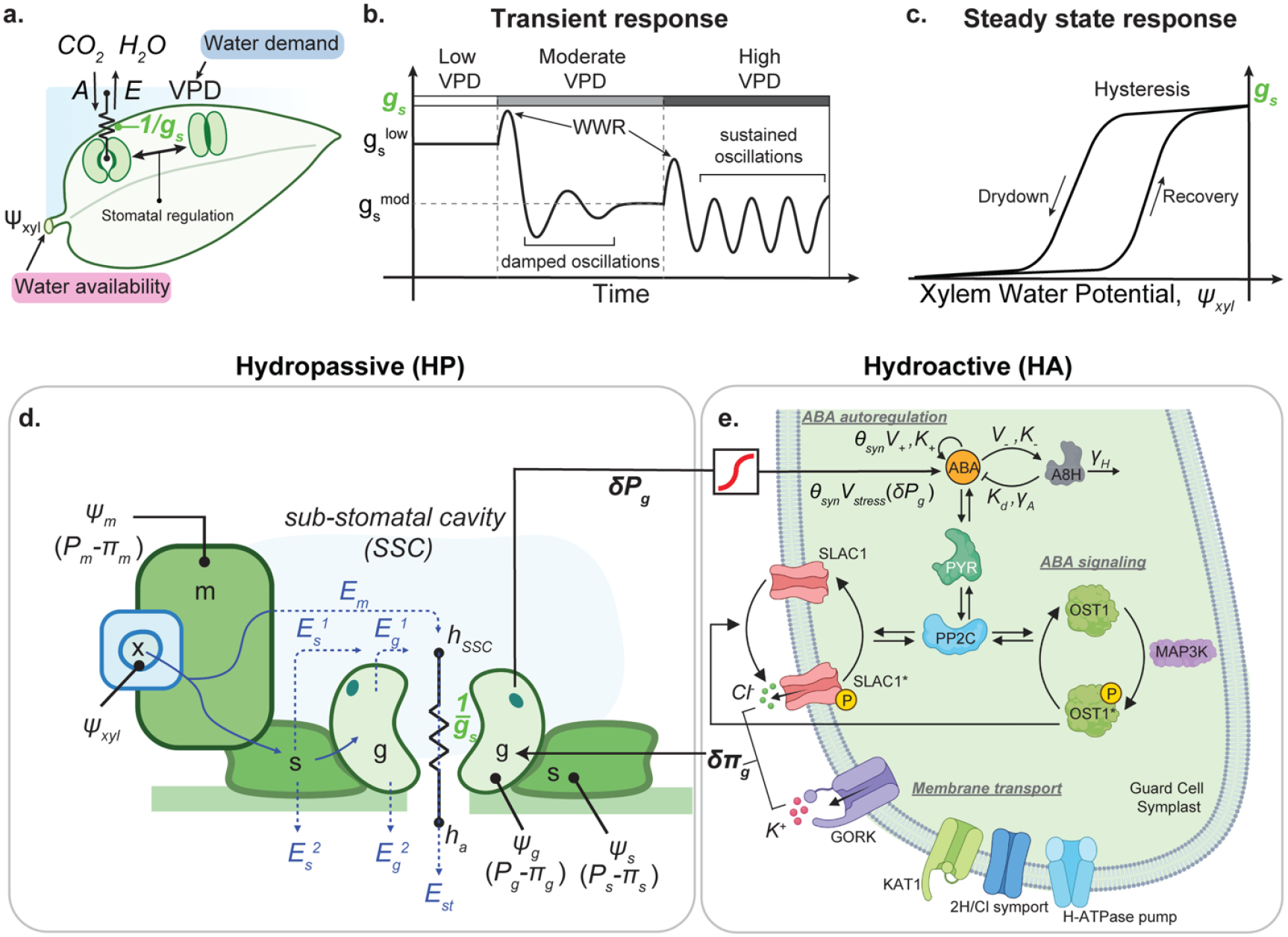
Multi-scale model of stomatal response to water stress. **a**. Schematic diagram of a leaf with exchange of carbon dioxide (*CO*_2_ ) and water vapor (*H*_2_ *O*) with the atmosphere regulated by stomates. The rate of assimilation (flux of *CO*_2_, *A* [*µmol/m*^2^*/s*]) and the rate of transpiration (flux of *H*_2_ *O, E* [*mmol/m*^2^*/s*]) are controlled by stomatal aperture and the stomatal resistance (*1/g*_*s*_ [*m*^2^. *s/mmol*]) defined by this aperture. Stomatal resistance is the reciprocal of stomatal conductance (*g*_*s*_ [*mmol/ m*^2^. *s*]). Water availability is represented by water potential in the xylem, *ψ*_*xyl*_ [MPa]; water demand is represented by the vapor pressure deficit in the atmosphere, VPD. **b**. Schematic representation of transient responses of stomatal conductance to changes in VPD: for a step change to “Moderate VPD”, stomatal conductance decreases with increasing VPD, exhibiting damped oscillations; for a step change to “High VPD”, stomatal conductance can exhibit sustained oscillations. For each drop in VPD, stomatal conductance first shows a characteristic increase called a wrong way response (WWR). **c**. Schematic representation of steady state response of stomata to water availability can exhibit hysteresis under cycles of drought and recovery. Stomatal conductance exhibits stress memory by maintaining a higher equipotential value during drydown (xylem potential, *ψ*_*xyl*_ decreasing) versus during recovery (*ψ*_*xyl*_ increasing). **d**. Schematic diagram of leaf cross-section showing xylem (x), mesophyll (m), subsidiary cells (s), guard cells (g), and sub-stomatal cavity (SSC). The hydraulics of bundle sheath cells are included in the mesophyll; the hydraulics of other epidermal cells are included in the subsidiary cells. The water fluxes pass as a liquid (blue curves) from the xylem to irrigate the mesophyll, subsidiary, and guard cells and to sites of evaporation. Vapor fluxes (dashed blue curves) pass into the SSC (relative humidity, *h*_*SSC*_ ) from the mesophyll (*E*_*m*_ ), subsidiary 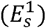, and guard 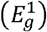, cells and contribute to the total flux (*E*_*st*_ ) limited by the stomatal resistance (1/*g*_*s*_ ); vapor flux also leaves the subsidiary 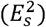 and guard 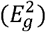 cells directly into the atmosphere at the leaf surface (relative humidity, *h*_*a*_ ). The model tracks the xylem water potential, *ψ*_*xyl*_, turgor pressures (*P*_*m*_, *P*_*s*_, *and P*_*g*_) and osmotic potentials (*π*_*m*_, *π*_*s*_, *and π*_*g*_) of the mesophyll, subsidiary, and guard cells. See SI section S2 for details. **e**. Schematic diagram of the abscisic acid (ABA)-mediated pathway for stomatal closure. Changes in turgor pressure in guard cells (δ*P*_*g*_ [*MPa*]) upregulates stress-induced *de novo* biosynthesis of ABA^38–40^, characterized by the rate *V*_*stress*_ (δ*P*_*g*_) [*nM/s*]. ABA increases its own production (*V*_+_ [*nM*/*s*], *K*_+_ [*nM*]) and its own degradation via upregulation of the CYP707A gene family which encodes the enzyme 8’-hydroxylase (A8H)^43,53–55^ (*V*_−_ [*nM/s*],*K*_−_ [*nM*]). The strength of ABA’s synthesis is determined by *θ*_*syn*_ . This enzyme catalyzes ABA catabolism following Michaelis-Menten kinetics ( *K*_*d*_ [*nM*], *γ*_*A*_ [1/*s*]), and undergoes its own degradation (*γ*_*H*_ [1/*s*]). Positive regulation is represented by pointed arrows, while inhibitory interactions are indicated by flat arrows. ABA binds its receptor PYR to form the ABA:PYR complex^58,59^. This complex non-competitively inhibits phosphatase PP2C by binding allosteric sites^60^. PP2C regulates a bicyclic cascade of futile cycles: in the first PP2C dephosphorylates protein kinase OST1* into its inactive form, OST1, while MAP3K phosphorylates OST1 back into its active form^61^, OST1*; in the second, PP2C dephosphorylates the open form of the anion channel, SLAC1* to its closed form, SLAC1 while OST1* phosphorylates SLAC1 into its open state, SLAC1*^48,49^. The activation of SLAC1* with elevated ABA facilitates the efflux of *Cl*^−^ ions, leading to depolarization of the guard cell plasma membrane^41^. This depolarization opens the GORK cation channel, allowing *K*^+^efflux and resulting in a decrease in guard cells osmotic potential^41^, δ *π*_*g*_ [*MPa*]. This change in osmotic potential in guard cells feeds back into change in guard cell turgor pressure and stomatal conductance. This model also incorporates additional key transporters, including KAT1 *K*^+^ channel, H-ATPase pump, and 2H/Cl symporter, which regulate ion fluxes and maintain guard cell homeostasis. See SI section S3 for details.

Both the steady state^5–8^ and transient responses^9–16^ of *g*_*s*_ to these parameters can influence plant function and provide insights into the underlying physiological processes that regulate stomatal aperture. In angiosperms, with respect to variations in water demand (Fig. 1b), one observes decreasing steady state values of *g*_*s*_ for increasing VPD 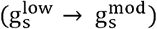 and characteristic transient responses to abrupt changes that include wrong-way response (WWR), damped oscillations, and sustained oscillation^16–20^. With respect to variations in water availability, *g*_*s*_ decreases with decreasing *ψ*_*xyl*_, and, for cycles of drought and recovery, stomatal conductance can show “stress memory” or hysteresis, with higher values of *g*_*s*_ during drydown than during recovery for the same values of *ψ*_*xyl*_ (Fig. 1c)^21–23^.

Despite the recognized importance of stomatal regulation, we lack a complete understanding of the mechanisms by which such variations in stomatal conductance emerge in response to variations in environment conditions^24–27^. Here, we focus on mechanistic mathematical models of stomatal responses to water availability (*ψ*_*xyl*_ ) and demand (VPD).

Following the seminal work of Cowan^9^, investigators have explored various hydraulic architectures to represent hydropassive (HP) dynamics, with explicit accounting for the resistance to flow, tissue capacitance, and turgor and osmotic pressures along the path between xylem and guard cells^7,9–11^. Cowan^9^ and Delwiche^10^ explored transient responses in these HP models and identified parameter regimes that led to damped and sustained oscillations in *g*_*s*_ in response to abrupt changes in VPD. Buckley later refined this HP framework and found that, with physiologically realistic parameters, it could not reproduce observed water-stress responses^7,11,25^. Peak and Mott presented an alternative HP scenario in which guard cells are irrigated via coupling to the vapor in the sub-stomatal cavities^28^. This scenario has been called into question by the recent elucidation of large vapor undersaturation within leaves^29–31^.

Several groups have coupled HP and HA processes in stomatal models. Cowan first explored HA processes within his HP framework by imposing time-dependent changes in osmotic potential within the guard cells, capturing guard cell-localized osmoregulation without explicit molecular details^9^. In his refined HP framework, Buckley explored osmoregulation driven by photosynthetic activity^7^. A team led by Blatt developed a coupled HP-HA model^14–16^ with the HP scenario of Peak and Mott^28^ and their own detailed mechanistic HA model (“OnGuard”) that accounts for biokinetics of membrane-mediated and metabolic processes that control osmotic adjustment; they account explicitly for biochemical responses to light and CO_2_ but not directly to water status via ABA or other pathways. They showed qualitative agreement of their passive responses to water stress and suggested that ABA may not be required for stomatal response to water status^16^. Notably, they did not predict damped or persistent oscillations or hysteresis (Fig. 1b-c). Investigating the mechanisms of temporal oscillations and spatial patchiness, Mott and co-authors have focused on the tissue-scale coupling between stomates^32–36^. Recently, Peak and Mott^36^ integrated an HP model of stomate-stomate coupling mediated by hydraulic and vapor paths with an HA model of the biokinetics of osmotic responses to light and internal concentrations of CO_2_. This model predicted WWR and damped oscillations in response to a step change in VPD and persistent oscillations in response to a transition to red light; they did not confront their model directly with any measurements. See SI section S1 for additional discussion of the background on models.

In parallel, numerous labs have elucidated gene regulatory, signaling, and membrane-mediated processes that occur in leaf tissues and guard cells in response to water stress and play roles in regulating these cells’ state of turgor and thus also stomatal aperture and *g*_*s*_ ^17,37–42^. In this literature, we note that: (i) ABA has been identified as an important signal for HA regulation^25,43–46^, without convergence on its localization (i.e., to guard cells or adjacent tissues)^38–40^, or its necessity^16,47^; (ii) osmoregulation mediated by SLAC1 anion channels has been identified as a critical, guard-cell specific process downstream of ABA signaling that is involved in stomatal closure^48–50^; and (iii) no pathway of mechanotransduction coupling cell and tissue water status to active cell and tissue responses has yet been identified^40,51,52^.

To date, no one has proposed a coupled HP-HA model that accounts explicitly for the ABA-mediated responses to water status in guard cells that could play roles in stomatal regulation. Here, we develop a coupled HP-HA model that mechanistically integrates canonical leaf hydraulics (Fig. 1d)^7,11–13^ with a biokinetic model of ABA processes within the guard cells (Fig. 1e)^37–46,48–50,53–61^. This HA model includes an empirical turgor-dependent relation for ABA synthesis^38–40^, and, critically, the explicit biokinetics of ABA autoregulation^43,53–57^, ABA signaling^37,42,44,45,48–50,58–61^, and membrane-mediated osmoregulation processes^41,50^. We show that ABA autoregulation, when coupled with HP dynamics, provides the core non-linearity to drive stomatal responses to VPD and *ψ*_*xyl*_, including the key emergent phenomena of stomatal oscillations (Fig. 1b) and the bi-stability required for hysteresis (Fig. 1c). See Methods and SI sections S2-S5 and S7 for details on the formulation and parameterization of this model.

## Results

We parametrized our model (Fig. 2a) by confronting its predictions with measurements reported by Merilo^17^ of stomatal response to a step change in VPD in three genetic variants of *Arabidopsis thaliana* (*At*): *At* wildtype (Col-0, “wt” – green circles and curve), an *At* ABA signaling mutant (*ost1-3*, “sig” – blue triangles and curve), and *At* ABA synthesis mutant (*aao3-2*, “syn” – red diamonds and curve) shown in Fig. 2b.

**Figure 2.**
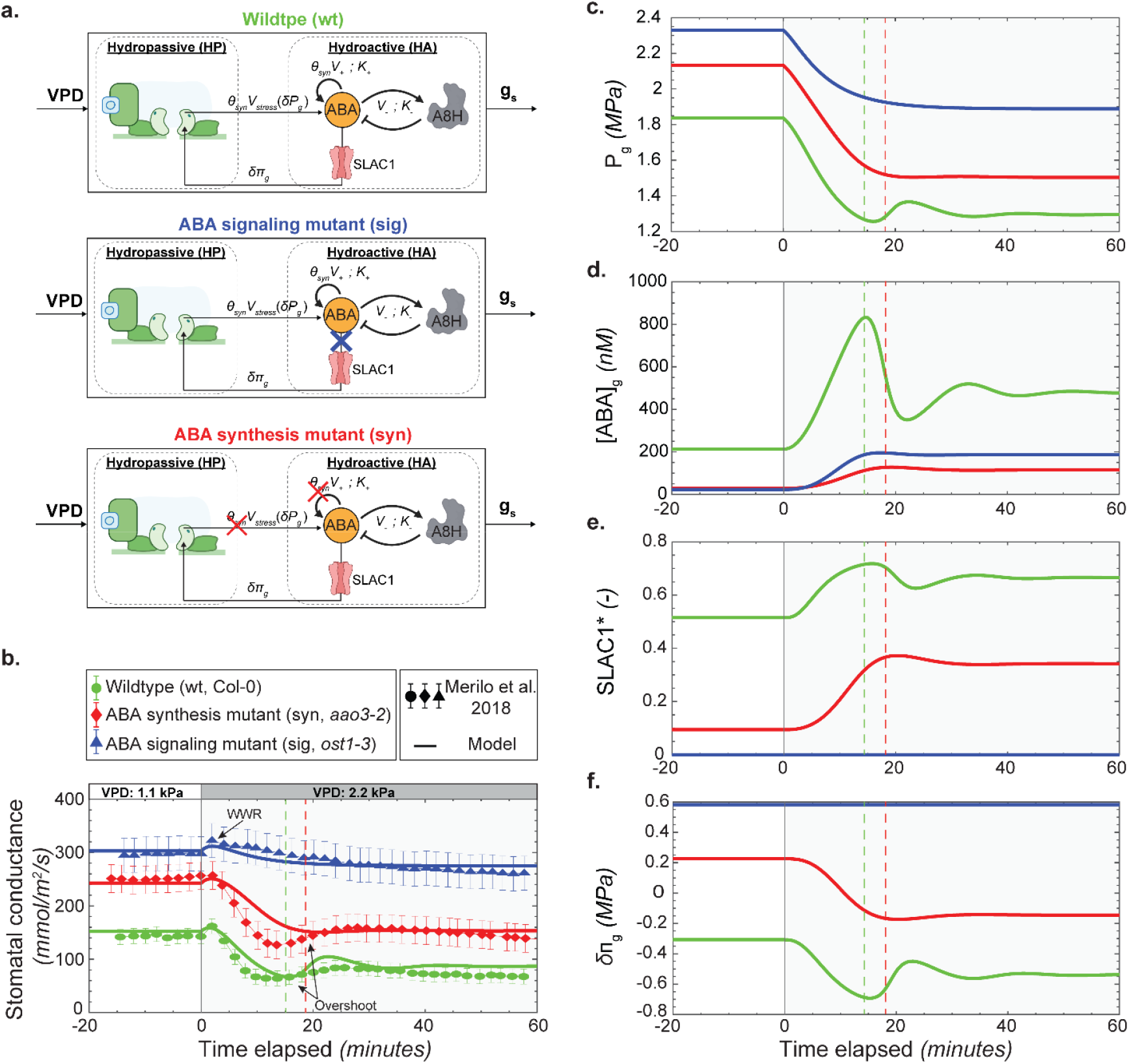
Transient stomatal responses to changes in vapor pressure deficit (VPD) across genetic variants. **a.** Schematics of the coupled model variants applied to Wildtype (wt), ABA signaling mutant (sig), and ABA synthesis mutant (syn). The schematic shows the hydropassive (HP) and hydroactive (HA) components as defined in Fig. 1. In wt, the HA component features the intact ABA autoregulation motif driving SLAC1 activation. In sig, the pathway is disrupted downstream of ABA autoregulation but upstream of channel activation (blue cross), representing diminished OST1 function. In syn, the ABA production is attenuated (red cross) via a reduction in *θ*_*syn*_ . **b**. Predicted and measured transients for a step change in VPD from 1.1 kPa to 2.2 kPa for: stomatal conductance, *g*_*s*_. The trajectories (solid curves) and data (symbols in (**b**)) are for representations of wildtype (wt, Col-0 – green curves and circles), an ABA synthesis mutant in this same background (syn, *aao3-2* – red curves and diamonds), and an ABA signaling mutant (sig, *ost1-3* – blue curves and triangles). Data are from Merilo^17^ for these genetic variants and this same perturbation in VPD. **c.-f**. Predicted transients corresponding to those in (**b**) for: guard cell turgor, *P*_*g*_ (**c**), concentration of ABA in guard cells, [*ABA*]_*g*_ (**d**), fractional activation of anion channel, SLAC1* (**e**), and osmotic potential change in guard cells, *δ π*_*g*_ (**f**). This step in VPD corresponds to a relative humidity (RH) change from 80% to 60%, at 36°C. Parameters for the ABA motif were: *K*_+_ = *K*_−_= 1560 *nM, V*_*stress*_ = 8.85 *nM*/*s, V*_+_ = 67.6 *nM*/*s, V* ^−^ = 1.5 *nM*/*s* with *θ*_*syn*_ =1 for wildtype and *θ*_*syn*_ = 0.12 for synthesis mutant. See main text and SI section S3 for details on model and representations of genetic variants.

### Hydropassive framework can explain the features of an ABA signaling mutant

For the HP portion of our model, we modified the framework of Buckley^7,11–13^ to make the calculations of evaporative fluxes from mesophyll (*E*_*m*_ ), subsidiary cell (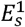 and 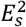), and guard cell (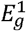 and 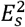) explicit and allow for internal undersaturation (*h*_*ssc*_ < 1) (Fig. 1d) (SI section S2.A and SI Table S7). Assuming the responses of the signaling mutant (Fig. 2a-middle; Fig. 2b, blue triangles) were dominated by hydropassive processes, we used this transient to adjust hydraulic parameters (hydraulic resistances, capacitances, osmotic potentials, evaporative resistances, and mechanics of guard and subsidiary cells - Fig. 1d) within ranges defined by the literature^7,11–13^.

Setting the total concentration of OST1 protein three orders of magnitude lower than the wildtype concentration (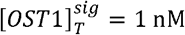 vs.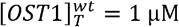) in our HA biokinetic model (Fig. 2a, middle), the predicted, HP-dominated responses (Fig. 2b, blue curve) agree within uncertainty with the measurements (Fig. 2b, blue triangles). Notably, the predictions capture high initial steady state values of *g*_*s*_ (*t*< 0) and weak loss of steady state conductance in response to increasing water demand (*g*_*s*_ (*t* ≤ 0 min) ≅ 300; *g*_*s*_ (*t* = 60 min) ≅ 280). They also capture a short-lived WWR immediately after the imposed change in *VPD* that arises due to the opposing effect of subsidiary cell turgor (*P*_*s*_ ) and guard cell turgor (*P*_*g*_) on stomatal aperture and the phase lag between these pressures (*P*_*s*_ responds before *P*_*g*_), as observed previously^9,10^. We conclude, as did Buckley^7,11,24,25^, that HP processes can explain the characteristic WWR observed in angiosperms but cannot explain the magnitude or details (e.g., damped oscillations) of responses to change in evaporative demand.

### ABA dynamics control the stomatal response in hydroactive genotypes

Using the HP parameters established above, we next examined predictions of the guard-cell-localized, ABA-mediated HA processes in our coupled HP–HA framework for wildtype responses (Fig. 2a-top) and for a synthesis mutant (Fig. 2a-bottom). Key details of the HA model are provided in Methods and SI section S3. We identify the rate processes of ABA autoregulation as the key control points for both biochemical and stomatal responses (SI Eqs. S53 and S56). The dynamics are governed by the interplay of: (i) synthesis driven by stress (*V*_*stress*_ ), (ii) positive feedback of ABA on its own synthesis (*V*_+_ ), and (iii) negative feedback of ABA on itself via A8H (*V* _−_). Importantly, the parameters *V*_−_ and ratio *V*_+_ */V*_*stress*_ govern whether the model predicts qualitative features such as damped or sustained oscillations and hysteresis in stomatal conductance (SI Fig. S6, S7, and S8). See SI sections S5.B-D for details.

To select parameter values, we confront the predictions of this model of ABA dynamics with transient measurements of *g*_*s*_ from Merilo^17^ for wildtype (wt – green circles in Fig. 2b) and an *aao3-2* mutant (syn – red diamonds in Fig. 2b). We model the *aao3-2* mutant as having a weaker total synthesis rate (indicated by the red cross in Fig. 2a-bottom; see *θ*_*syn*_ in Eq. S53 that modulates rates of ABA synthesis by both stress (*V*_*stress*_ ) and positive autoregulation (*V*_+_)) and weaker cooperativity in the ABA biosynthetic pathway relative to wildtype with all other parameters the same between the two. We present predictions for a set of parameters that provide quantitative agreement with steady state conductance for both genotypes (i.e., green and red curves for *g*_*s*_ (*t* ≤0 min) and *g*_*s*_ (*t* = 60 min) in Fig. 2b) and qualitative agreement with features of the transients, particularly the damped oscillations in the wildtype plants. Notably, based on comparisons with this data and due to the non-linearity of the governing equations, we cannot exclude other parameter values. In SI Fig. S10, we present satisfactory predictions with an alternative set of parameters. As we will discuss further below, we identify the parameterization used in Fig. 2 as preferred based on the predictions it yields with stronger evaporative demand.

To gain mechanistic insight into the predicted dynamics, we plot the trajectories of turgor pressure (*P*_*g*_, Fig. 2c), ABA concentration ([*ABA*]_*g*_, Fig. 2d), activated fraction of SLAC1 (SLAC1*, Fig. 2e), and change in osmotic potential (*δ π*_*g*_, Fig. 2f) in the guard cells. Focusing on wildtype dynamics (green curves), we see the following progression: (i) VPD acts through HP processes to lower *P*_*g*_ (Fig. 2c); (ii) the drop in *P*_*g*_ initiates synthesis of ABA (Fig. 2d); (iii) rising concentration of ABA drives its own degradation such that it quickly reaches a maximum (marked with dashed green vertical line) and begins to fall. The observed four-fold change in guard cell ABA levels in the nanomolar range aligns with estimates reported in literature^62^; and (iv) the time for activated SLAC1 (Fig. 2e) and osmotic potential (Fig. 2f) to rise to their maxima matches that for [*ABA*]_*g*_ (Fig. 2d) as does the time to the first minimum in *g*_*s*_ (Fig. 2b). We conclude that the rate and feedback encoded in ABA autoregulation plays the dominant role in defining the HA process in this model, with ABA signaling to SLAC1 and membrane-mediated osmoregulation both occurring rapidly relative to ABA synthesis and degradation. This predicted damped oscillatory response is consistent with gas exchange measurements following step changes in humidity^16–19^. While damping in oscillations can be a signature of ABA-mediated responses, our ABA model predicts that such responses can also occur without oscillations, a result consistent with other experimental observations^20^ (see Fig S9 role of *V*_−_ in controlling oscillations). See SI section S5.E and Fig. S9 for additional exploration of speed of stomatal closure and comparison with experiments from Merilo^17^ and the important origins of the predicted dynamics.

### Oscillations in stomatal conductance emerge from ABA autoregulatory motif

In Fig. 3, we follow previous modeling efforts^9,10,36,63^ in using the qualitative change in stomatal dynamics from damped to persistence oscillations to identify a source of non-linearity in the governing processes. Cowan^9^ presented an HP model in which the only non-linearity came from the coupled heat and mass transfer from leaf’s interior to atmosphere. In contrast, Peak and Mott^36^ presented multi-stomate models in which they include non-linear, hydropassive coupling between guard cells and light- and CO_2_-dependent processes in individual guard cells. They predict the damped oscillation in response to a step increase in VPD and oscillations after a transition from white to red light; they did not report a transition from damped to persistent oscillations for a single type of perturbation. Neither of these studies provided a mechanistic explanation for this experimentally observed phenomenon (Fig. S14); instead, they only described the transition qualitatively using simplified hydraulic representations with parameter estimates disconnected from the underlying biological processes.

**Figure 3.**
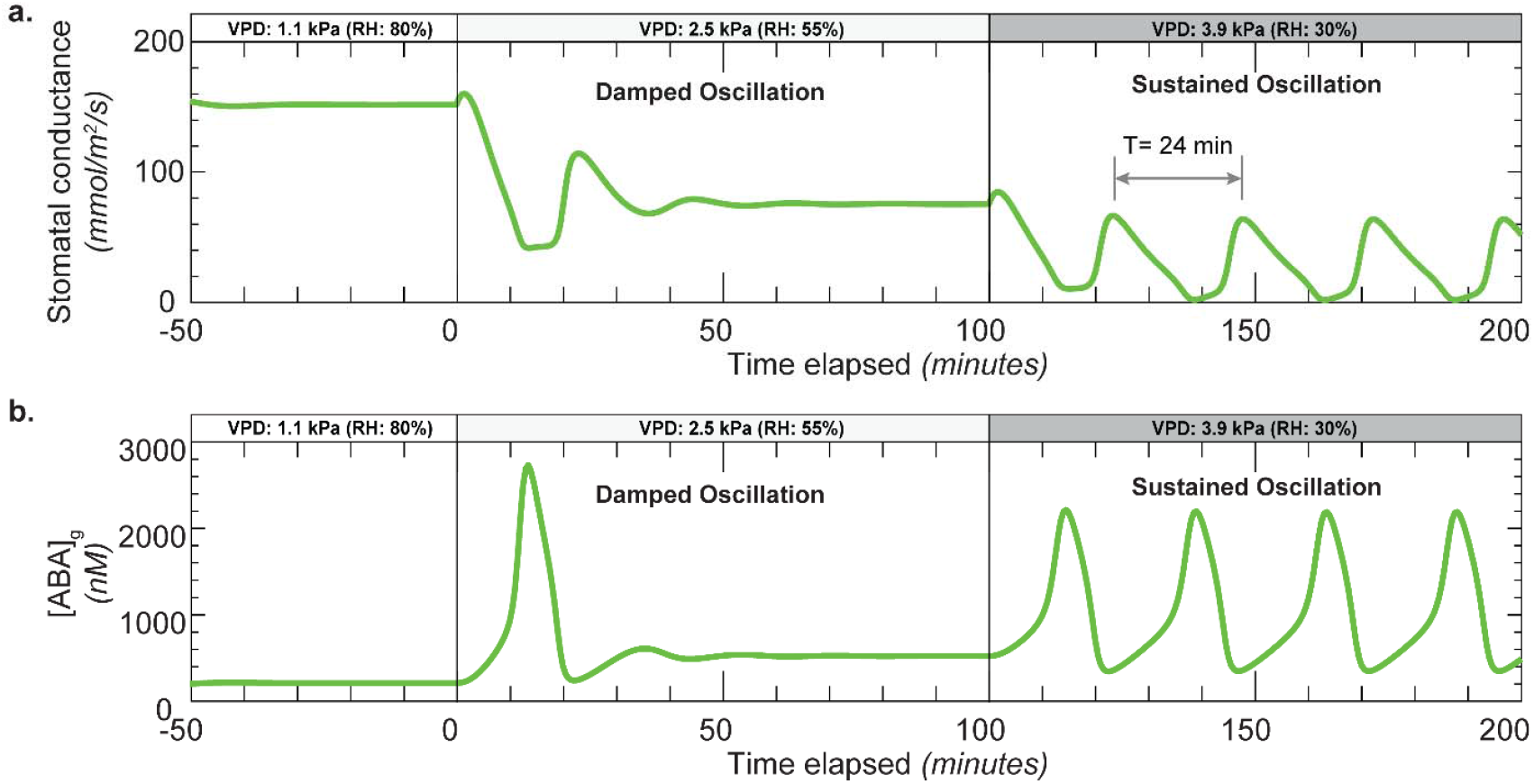
Emergence of oscillations in stomatal conductance with increasing vapor pressure deficit (VPD). **a.-b.** Predicted transients of concentrations of stomatal conductance (**a**) and ABA concentration in guard cell, [*ABA*]_*g*_ (**b**) for a series of step changes in VPD from 1.1kPa to 2.5 kPa at *t* = 0 mins and from VPD 2.5 kPa to 3.9 kPa at *t* = 100 mins, corresponding to a RH decrease from 80% to 55% and to 30% at 36°C, respectively. See main text and SI section S5 for additional information on selection and significance of model parameters. Parameters used were the same as those used in Fig. 2.

Here, we use our coupled HP-HA model parameterized to match the dynamics of wildtype *At*^17^ (Fig. 2b) to explore the transient responses of stomata to successive step increases in VPD of equal size (Fig. 3). Fig. S15 presents an experimental example of this type of multi-step transition. After a first step increase in VPD from 1.1 kPa to 2.5 kPa (80% to 55% RH at 36℃) at *t* = 0 mins, we predict damped oscillations in *g*_*s*_ (Fig. 3a) and guard cell ABA, [*ABA*]_*g*_ (Fig. 3b); and after a second step increase in VPD from 2.5 kPa to 3.9 kPa (55% to 30% RH at 36℃) at *t* = 100 mins, we predict sustained oscillations in *g*_*s*_ and [*ABA*]_*g*_.

To understand the basis of this transition to sustained oscillations, we have explored variants in our coupled HP-HA model. Importantly, we find that the fully coupled model shows this transition (Fig. S12a and S13a) whereas if we cut ABA-regulated osmoregulation (as for *ost1* mutant) both damped (Fig. S11b) and persistent (Fig. S12b) oscillations are eliminated, even if an equivalent perturbation is imposed on the HP dynamics via a step change in osmotic potential in the guard cells (Fig. S12c and S13c). Thus, the HP processes in our model cannot explain the observed tendency toward oscillations. On the other hand, when we directly perturb the concentration of ABA in the guard cell by imposing a perturbation in its stress dependent synthesis rate *V*_*stress*_, we retrieve both damped (Fig. S12d) and persistent (Fig. S13d) oscillations. Additionally, we find, as suggested by Cowan^9^ that VPD acts as an amplifier of non-linear response that leads to oscillations, the effect of which can be modulated by the strength of mechanobiological transduction (*V*_*stress*_ (*δP*_*g*_)) in Figs. 1d-e; Fig. S7), and by the negative feedback parameter, *V*_−_ that controls ABA catabolism via A8H (Fig. 1e; Fig. S8 d.-f.). We conclude that ABA autoregulation provides critical non-linear feedback that, in our framework, explains the transition from damped to persistent oscillations with increasing evaporative demand.

### Hysteresis in stomatal conductance emerges from ABA autoregulation

We now explore whether the non-linearity encoded in ABA autoregulation can capture another important feature of stomatal regulation, namely the hysteretic response in steady state stomatal aperture to cycles of water availability (drydown and recovery); this effect has been proposed to serve as a stress memory that can protect plants from recurrent drought (Fig. 1c)^21–23^. To generate the bi-stability (i.e., having two steady state solutions^64–66^) required for hysteresis, we utilized a distinct parameterization by strengthening ABA’s positive feedback gain (*V*_+_*/V*_*stress*_ ) 2-fold (Fig. 4a). This adjustment could represent a physiological shift from a “naïve” (monostable) to a “primed” (bi-stable) state (Fig. S6). While the system parametrized for transients (Fig. 2 and 3) operates in a monostable regime, experimental evidence confirms that prolonged drought can create the primed, bi-stable state via epigenetic remodeling^21,67,68^. This remodeling results in increased-induction of key ABA synthesis genes upon subsequent stress exposure^22^, thereby increasing the gain of the positive feedback loop in ABA autoregulation. With this adaptation of parameters, predictions of the model still show reasonable agreement with wildtype stomatal dynamics (Fig. S15) but do not show persistent oscillations at elevated VPD. See SI section S6.B for additional discussion.

**Figure 4.**
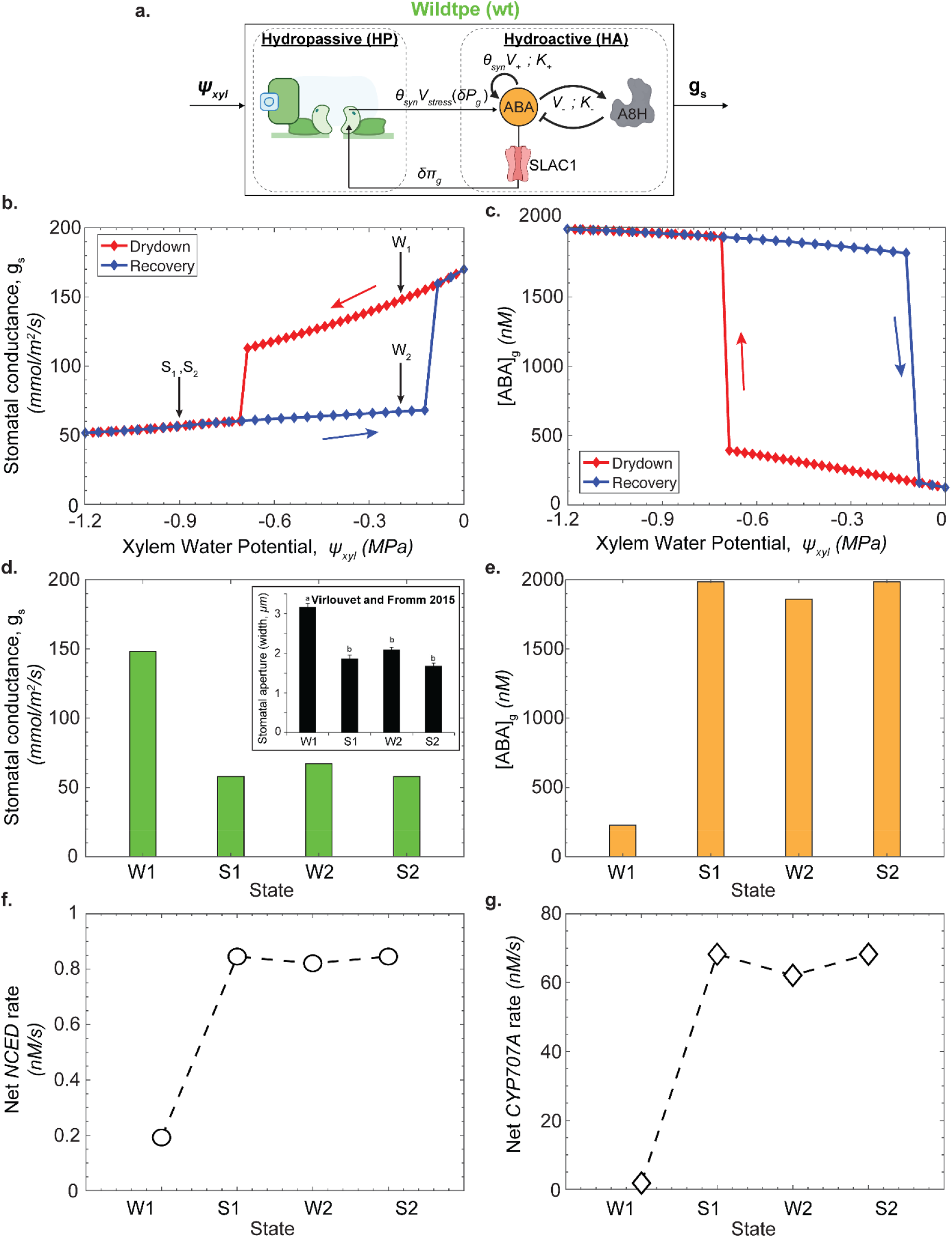
Bi-stability of ABA autoregulation can lead to hysteresis in steady state stomatal conductance through a cycle of drought. **a.** Schematic of the coupled model showing the hydropassive (HP) and hydroactive (HA) components as defined in Fig. 1. This case has stronger positive feedback in HA relative to that used in Fig. 2 and Fig. 3, representing stronger effect of ABA’s positive feedback synthesis loop. **b.-c**. Predicted hysteresis of steady state stomatal conductance, *g*_*s*_ (**b**) and concentrations of ABA in the guard cell, [*ABA*]_*g*_ (**c**). During drydown (decreasing xylem water potential, *ψ*_*xyl*_; red dots) the trajectory of *g*_*s*_ stays higher and the trajectory of [*ABA*]_*g*_ stays lower than their respective trajectories during recovery after rewetting (increasing xylem water potential, *ψ*_*xyl*_; blue dots). The sequential points along a simulated trajectory of steps of drydown and recovery are labeled as *W*_1_ → *S*_1_ → *W*_2_ → *S*_2_ . **d.-g**. Predicted stomatal conductance (**d**; **Inset:** experimental stomatal aperture hysteresis in *At* from Virlouvet and Fromm 2015^22^), guard cell ABA concentration, [*ABA*]_*g*_ (**e**), ABA synthesis rate via *NCED* (**f**), and A8H synthesis rate via *CYP707A* (**g**) over the sequence of states labeled in (**b**) through the following steps: first drying (*W*_1_ → *S*_1_); rewatering (*S*_1_ →*W*_2_); second drying (*W*_2_ → *S*_2_ ). Predictions were made for constant VPD = 1.1 kPa (RH=80% at 36°C). Parameters used were *K*_+_ = *K*_−_= 1560 *nM, V*_*stress*_ = 7.37 *nM*/*s, V*_+_ = 94 *nM*/*s, V*_−_ = 1.5 *nM*/*s* with *θ*_*syn*_ =1 for wildtype.

To simulate cycles of water availability, we varied xylem water potential (*ψ*_*xyl*_ ) keeping VPD constant at 1.1kPa and tracked the trajectories of steady state *g*_*s*_ (Fig. 4b) and [*ABA*]_*g*_ (Fig. 4c) through drydown (red symbols) and recovery (blue symbols). For drydown (decreasing *ψ*_*xyl*_), *g*_*s*_ remains high (Fig. 4b) and [*ABA*]_*g*_ remains low (Fig. 4c) from the unstressed (*ψ*_*xyl*_ *=*0 MPa) to a moderate stressed state (*ψ*_*xyl*_ ≅ −0.7 MPa) before transitioning to a low *g*_*s*_/high [*ABA*]_*g*_ state for continued lowering of *ψ*_*xyl*_; for recovery, the stomates remain in their low *g*_*s*_/high [*ABA*]_*g*_ state until near-complete recovery has been achieved (*ψ*_*xyl*_ ≅ −0.1 MPa), at which point the predicted state jumps back to the original high conductance state.

In Fig. 4d-g, we extract *g*_*s*_ (Fig. 4d; inset: experimental stomatal aperture hysteresis in *At* from Virlouvet and Fromm^22^), [*ABA*]_*g*_ (Fig. 4e), and the net rate of *NCED* (proxy for rate of ABA synthesis – Fig. 4f) and *CYP707A* (proxy for catabolic degradation of ABA – Fig. 4g) for four states visited as follows: from a well-watered state (W1) on a drydown to a stressed state (S1), followed by recovery to a well-watered state (W2) and a second drydown to a stressed state (S2). We note that while the water potentials for the two well-watered states were the same (*ψ*_*xyl*_ ≅ − 0.2), the predicted macroscopic state of the stomates (Fig. 4d) and underlying molecular states (Figs. 4e-g) were different for W1 and W2, reflecting differences in the two history-dependent solutions to the model equations.

Importantly, as we present in Fig. S16, these predictions of hysteretic behavior across scales – from ABA-related processes in guard cells (i.e., expression levels of *NCED* and *CYP707A*) to stomatal regulation agree semi-quantitively with molecular and macroscopic measurements reported by Virlouvet and Fromm^22^ in *At*. We note that the autoregulation of ABA in guard cells (Fig. 4a) drives this hysteresis in our model, as illustrated by the analysis of the equations governing autoregulation in Fig. S6. Although previous investigators have suggested a role for ABA in generating hysteresis^22,23^, no predictive models of this process have been put forward. More generally, we are unaware of models of stomatal response that capture this potentially critical form of stress memory and adaptation to long term variations in drought stress.

## Discussion

Our coupled hydropassive–hydroactive (HP–HA) model (Fig. 1e-f) builds on 50 years of developments of hydraulic models^7,9–13^ and, for the first time, introduces a framework to integrate the richness of genetic^40,43,46^, molecular^44,45,50^, and membrane^41,48,49^ processes by which ABA could modulate stomatal conductance. With this model, we identify guard cell-localized autoregulation of abscisic acid (ABA) as a central driver of HA stomatal responses including the emergent phenomena of damped and sustained oscillations and of hysteresis during drought and recovery. This localization provides a unified physical basis for responses to changes in water availability and demand without invoking elusive humidity sensors^26,69^, explaining both the rapid response to water demand (VPD), and the role of hysteresis as a functional “stress memory” that anticipates recurrent deficits in water availability (*ψ*_*xyl*_). While not relevant to the existing data in *At* used here, we note that our formulation of HP processes allows for accounting for newly elucidated, non-stomatal regulation of transpiration^29–31^ by explicitly modeling the evaporative processes within the leaf (Fig. 1d).

This framework provides new avenues to pursue important outstanding questions about stomatal responses to water stress and demand by allowing for the encoding of genotype-specific molecular and tissue-scale hypotheses. For example, while we show that mechanobiological coupling of guard cell turgor to guard cell-localized upregulation of ABA (*V*_*stress*_ (*δP*_*g*_); Fig. 1e) can explain observed stomatal dynamics in *At* (Fig. 2b) in a manner consistent with existing measurements of foliar ABA in other species^38,39^ and regulation of ABA synthesis^40,43^, the model could allow for the falsification of specific hypotheses (e.g., localized mechanotransduction and ABA synthesis) with the availability of new data^70^. Particularly valuable data would include simultaneous, real-time measurements of cytosolic ABA concentration (via FRET-based reporters^62^) and guard cell turgor (via pressure probe^71,72^); and spatially resolved flux measurements of ABA to deconstruct the *V*_*stress*_ term using guard-cell specific promoters driving reporters for key genes (e.g., *NCED* and *CYP707A*) to quantify their relative *in vivo* contributions during a stress response.

With our focus on water stress responses of individual stomates, our model complements and should be integrated with the comprehensive treatments of ion transport and metabolic networks for the response to light and CO_2_ as in the “OnGuard” models^14–16^ and with hypotheses of stomate-stomate coupling such as proposed by Mott and Peak^36^. Such integration would yield a unified, multi-signal systems model capable of quantifying, for example, how ABA-driven stomatal closure competes with light-driven opening, how CO_2_ modulates the ABA pathway, and how fully detailed, single-cell processes can couple to define whole-leaf spatial and temporal patterning of stomatal dynamics to address collective dynamics like stomatal patchiness. Further integration with models of photosynthesis^73^ would form an unprecedented basis to pursue criteria for optimal stomatal function, providing a mechanistic foundation for the long-standing ideas about evolutionary optima^24,74^ and for the bioengineering of programmable plants that allow for resource-use efficient crop production^75,76^. Finally, scaling this molecular-to-tissue framework to canopy processes^77^ would allow for mechanistic prediction of ecosystem-level plant-atmosphere coupling that is strongly controlled by stomatal conductance^26,78,79^.

## Methods

This section provides a concise overview of the hydropassive (HP) and hydroactive (HA) model formulations. Detailed derivations for both models, a complete list of parameters, and the solution approach are presented in the supplementary information (sections S2-S4 and S7).

### Hydropassive model

The HP model describes the regulation of stomatal aperture via changes in guard cell volume and turgor driven primarily by hydraulic signals. Stomatal conductance is determined by guard cell turgor pressure and the backpressure from subsidiary cells^71,72^. At the leaf scale, passive water movement through the xylem, mesophyll, subsidiary cells, and guard cells affects guard cell turgor pressure, which in turn influences stomatal conductance. These physical processes in the HP model are represented using an electrical analogy of hydraulic capacitances, resistances and water fluxes driven by differences in water potential^7,9–13^. The model formulation is provided in equations S1–S39 and parametrized using values in Tables S4–S5.

### Hydroactive model

The HA model incorporates stress-triggered, guard cell-localized ABA-mediated regulation of stomatal aperture^46^. This regulation is controlled by the movement of ions across the guard cell membrane, which alters the guard cell osmotic potential. Changes in osmotic potential then modify cell’s total water potential, guard cell turgor, and ultimately stomatal conductance (Fig 1f.).

The model incorporates findings that stomatal response to reduced humidity is driven by *de novo* ABA synthesis via *NCED* expression^38–40,46,56,57,62^. It has been shown that guard cells in *At* produce ABA under reduced humidity, expressing genes required for ABA biosynthesis driven by *NCED* expression^46,62^. This synthesis pathway is triggered by decreasing leaf turgor^38–40^. We model this stress-dependent ABA synthesis rate as *V*_*stress*_ (*δP*_*g*_). The term *V*_*stress*_ [*nM/s*] represents the “net” increase in local guard cell ABA levels due to water stress and can subsume the rate of increase from other sources of ABA such as release from its conjugated form, or transported ABA from tissues distal to guard cells.

Increased ABA binds to its PYR receptor, inhibiting the phosphatase PP2C. PP2C regulates futile cycles involving OST1 and the anion channel SLAC1^44,45,50,58–61^. ABA-mediated activation of SLAC1 facilitates chloride ion efflux^48,49^, leading to plasma membrane depolarization and subsequent potassium ion efflux^41,50^. This loss of solutes alters guard cell osmotic potential, reduces turgor, and decreases stomatal conductance. The HA model formulation is provided in equation S40–S102 and parameterized using values in Tables S6–S10.

Key features of this model include: (i) an empirical relation between ABA synthesis rate and guard cell hydraulic status^70,80^ (red sigmoidal curve in Fig. 1e), derived from the measurements of McAdam^38,39^ (SI Eq. S103, section S4.A; Fig. S5); mechanistic details of this critical mechanobiological coupling remain unknown^40,51^; (ii) coarse-grained biokinetic rate equations describing how ABA regulates its own synthesis and degradation via the 8′-hydroxylase (A8H) cytochrome P450, informed by *in vitro* data^43,46,53–57^ (SI Eqs. S49–S56, section S3.B.4); (iii) ABA-dependent signaling through the PYR receptor and coupled futile phosphorylation cycles controlling the SLAC1 anion channel, following *in vitro* characterizations^48,49,58,59,61^ (SI Eqs. S57–S80, section S3.B.4); and (iv) a Goldman–Hodgkin–Katz-type description of osmoregulation mediated by major ion channels (SLAC1, GORK, KAT1), 2H/Cl symports, and H^+^-ATPase pumps, parameterized using electrophysiological and biochemical data^14–16^ (SI Eqs. S81– S102, section S3.C). We provide details on the assumptions, governing equations, and parameters of this HA model in SI section S3.C.1 and Table S6-S10.

### Coupling

The HP and HA models are coupled at two key points. (i) Linking Water Status to ABA Biosynthesis: Changes in guard cell turgor pressure, driven by water status in the hydropassive model, trigger ABA biosynthesis *V*_*stress*_ (*δP*_*g*_). This represents water stress stimuli from the hydropassive model informing biochemical process in the hydroactive model. (ii) Feedback on Turgor Pressure: Changes in guard cell osmotic potential, governed by the hydroactive model, feedback to regulate turgor pressure (*δ π*_*g*_). Changes in the guard cell osmotic potential then modify cell’s total water potential, and guard cell turgor. This physiological osmotic adjustment thus affects stomatal conductance, completing the feedback loop between the two models.

### Model Parametrization

The HP model was parameterized to reproduce the stomatal response of the ABA-insensitive *ost1* mutant^17^; these parameters were then held constant for all genotypes. The HA model required additional parameters for guard cell electrophysiology, ABA signaling, and ABA regulation. Electrophysiology parameters were sourced from literature on *At*, supplemented with data from *Vicia faba* where necessary. ABA signaling rate parameters were taken directly from the literature.

Six unknown parameters for the ABA regulation motif (*V*_*stress*_, *V*_+_, *V* _−_,,*K*_+_ *K*_−_, *θ*_*syn*_) were calibrated by matching model predictions to six experimental measurements from Merilo^17^, creating a fully constrained system. Using a non-dimensionalized formulation of the ABA equations (See SI section S5.B), we isolated the relative timescales of ABA and hydraulic dynamics. We employed a sequential fitting strategy: wildtype parameters were determined by fixing *θ*_*syn*_ *=* 1, constraining characteristic concentration ratios (.*λ*_−_, . *λ*_+_), and fitting both transient and steady-state responses using the dimensionless groups (Ω_1_, Ω_2_, Ω_3_). Parameter values were selected from regions that reproduced the experimentally observed damped oscillations. The ABA synthesis mutant was then parameterized by varying *θ*_*syn*_ while keeping other parameters fixed. Full details of the non-dimensionalization, fitting procedure, and bifurcation analysis are provided in SI sections S5. Comprehensive tables listing all model parameters, their sources, and the methodology for parameter fitting are provided in SI section S7.

## Code Availability

All MATLAB code used to generate the results in this study is permanently archived in a Zenodo repository at: https://doi.org/10.5281/zenodo.17888362. The most current version of the code is also available on GitHub at: https://github.com/desai-sahil/sys-bio-gs.git. Refer to SI section S7.C for details on steps to run the code to reproduce the results in main text.

## Acknowledgements

We thank F. E. Rockwell, V. Bacheva, S. Sen, I. Gabay, E. Wu, J. Belding, and P. Jain for insightful discussions. This work was supported by the Center for Research on Programmable Plant Systems (CROPPS), a Science and Technology Center of the National Science Foundation under Grant No. DBI-2019674.

