## Supplemental Information for "Abscisic acid-mediated water stress regulation can mechanistically explain oscillations and water stress memory in stomatal conductance"

### This PDF file includes:

**S8 References** **51**

### Supporting Information Text

### S1. Review and assessment of existing models of stomatal regulation

Following the tradition of Cowan[1] and Delwiche[2], Buckley 2003[3] presented an isothermal mechanistic model for predicting stomatal conductance by integrating both hydromechanical and biochemical processes. The authors developed a closed-form model that combined water potential gradients arising from transpiration with hydromechanical interactions among guard cells, epidermal cells, and the surrounding tissues, while also incorporating biochemical regulation derived from Farquhar's photosynthesis-based model[4] applied to guard cells, enabling the simulation of stomatal responses to light and carbon dioxide.

Similar to the anatomical arrangement used by Delwiche[2], the Buckley model[3] adopted a more physiologically realistic architecture relative to that originally proposed by Cowan[1] in which guard cells were positioned in series with subsidiary cells. Physiological parameters were informed by pressure-probe experiments, strengthening the model's empirical grounding. Transpiration rate was formulated as a function of stomatal conductance and the vapor pressure gradient between the saturated internal airspace and the atmosphere, with feedback effects mediated by changes in water potentials of the guard and epidermal cells.

A central assumption of the Buckley model[3] was that changes in guard cell osmotic potential were directly proportional to subsidiary cell turgor. This simplification served as a proxy for the osmo-mechanical coupling regulated by solute transport across the guard cell membrane. While this assumption enabled a functional link between steady-state stomatal conductance and environmental drivers, it lacked mechanistic detail regarding the underlying osmoregulatory processes and did not capture rate-limiting biochemical steps. As a result, the model was unable to generalize across genetic variants and lacked predictive power for mutants with altered molecular regulation. With this model, Buckley argued that rectification in stomatal behavior requires hydroactive regulation of guard cell osmotic potential, emphasizing the importance of active solute transport mechanisms in determining stomatal responses.

Buckley 2006[5] and Buckley 2011[6] extended this framework by building a transient model upon his earlier formulation. The updated model incorporated a bundle sheath extension compartment and capacitance for epidermal compartment, allowing it to predict how bundle sheath extensions influence the transients in stomata in response to humidity shifts and leaf excision. Buckley's model predicts the WWR as a consequence of hydromechanical feedback: a rapid drop in subsidiary cell turgor increases the mechanical advantage over the guard cells, transiently opening the stomata before biochemical regulation reasserts control and restores closure. Similarly, earlier hydromechanical models by Cowan[1] and Delwiche[2] could reproduce the WWR through purely mechanical interactions between guard and subsidiary cells without invoking biochemical feedback.

Building on the hydropassive foundation established by Buckley[3,5,6], we followed his treatment of the following: (i) hydraulic circuit representation of tissue compartments, resistances, capacitances, and osmotic potentials (Fig. S1) and (ii) the parameter values that defined these hydraulic elements (Table S5). Relative to Buckley's approach, we modified the formulation of evaportive loss. We implement the following distinct approaches relative to Buckley's formulation: (i) we evaluate the evaporative processes from mesophyll ($E_m$), subsidiary ($E_s^1$), and guard ($E_g^1$) cells into the intercellular airspaces and from the stomatal pore ($E_{st}$) and subsidiary ($E_s^2$) and guard ($E_g^2$) cells into the external atmosphere (Fig. S1) explicitly based on local differences in water potential between the tissue and air. This formulation provides more realistic coupling of the hydraulic compartments to the vapor phase. It also allows for the treatment of undersaturation in the substomatal cavity ($h_{ssc} < 1$), although we do not explore such cases in this study; (ii) We replace the phenomenological assumption used in earlier models for guard cell osmotic potential with a detailed hydroactive (HA) model. By adding explicit kinetics for ABA autoregulation and ion transport, our coupled HP-HA framework (Section S4) moves beyond the passive 'wrong-way' responses predicted by hydraulic mechanics alone. This integration allows the model to mechanistically predict complex emergent behaviors, such as sustained oscillations and hysteresis, and to recapitulate the distinct stomatal dynamics of specific genetic mutants.

While we are unaware of published mathematical models of the ABA autoregulation and signaling treated here, our treatment of these processes (Fig. S2 and Fig. S3) is guided by the systems biology approaches to model related gene-regulatory and signaling processes[7–9]. For our treatment of the membrane processes that define osmoregulation in the guard cells, we were informed by the work of Blatt and colleagues in the development of OnGuard[10–12]. Our coarse grained representation of these processes draws on well-established approaches in electrophysiology[10,13,14].

### S2. Hydropassive hydraulic model

The hydraulic model accounts for movement of liquid water in the outside xylem zone (OXZ) from xylem to bundle sheath, bundle sheath to mesophyll, and from mesophyll to epidermis – first to subsidiary cells and then to the guard cells (Fig. S1). We subsume the bundle sheath into the mesophyll and the rest of the epidermis into the subsidiary cells here. In this model we consider transpiration from mesophyll, subsidiary and guard cells. The perturbations to the model are introduced through a change in xylem water potential ($\psi_{xyl}$) or a change in atmospheric relative humidity ($h_a$) through changes in vapor pressure deficit ($VPD$).

Hydraulic responses involve stomatal aperture modulation driven by changes in guard cell turgor. In angiosperms, stomatal conductance is considered to be primarily regulated by deformations in the guard and subsidiary cells driven by changes in their states of turgor[15–17]. Hence, it is important to represent all the components of the OXZ that have the potential to affect the state of turgor in subsidiary and guard cells. In this model, outside xylem tissue is represented consisting of the guard cells, subsidiary cells and spongy mesophyll cells using an electrical circuit analog (Fig. S1). Hydraulic resistors (zig-zag lines), capacitors (parallel lines of equal length) , and potential sources representing osmotic potential (parallel lines of unequal length) are the primary components, and water flux is the analogous current flow driven by water potential gradient analog of voltage.

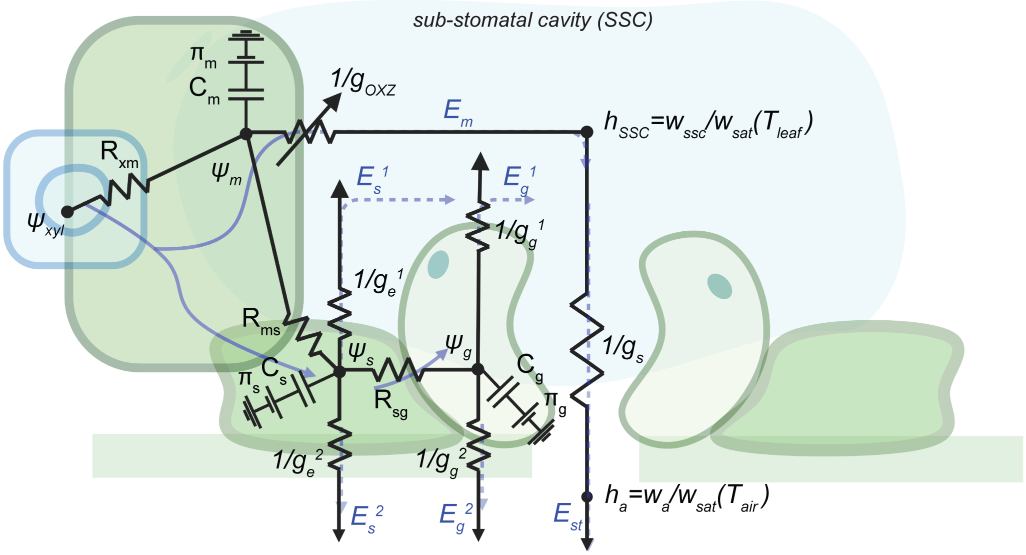

**Fig. S1. Resistive-capacitive (RC) representation of the hydropassive hydraulic circuit**. Schematic representation of the hydraulic circuit used to model hydropassive responses, superimposed on a sketch of the leaf cross section from Fig. 1c. Following Cowan's[1] notation, the circuit contains resistances ($R$ $[MPa.m^2.s/mmol]$ - zig-zag lines), capacitances ($C$ $[mmol/MPa.m^2]$ - a pair of parallel lines of equal length), osmotic potential sources ($\pi[MPa]$ - a pair of parallel lines of unequal length), and grounds (a triplet of parallel lines of different length) across the xylem (x), mesophyll (m), subsidiary (s), guard (g) cells, and sub-stomatal cavity (SSC). The hydraulic charge on the capacitors is the turgor pressure, $P$ $[MPa]$. Water potential $\psi$ $[MPa]$ is the voltage analog and the water flux $E$ $[mmol/m^2.s]$ is the current analog of an electrical circuit. Fluxes and conductances are defined per unit area of leaf. The hydraulics of bundle sheath cells are included in the mesophyll; the hydraulics of other epidermal cells are included in the subsidiary cells. The water fluxes pass as a liquid (blue curve) from the xylem to the mesophyll, subsidiary, guard cells, and to sites of evaporation. Vapor fluxes (dashed blue curves)) pass into the SSC (relative humidity, $h_{SSC}$ from the mesophyll ($E_m$), subsidiary ($E_s^1$), and guard ($E_g^1$) cells and contribute to the total flux ($E_{st}$) limited by the stomatal resistance ($1/g_s$ $[m^2.s/mmol]$); vapor flux also leaves the subsidiary ($E_s^2$) and guard ($E_g^2$) cells directly into the atmosphere at the leaf surface (relative humidity. $h_a$). The governing equations for this circuit are provided in Section S2 (Eqs. S37-S39).

**A. Water relations coupling xylem to guard cells.** For each of the hydraulic compartments – mesophyll, subsidiary, and guard cell (Fig. S1) – we track total symplastic water potential ($\psi_i$ [MPa]), turgor pressure ($P_i$ [MPa]), and osmotic potential ($\pi_i$ [MPa]). Here the subscript $'i'$ indicates that the parameter can refer to the mesophyll (i = m), subsidiary cell (i = s), or guard cell (i = g). The total water potential can be expressed as:

$$\psi_i = P_i - \pi_i \quad \text{[S1]}$$

Small changes in cellular pressure can be related to volume changes due to water moving in and out of the cells. For elastic cell walls, a simple linear relationship is used as in Delwiche[2]:

$$\delta P_i = \varepsilon_i \Big(\frac{\delta V_i}{V_i^0}\Big) \quad \text{[S2]}$$

$$P_i - P_i^0 = \varepsilon_i \Big(\frac{V_i - V_i^0}{V_i^0}\Big) \quad \text{[S3]}$$

The parameters in Eq. S3 are defined as follows and values are provided in Table S5.

- $\varepsilon_i$ [MPa]: cell wall elasticity
- $P_i^0$ [MPa]: cell reference turgor pressure
- $P_i$ [MPa]: instantaneous cell turgor pressure
- $V_i^0$ [mmol/m$^2$] is the cell reference volume per leaf area
- $V_i$ [mmol/m$^2$] is the instantaneous cell volume per leaf area

The osmotic potential of the cells is represented as:

$$\pi_i = \frac{n_p RT}{V_i} \quad \text{[S4]}$$

The parameters in Eq. S4 are defined as follows and values are provided in Table S5.

- $n_p$ [mol]: moles of solute in the cell
- $R$ [J/mol/K]: gas constant
- $T$ [K]: temperature of leaf

Changes in osmotic potential are due to changes in cell volume and changes in moles of solute in the cell:

$$\delta \pi_i = -\frac{n_p RT}{V_i^0}\Big(\frac{\delta V_i}{V_i^0}\Big) + \delta n_p \frac{RT}{V_i^0} \quad \text{[S5]}$$

$$\delta \pi_i = -\pi_i^0 \Big(\frac{\delta V_i}{V_i^0}\Big) + \delta \bar{\pi}_i \quad \text{[S6]}$$

Where, $\pi_i^0$ [MPa] $= n_p RT/V_i^0$ is the reference osmotic potential of the cell at $P_i^0$ and $V_i^0$ and $\delta\bar{\pi}_i$ is the active biochemically induced change in osmotic potential. The active biochemically induced change in osmotic potential is only considered for guard cells ($\delta\bar{\pi}_g$), as described in Section S3 (Eq. S99).

Change in total water potential of the cell can be expressed as:

$$\delta \psi_i = \delta P_i - \delta \pi_i \quad \text{[S7]}$$

Using equations (S2) and (S6)

$$\delta \psi_i = \delta P_i + \pi_i^0 \frac{\delta V_i}{V_i^0} - \delta \bar{\pi}_i \quad \text{[S8]}$$

$$\delta\psi_i = \delta P_i\left(1 + \frac{\pi_i^0}{\varepsilon_i}\right) - \delta\bar{\pi}_i \quad \text{[S9]}$$

$$\psi_i - \psi_i^0 = (P_i - P_i^0)\left(1 + \frac{\pi_i^0}{\varepsilon_i}\right) - \delta\bar{\pi}_i \quad \text{[S10]}$$

Where, $\psi_i^0$ [MPa] is the reference total water potential at reference $\pi_i^0$ and $P_i^0$.

Stomatal conductance is proportional to the aperture of stomata which can be expresses as a linear function of guard cell turgor, subsidiary cell turgor and mechanical advantage[3,5]:

$$g_s = \chi(P_g - m * P_s) \quad \text{[S11]}$$

The parameters in Eq. S11 are defined as follows and parameter values are provided in Table S4.

- $g_s$ [mmol/m$^2$/s]: stomatal conductance per leaf area basis
- $\chi$ [mmol/m$^2$/s/MPa]: proportionality constant linking turgor to stomatal conductance
- $m$ [$-$]: mechanical advantage of subsidiary cells over guard cells
- $P_g$ [MPa]: turgor pressure of guard cells
- $P_s$ [MPa]: turgor pressure of subsidiary cells

In Fig. S1 we represented the vapor pressure $w_i$ in terms of relative humidity $h_i = w_i/w_{sat}$ where $w_{sat}$ [MPa] is the saturation vapor pressure at temperature $T$ [K]. Water potential and vapor pressure are related by Kelvin equation:

$$\psi_i = \frac{RT}{\bar{\nu}_w} log\left(\frac{w_i}{w_{sat}}\right) \quad \text{[S12]}$$

Where, $\bar{\nu}_w$ [m$^3$/mol] is the molar concentration of water at temperature $T$.

Rewriting Kelvin equation for vapor pressure:

$$w_i = w_{sat} * \exp\left(\frac{\psi_i}{RT/\bar{\nu}_w}\right) \quad \text{[S13]}$$

Linearizing Kelvin equation for small deviations from saturation:

$$w_i \approx w_{sat} * \left(1 + \frac{\psi_i}{RT/\bar{\nu}_w}\right) \quad \text{[S14]}$$

The value for the conductance to transpiration from the mesophyll to the sub-stomatal cavity was sourced from recent studies on maize[18] and tomato[19]. These values were originally determined using the AquaDust[20].

$$k_{oxz} = 0.36 + \frac{2.59}{1 + exp(-15.61 * (\psi_{xyl} + 0.76))} \quad \text{[S15]}$$

Where $k_{oxz}$ [mmol/m$^2$/s/MPa] represents the liquid conductance of the outside xylem zone to water. In this study, we set $k_{oxz}$ at its maximum value ($k_{oxz} = 2.95\ mmol/m^2/s/MPa$) to avoid confounding non-stomatal effects with stomatal changes.

$$g_{oxz} = \frac{RT}{\bar{\nu}_w} k_{oxz} \quad \text{[S16]}$$

The hydraulic conductance to vapor transpiration from mesophyll, $g_{oxz}$[mmol/m$^2$/s] is inferred from $k_{oxz}$.

### B. Definition of fluxes.

***B.1. Inter-compartment fluxes.*** In Eqs. S17-S19, the form of the flux equations implies transmembrane transport for which the difference in total water potential is the relevant driving force for flow based on an assumption that the plasma membranes present perfect semi-permeable properties (reflection coefficient of 1 for all solute). In the mesophyll and subsidiary compartments, these equations also neglect, for simplicity, gradients of potential within these multi-cellular structures. For $J_1$ in Eq. S17, the flux from xylem (apoplast) to mesophyll (symplast) must include a transmembrane step. The form in Eq. S17 attributes the limiting resistance into and across the mesophyll to this step. While purely symplastic (i.e., via plasmadesmata) paths may exist between the mesophyll and the epidermis (included within the subsidiary compartment), in the absence of active adjustment of osmotic potentials in these compartments, differences in total water potential can serve approximately as the driving force for both the symplastic and trans-cellular fluxes that could contribute to $J_2$ in Eq. S18. For flux, $J_3$ in Eq. S19 from the subsidiary compartment to the guard cells, we expect trans-cellular flux because mature stomata lack plasmodesmatal connections between guard cells and subsidiary cells[21].

$$J_1 = \frac{\psi_{xyl} - \psi_m}{R_{xm}} \quad [S17]$$

$$J_2 = \frac{\psi_m - \psi_s}{R_{ms}} \quad [S18]$$

$$J_3 = \frac{\psi_s - \psi_g}{R_{sg}} \quad [S19]$$

The parameters in Eqs. S18-S19 are defined as follows and parameter values are provided in Table S5.

- $J_1$ [mmol/m$^2$/s]: flux per leaf area basis from xylem to mesophyll (positive in that direction)
- $J_2$ [mmol/m$^2$/s]: flux per leaf area basis from mesophyll to subsidiary (positive in that direction)
- $J_3$ [mmol/m$^2$/s]: flux per leaf area basis from subsidiary to guard (positive in that direction)
- $R_{xm}$ [m$^2$.s.MPa/mmol]: hydraulic resistance per leaf area basis between xylem and mesophyll
- $R_{ms}$ [m$^2$.s.MPa/mmol]: hydraulic resistance per leaf area basis between mesophyll and subsidiary
- $R_{sg}$ [m$^2$.s.MPa/mmol]: hydraulic resistance per leaf area basis between subsidiary and guard
- $\psi_m$ [MPa]: mesophyll symplast total water potential
- $\psi_s$ [MPa]: subsidiary symplast total water potential
- $\psi_g$ [MPa]: guard symplast total water potential

***B.2. Storage fluxes.*** The storage flux is the flux that causes a change in the volume of cells written as:

$$J_m = \frac{dV_m}{dt} \quad [S20]$$

$$J_s = \frac{dV_s}{dt} \quad [S21]$$

$$J_g = \frac{dV_g}{dt} \quad [S22]$$

The parameters in Eqs. S24-S26 are defined as follows:

- $J_m$ [mmol/m$^2$/s]: flux per leaf area basis going into mesophyll (positive in that direction)
- $J_s$ [mmol/m$^2$/s]: flux per leaf area basis going into subsidiary (positive in that direction)
- $J_g$ [mmol/m$^2$/s]: flux per leaf area basis going into guard (positive in that direction)

To express the flux in terms of rate of change of water potential, we use capacitance, $C_i$ [mmol/m$^2$/MPa] of the cell which is a measure of change in volume due to a change in cell water potential.

$$C_i = \frac{\partial V_i}{\partial \psi_i} = \frac{V_i^0}{\varepsilon_i + \pi_i^0} \quad \text{[S23]}$$

Using Eq. S23, Eqs. S24 - S26 can be written as:

$$J_m = \frac{dV_m}{dt} = C_m \frac{d\psi_m}{dt} \quad \text{[S24]}$$

$$J_s = \frac{dV_s}{dt} = C_s \frac{d\psi_s}{dt} \quad \text{[S25]}$$

$$J_g = \frac{dV_g}{dt} = C_g \frac{d\psi_g}{dt} \quad \text{[S26]}$$

***B.3. Transpiration fluxes.*** The transpiration flux that is lost from each cell is expressed as a product of the vapor pressure driving force and corresponding hydraulic vapor conductance. Liquid water changes its phase into vapor before leaving the cell. For mesophyll cells transpiring into the sub-stomatal cavity, the driving force is the difference in partial vapor pressure of liquid water in mesophyll cell and the partial vapor pressure of the air in the sub-stomatal cavity. The guard and subsidiary cells can transpire into the sub-stomatal cavity or directly into the atmosphere through a cuticle.

$$E_m = g_{oxz}\Big(\frac{w_m - w_{ssc}}{P_{atm}}\Big) \quad \text{[S27]}$$

$$E_s^1 = g_e^1\Big(\frac{w_s - w_{ssc}}{P_{atm}}\Big) \quad \text{[S28]}$$

$$E_s^2 = g_e^2\Big(\frac{w_s - w_a}{P_{atm}}\Big) \quad \text{[S29]}$$

$$E_g^1 = g_g^1\Big(\frac{w_g - w_{ssc}}{P_{atm}}\Big) \quad \text{[S30]}$$

$$E_g^2 = g_g^2\Big(\frac{w_g - w_a}{P_{atm}}\Big) \quad \text{[S31]}$$

$$E_{st} = g_s\Big(\frac{w_{ssc} - w_a}{P_{atm}}\Big) \quad \text{[S32]}$$

The parameters in Eqs. S27-S32 are defined as follows and parameter values are provided in Table S5.

- $E_m$ [mmol/m$^2$/s]: flux per leaf area basis going from mesophyll to sub-stomatal cavity
- $E_s^1$ [mmol/m$^2$/s]: flux per leaf area basis going from subsidiary to sub-stomatal cavity
- $E_s^2$ [mmol/m$^2$/s]: flux per leaf area basis going from subsidiary to atmosphere
- $E_g^1$ [mmol/m$^2$/s]: flux per leaf area basis going from guard to sub-stomatal cavity

- $E_g^2$ [mmol/m$^2$/s]: flux per leaf area basis going from guard to atmosphere
- $E_{st}$ [mmol/m$^2$/s]: flux per leaf area basis leaving the leaf internal air-space through stomata
- $g_m^{H_2O}$ [mmol/m$^2$/s]: conductance to transpiration from mesophyll into sub-stomatal cavity
- $g_e^1$ [mmol/m$^2$/s]: conductance to transpiration from subsidiary into sub-stomatal cavity
- $g_e^2$ [mmol/m$^2$/s]: cuticular conductance to transpiration from subsidiary into atmosphere
- $g_g^1$ [mmol/m$^2$/s]: conductance to transpiration from guard into sub-stomatal cavity
- $g_g^2$ [mmol/m$^2$/s]: cuticular conductance to transpiration from guard into atmosphere
- $w_{ssc}$ [MPa]: vapor pressure of air-space in OXZ
- $w_m$ [MPa]: vapor pressure of water in mesophyll cell
- $w_s$ [MPa]: vapor pressure of water in subsidiary cell
- $w_g$ [MPa]: vapor pressure of water in guard cell
- $w_a$ [MPa]: vapor pressure of water in atmospheric air
- $P_{atm}$ [MPa]: total atmospheric pressure

**C. Flux balance.** Applying flux balance in the hydraulic circuit at cell nodes:

At the mesophyll node ($\psi_m$):

$$J_1 = J_m + J_2 + E_m \tag{S33}$$

At the subsidiary node ($\psi_s$):

$$J_2 = J_s + J_3 + E_s^1 + E_s^2 \tag{S34}$$

At the guard node ($\psi_g$):

$$J_1 = J_g + E_g^1 + E_g^2 \tag{S35}$$

At the internal leaf air-space node ($w_{ssc}$):

$$E_{st} = E_m + E_s^1 + E_g^1 \tag{S36}$$

**D. Governing equations.** Using Eqs. S24 - S26 and Eqs. S33 - S35 (RHS with Eqs. S17-S19) we write the system of ordinary differential equations (ODEs) for total potential change in mesophyll, subsidiary, and guard cells:

$$\frac{dV_m}{dt} = C_m \frac{d\psi_m}{dt} = \frac{\psi_{xyl} - \psi_m}{R_{xm}} - \frac{\psi_m - \psi_s}{R_{ms}} - E_m \tag{S37}$$

$$\frac{dV_s}{dt} = C_s \frac{d\psi_s}{dt} = \frac{\psi_m - \psi_s}{R_{ms}} - \frac{\psi_s - \psi_g}{R_{sg}} - E_s^1 - E_s^2 \tag{S38}$$

$$\frac{dV_g}{dt} = C_g \frac{d\psi_g}{dt} = \frac{\psi_s - \psi_g}{R_{sg}} - E_g^1 - E_g^2 \tag{S39}$$

We present the coupling of these processes to the hydroactive ones in Section S4. In Section S7, we present the parameter values (Tables S4, S5) and the method of solution (Section S7C).

### S3. Hydroactive biochemistry model

In this study we highlight the key family of genes and proteins that are known to be involved in the core Abscisic Acid (ABA) dependent synthesis and signaling pathway. As illustrated in Fig. S2 and describe below, we pursue a coarse-grained representation of the multiple genes involved in the synthesis and degradation of ABA, replacing each step with a single representative gene (NCED for synthesis of ABA and CYP707A for synthesis of A8H) (Fig. S2a). This type of reduced model has proven useful in a variety of applications and allows us to map our hypothetical structure on to well-studied motifs, as in Fig. S2b for the combined positive-negative feedback loop motif[8,9,22].

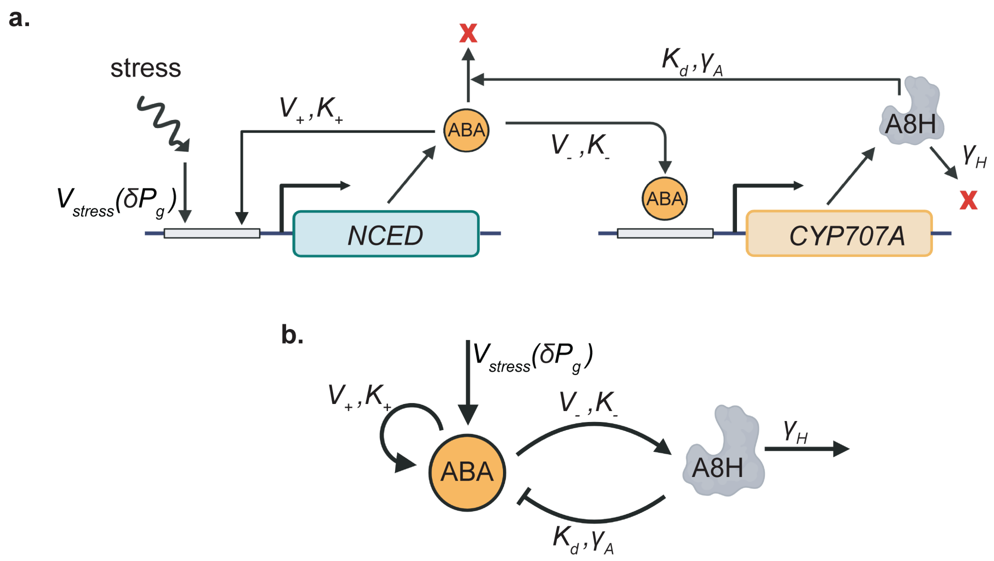

**Fig. S2. ABA regulation motif. a.** Schematic diagram of Abscisic Acid (ABA)-mediated pathway for stomatal closure. Changes in turgor pressure in guard cells ($\delta P_g$ $[MPa]$) upregulates stress-induced *de novo* biosynthesis of ABA (rate-limiting *NCED* shown here), characterized by the stress-regulated kinetic rate $V_{stress}$ $(\delta P_g)$ $[nM/s]$. ABA upregulates its own production ($V_+$ $[nM/s]$, $K_+$ $[nM]$), and its own degradation via upregulation of *CYP707A* gene family that encodes the enzyme 8'-hydroxylase, A8H, governed by Michaelis-Menten kinetics ($V_-$ $[nM/s]$, $K_-$ $[nM]$). A8H catalyzes ABA catabolism following Michaelis-Menten kinetics ($K_d$ $[nM]$, $\gamma_A$ $[1/s]$). Degradation of A8H is treated as first order ($\gamma_H\,[1/s]$). **b.** Shorthand representation of ABA regulation motif in (**a**), as shown in Fig. 1d. Positive regulation is represented by pointed arrows, while inhibitory interactions are indicated by flat arrows. The governing equations for this circuit are provided in Section S3A (Eqs. S53 and S56).

.

**A. Abscisic Acid regulation motif.** The *de novo* biosynthesis of ABA is a multi-step process that starts with carotenoids in chloroplast and ends with the formation of ABA in the cytoplasm[23]. In guard cells, ABA synthesis results from upregulation of ABA biosynthetic genes[24,25], with a strong association to the rapid activation of 9-cis-epoxycarotenoid dioxygenase ($NCED$) genes. The NCED gene family encodes the rate-limiting enzyme in the ABA biosynthesis pathway, with $NCED3$ and $NCED5$ playing a pivotal regulatory role. In multi-step reactions, the rate-limiting step dominates the overall reaction rate, acting as a kinetic bottleneck. For this reason, we let regulation of a single, effective gene "$NCED$" represent the full multi-gene process (Fig. S2a). We account for two modes of induction of this NCED pathway: the first is in response to changes in the state of turgor of the guard cells ($V_{stress}(\delta P_g)$). This hypothetical response to stress is motivated by the observation that $NCED3$ is rapidly induced by high $VPD$[26] and by the observation that ABA levels in leaf tissues increases with decreasing water potential[27,28]. The second mode of induction is in response to the level of ABA as a positive feedback ($V_+$ and $K_+$). This hypothesis of positive feedback is motivated by reports of such positive autoregulation by ABA on multiple genes in its biosynthetic pathway[29,30]. Reaction (R1) accounts for both these modes of induction. $V_+$ [nM/s] is the maximal synthesis rate of ABA from auto-upregulation of biosynthesis genes, $K_+$ [nM] is the half-maximum concentration of ABA for maximal auto-upregulation, and $n$ is the number of genes that are upregulated by ABA.

$$\xrightarrow[K_+,\ n]{V_{stress}(\delta P_g),\ V_+} \mathrm{ABA} \qquad \text{[R1]}$$

Abscisic acid also regulates its own degradation via upregulation of *CYP707A* genes that encode for an enzyme 8'-hydroxylase (A8H, written as H)[23,31]. Specifically, $CYP707A1$ and $CYP707A3$ genes are upregulated in an ABA-dependent manner under low VPD[26,31,32]. H is involved in enzymatic degradation of ABA into Phaseic Acid (PA)[23,31,33]. Understanding the principles governing biological network dynamics requires considering both synthesis and degradation rates[34]. Since degradation rates often control time-scales of physiological responses, they are critical to understanding the temporal dynamics of ABA regulation. Reaction (R2) accounts for ABA dependent upregulation of H, where $V_-$ [nM/s] is the maximal ABA dependent upregulation rate of H, and $K_-$ [nM] is the half-maximum concentration of ABA for H upregulation. Reaction (R3) is the Michaelis Menten representation of enzyme-substrate degradation reaction between ABA and H where $K_d$ [nM] and $\gamma_A$ [1/s] are the Michaelis constant and rate constant respectively for formation of PA. Reaction (R4) accounts for the degradation of enzyme H over time through a rate constant $\gamma_H$ [1/s].

$$\xrightarrow[K_-]{V_-} \mathrm{H} \qquad \text{[R2]}$$

$$\mathrm{ABA} + \mathrm{H} \underset{}{\overset{K_d}{\rightleftarrows}} \mathrm{ABA\!:H} \xrightarrow{\gamma_A} \mathrm{PA} + \mathrm{H} \qquad \text{[R3]}$$

$$\mathrm{H} \xrightarrow{\gamma_H} \qquad \text{[R4]}$$

**B. Abscisic Acid mediated protein signaling.** ABA has been hypothesized to be a primary regulator of stomatal movements under water deficit conditions[24,25,35,36]. Mutant plants deficient in ABA biosynthesis or signal transduction consistently exhibit severe drought-susceptible phenotypes, underscoring the critical role of ABA in mediating plant responses to water stress[37]. Under water stress, rapid *de novo* biosynthesis of ABA in guard cells has been shown to be involved in provoking stomatal responses[24]. Key proteins involved in ABA signaling have been well-characterized, including PYR receptors, PP2C protein phosphatases, OST1 protein kinases, and the SLAC1 anion channel. These components form the primary ABA signaling cascade responsible for stomatal closure, functioning independently of other signaling pathways.

Increasing evidence highlights the involvement of secondary messengers, such as $Ca^{2+}$, in stomatal closure[38]. While $Ca^{2+}$ acts as a secondary messenger in the ABA pathway, its role is complementary rather than essential. Experimental studies have demonstrated that ABA can regulate stomatal closure independently of cytosolic $Ca^{2+}$ elevation[39,40]. Once ABA-dependent signaling inactivates PP2C phosphatases, $Ca^{2+}$ can activate the SLAC1 anion channel through calcium-dependent protein kinase (CPK) activity, as observed in *in vitro* expression systems[41]. However, at higher ABA concentrations, the requirement for CPK activity is bypassed, as evidenced by studies on *cpk* mutants[41], indicating that the core ABA pathway alone is sufficient to drive stomatal responses. Given this sufficiency, we focus our model on the primary ABA pathway. This allows us to capture the essential regulatory mechanisms driving stomatal closure while avoiding the added complexity and uncertainties associated with secondary calcium-dependent processes. Although this framework maintains biological relevance, future integration of $Ca^{2+}$ pathways could offer further insights into the coordination and fine-tuning of stomatal closure mechanisms.

***B.1. Bicyclic signaling cascade.*** $OST1$ is a protein kinase that plays an important role in abiotic water stress signaling via ABA[42]. OST1 belongs to the SnRK2 family among which, three SnRK2.2, SnRK2.3, and OST1/SnRK2.6 are strongly activated in response to ABA. These genes are collectively represented by OST1 in this model. In guard cell cytoplasm, $OST1$ can exist in either a phosphorylated ($OST1^*$) or

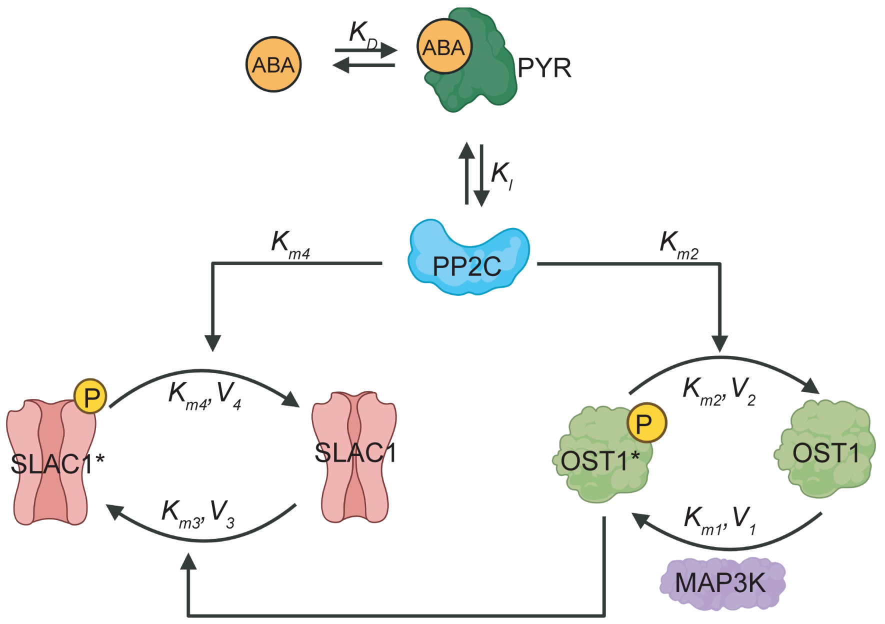

**Fig. S3. ABA signaling mediated by cascading futile cycles**. ABA binds its receptor PYR to form the ABA:PYR complex (disassociation constant, $K_D$ $[nM]$). This complex non-competitively inhibits phosphatase PP2C by binding allosteric sites (inhibition disassociation constant, $K_I$ $[nM]$). PP2C regulates a bicyclic cascade of futile cycles: in the first PP2C dephosphorylates protein kinase OST1* into its inactive form OST1 (Michaelis-Menten $K_{m2}$ $[nM]$, reaction velocity $V_2$ $[nM/s]$) while MAP3K phosphorylates OST1 back into its active form OST1* (Michaelis-Menten $K_{m1}$ $[nM]$, reaction velocity $V_1$ $[nM/s]$); in the second, PP2C dephosphorylates the open form of the anion channel, SLAC1* to its closed form SLAC1 (Michaelis-Menten $K_{m3}$ $[nM]$, reaction velocity $V_3$ $[nM/s]$) while OST1* phosphorylates SLAC1 into its open state SLAC1* (Michaelis-Menten $K_{m4}$ $[nM]$, reaction velocity $V_4$ $[nM/s]$).The governing equations for this circuit are provided in Section S3B (Eqs. S72 and S80).

a dephosphorylated ($OST1$) state. The activation of $OST1$ into its phosphorylated state is carried out by kinase $MAP3K$[43]. Its deactivation into dephosphorylated state is carried out by phosphatase $PP2C$ (HAB1/HAB2/ABI1/ABI2)[38,44–46]. These two pathways form a "futile cycle". Reactions (R5) and (R6) capture the two phosphorylation and dephosphorylation reaction for $OST1$ represented in Michaelis Menten enzyme-substrate reaction form.

$$\mathrm{MAP3K} + \mathrm{OST1} \underset{d_1}{\overset{a_1}{\rightleftarrows}} \mathrm{MAP3K}:\mathrm{OST1} \xrightarrow{k_1} \mathrm{OST1}^* + \mathrm{MAP3K} \qquad [\mathrm{R5}]$$

$$\mathrm{PP2C} + \mathrm{OST1}^* \underset{d_2}{\overset{a_2}{\rightleftarrows}} \mathrm{PP2C}:\mathrm{OST1}^* \xrightarrow{k_2} \mathrm{OST1} + \mathrm{PP2C} \qquad [\mathrm{R6}]$$

Downstream, phosphorylated $OST1$ is a part of another futile cycle for $SLAC1$ anion channel as a kinase phosphorylating $SLAC1$. Phosphatase $PP2C$ is also responsible for dephosphorylating $SLAC1$[47,48]. Such a signaling motif formed by $OST1$ and $SLAC1$ together, is called a "bicyclic cascade" where the output of $OST1$ futile cycle forms the input to a downstream $SLAC1$ futile cycle. In our model of ABA-mediated signaling, SLAC1 is considered the primary mediator of anion flux. Loss of SLAC1 leads to plants with open stomata that displayed impaired responses to drop in the air humidity[49]. Hsu[50] showed that SLAC1 plays a relatively more substantial role than the rapid-type anion channel ALMT12/QUAC1 in stomatal VPD signaling. Data from patch-clamp study of mutants impaired in ABA signal transduction suggested that the S-type anion channel is more clearly involved in stomatal closing[51]. Reactions (R7) and (R8) capture the two phosphorylation and dephosphorylation reactions for $SLAC1$ represented in Michaelis Menten enzyme-substrate reaction form.

$$\mathrm{OST1}^* + \mathrm{SLAC1} \underset{d_3}{\overset{a_3}{\rightleftarrows}} \mathrm{OST1}^*:\mathrm{SLAC1} \xrightarrow{k_3} \mathrm{OST1}^* + \mathrm{SLAC1}^* \qquad [\mathrm{R7}]$$

$$\mathrm{PP2C} + \mathrm{SLAC1}^* \underset{d_4}{\overset{a_4}{\rightleftarrows}} \mathrm{PP2C}:\mathrm{SLAC1}^* \xrightarrow{k_4} \mathrm{PP2C} + \mathrm{SLAC1} \qquad [\mathrm{R8}]$$

***B.2. Non-competitive inhibition of PP2C.*** ABA is important for activation of this $OST1 - SLAC1$ bicyclic signal transduction cascade for stomatal regulation. In absence of ABA, the $OST1$ futile cycle is biased in favor of dephosphorylated OST1. In presence of ABA, ABA-mediated signaling begins with the binding of ABA to its receptor PYR in the guard cell cytoplasm to form a complex $ABA : PYR$ (reaction R9). Details regarding the specific proteins PYR/PYL/RCAR that participate in this reaction can be found in Dupeux [52]. This complex has high affinity to bind with PP2C [44,45] (reaction R10). $ABA\colon PYR$ complex binding to PP2C results in non-competitive inhibition of PP2C [44,45] leading to a decrease in dephosphorylation activity of PP2C. In non-competitive inhibition, the inhibitor ($ABA\colon PYR$) binds on sites of enzyme ($PP2C$) other than the active site and reduces enzyme affinity to the substrate (reactions R11-R14). The inhibition biases the $OST1$ futile cycle in favor of phosphorylated state of $OST1$. Once phosphorylated, $OST1$ goes on to phosphorylate $SLAC1$ which allows guard cell to regulate stomatal conductance via osmotic adjustment.

$$\mathrm{ABA} + \mathrm{PYR} \xrightleftharpoons{K_D} \mathrm{ABA\colon PYR} \qquad \text{[R9]}$$

$$\mathrm{PP2C} + \mathrm{ABA\colon PYR} \xrightleftharpoons{K_I} \mathrm{PP2C : ABA\colon PYR} \qquad \text{[R10]}$$

$$\mathrm{PP2C : OST1^*} + \mathrm{ABA\colon PYR} \xrightleftharpoons{K_I} \mathrm{PP2C : OST1^* : ABA\colon PYR} \qquad \text{[R11]}$$

$$\mathrm{PP2C : SLAC1^*} + \mathrm{ABA\colon PYR} \xrightleftharpoons{K_I} \mathrm{PP2C : SLAC1^* : ABA\colon PYR} \qquad \text{[R12]}$$

$$\mathrm{PP2C : ABA\colon PYR} + \mathrm{OST1^*} \xrightleftharpoons{K_{m2}} \mathrm{PP2C : OST1^* : ABA\colon PYR} \qquad \text{[R13]}$$

$$\mathrm{PP2C : ABA\colon PYR} + \mathrm{SLAC1^*} \xrightleftharpoons{K_{m4}} \mathrm{PP2C : SLAC1^* : ABA\colon PYR} \qquad \text{[R14]}$$

Here, $K_D$ [nM] is the disassociation constant for ABA binding to its receptor $PYR$, $K_I$ [nM] is the disassociation constant for $ABA : PYR$ inhibiting $PP2C$ as a result of their binding, $K_{m2}$ [nM] and $K_{m4}$ [nM] are the Michaelis constants for $OST1^*$ and $SLAC1^*$ dephosphorylation reactions respectively and also the binding constants for $PP2C : ABA : PYR$ to $OST1^*$ and $SLAC1^*$ respectively [53].

***B.3. Model assumptions and reaction parameters.***

1. **Assumptions:**

   We make the following assumptions for the reactions listed above:

   $(i)$ Quasi-steady state assumption for complexes in reactions (R3) and (R5) - (R8).

   $(ii)$ Rapid equilibrium assumption for reactions (R9) - (R14).

2. **Defining reaction constants**

   $(i)$ **Michaelis Menten reaction constants:**

$$K_2 = \frac{[ABA][H]}{[ABA : H]} \qquad \text{[S40]}$$

$$K_{m1} = \frac{d_1 + k_1}{a_1} = \frac{[MAP3K][OST1]}{[MAP3K : OST1]} \qquad \text{[S41]}$$

$$K_{m2} = \frac{d_2 + k_2}{a_2} = \frac{[PP2C][OST1^*]}{[PP2C : OST1^*]} \quad [S42]$$

$$K_{m3} = \frac{d_3 + k_3}{a_3} = \frac{[OST1^*][SLAC1]}{[OST1^* : SLAC1]} \quad [S43]$$

$$K_{m4} = \frac{d_4 + k_4}{a_4} = \frac{[PP2C][SLAC1^*]}{[PP2C : SLAC1^*]} \quad [S44]$$

The parameters in Eqs. S40 and S44 are provided in Table S7.

$(ii)$ **Equilibrium relationships:**

$$K_D = \frac{k_{-D}}{k_D} = \frac{[ABA][PYR]}{[ABA : PYR]} \quad [S45]$$

$$K_I = \frac{k_{-I}}{k_I} = \frac{[ABA : PYR][PP2C]}{[PP2C : ABA : PYR]} = \frac{[PP2C : OST1^*][ABA : PYR]}{[PP2C : OST1^* : ABA : PYR]} = \frac{[PP2C : SLAC1^*][ABA : PYR]}{[PP2C : SLAC1^* : ABA : PYR]} \quad [S46]$$

$$K_{m2} = \frac{[PP2C : ABA : PYR][OST1^*]}{[PP2C : OST1^* : ABA : PYR]} \quad [S47]$$

$$K_{m4} = \frac{[PP2C : ABA : PYR][SLAC1^*]}{[PP2C : SLAC1^* : ABA : PYR]} \quad [S48]$$

The parameters in Eqs. S45 and S46 are defined as follows and values are provided in Table S8.

- $k_D$, $k_I$ [1/nM/s]: rates of forward reaction
- $k_{-D}$, $k_{-I}$ [1/s]: rates of backward reaction.

#### B.4. Governing equations.

$(a)$ **Abscisic acid (ABA)**

The free cytosolic abscisic acid $[ABA]$ in the cytosol can be accounted from reaction (R1), (R3), and (R9) as follows:

$(i)$ **ABA synthesis:**

The basal production and auto-upregulation of ABA production are represented as:

$$\theta_{syn}\left\{V_{stress}(\delta P_g) + V_+\left(\frac{[ABA]^n}{K_+^n + [ABA]^n}\right)\right\} \quad [S49]$$

ABA-synthesis mutant conditions are simulated in our model by reducing the effectiveness of the ABA biosynthesis pathway, encompassing both *de novo* synthesis and positive auto-regulation. To achieve this, a pre-factor $\theta_{syn}$ where, $0 \leq \theta_{syn} \leq 1$, with $\theta_{syn} = 1$ for wildtype is multiplied to the net ABA synthesis (R1).

$(ii)$ **ABA degradation:**

The autoregulatory degradation of ABA by H can be represented as:

$$-\gamma_A[H]\Big(\frac{[ABA]}{K_d+[ABA]}\Big) \quad [S50]$$

$(iii)$ **ABA depletion:**

From rapid equilibrium assumption, ABA depletion caused by binding to its receptor can be represented as:

$$-k_d[H][PYR]+k_{-d}[ABA:PYR] \quad [S51]$$

$(iv)$ **Net time rate of change of free cytosolic $ABA$:**

$$\frac{d[ABA]}{dt}=\theta_{syn}\Bigg\{V_{stress}(\Delta P_g)+V_+\Big(\frac{[ABA]^n}{K_+^n+[ABA]^n}\Big)\Bigg\}-\gamma_A[H]\Big(\frac{[ABA]}{K_d+[ABA]}\Big)- k_d[H][PYR]+k_{-d}[ABA:PYR] \quad [S52]$$

From a rapid equilibrium assumption for ABA binding to its receptor PYR[31] in Eq. S51,

$$\frac{d[ABA]}{dt}=\theta_{syn}\Bigg\{V_{stress}(\Delta P_g)+V_+\Big(\frac{[ABA]^n}{K_+^n+[ABA]^n}\Big)\Bigg\}-\gamma_A[H]\Big(\frac{[ABA]}{K_d+[ABA]}\Big) \quad [S53]$$

$(b)$ **8'-hydroxylase (H)**

$(i)$ **H synthesis:**

ABA dependent upregulation of H:

$$V_-\Big(\frac{[ABA]}{K_-+[ABA]}\Big) \quad [S54]$$

$(ii)$ **H degradation:**

H is an enzyme that degrades over time as:

$$-\gamma_H[H] \quad [S55]$$

$(iii)$ **Net time rate of change of free cytosolic $H$:**

$$\frac{d[H]}{dt}=V_-\Big(\frac{[ABA]}{K_-+[ABA]}\Big)-\gamma_H[H] \quad [S56]$$

$(c)$ **Mass Balance on proteins**

$(i)$ **ABA cytosolic receptor PYR:**

$$[PYR]_T=[PYR]+[ABA:PYR]+[PP2C:ABA:PYR]+ [PP2C:OST1^*:ABA:PYR]+[PP2C:SLAC1^*:ABA:PYR] \quad [S57]$$

(ii) **MAP3K kinase:**

$$[MAP3K]_T = [MAP3K] + [MAP3K : OST1] \quad [S58]$$

(iii) **PP2C phosphatase:**

$$[PP2C]_T = [PP2C] + [PP2C : OST1^*] + [PP2C : SLAC1^*] + [PP2C : ABA : PYR] + [PP2C : OST1^* : ABA : PYR] + [PP2C : SLAC1^* : ABA : PYR] \quad [S59]$$

(iv) **OST1 kinase:**

$$[OST1]_T = [OST1] + [OST1^*] + [MAP3K : OST1] + [PP2C : OST1^*] + [OST1^* : SLAC1] + [PP2C : OST1^* : ABA : PYR] \quad [S60]$$

(v) **SLAC1 ion channel:**

$$[SLAC1]_T = [SLAC1] + [SLAC1^*] + [OST1^* : SLAC1] + [PP2C : SLAC1^*] + [PP2C : SLAC1^* : ABA : PYR] \quad [S61]$$

Where, $[X]_T$ is the total concentration of the protein in the guard cell cytoplasm.

(d) **Rate of change of phosphorylated $OST1^*$:**

(i) $OST1^*$ **formation:**

From reactions (R5), (R6), and (R7)

$$k_1[MAP3K : OST1] + d_2[PP2C : OST1] + d_3[OST1^* : SLAC1] + k_3[OST1^* : SLAC1] \quad [S62]$$

(ii) $OST1^*$ **depletion:**

From reactions (R6) and (R7)

$$-a_2[PP2C][OST1^*] - a_3[OST1^*][SLAC1] \quad [S63]$$

(iii) **Net time rate of change of $OST1^*$:**

$$\frac{d[OST1^*]}{dt} = k_1[MAP3K : OST1] + d_2[PP2C : OST1] + d_3[OST1^* : SLAC1] + k_3[OST1^* : SLAC1] - a_2[PP2C][OST1^*] - a_3[OST1^*][SLAC1] \quad [S64]$$

From Eq. S43,

$$-a_3[OST1^*][SLAC1] + (d_3 + k_3)[OST1^* : SLAC1] = 0 \quad [S65]$$

Using Eq. S65 in Eq. S64,

$$\frac{d[OST1^*]}{dt} = k_1[MAP3K : OST1] + d_2[PP2C : OST1] - a_2[PP2C][OST1^*] \quad [S66]$$

Using mass balances from Eqs. S57-S61 and equilibrium equations Eqs. S41-S48 in Eq. S66,

$$\frac{d[OST1^*]}{dt} = k_1[MAP3K]_T\Big(\frac{[OST1]}{K_{m1} + [OST1]}\Big) - \frac{k_2[PP2C]_T}{(1 + \frac{[ABA:PYR]}{K_I})}\Bigg(\frac{[OST1^*]}{K_{m2}(1 + \frac{[SLAC1^*]}{K_{m4}}) + [OST1^*]}\Bigg) \quad [S67]$$

Where $[OST1]$ can be evaluated from solving mass balance in Eq. S60.

Define:

$$V_1 = k_1[MAP3K]_T \quad [S68]$$

$$V_2^{max} = k_2[PP2C]_T \quad [S69]$$

$$V_2 = \frac{k_2[PP2C]_T}{\Big(1 + \frac{[ABA:PYR]}{K_I}\Big)} \quad [S70]$$

$$K'_{m2} = K_{m2}\Big(1 + \frac{[SLAC1^*]}{K_{m4}}\Big) \quad [S71]$$

Eq. S67 can be re-written as:

$$\frac{d[OST1^*]}{dt} = V_1\Big(\frac{[OST1]}{K_{m1} + [OST1]}\Big) - V_2\Big(\frac{[OST1^*]}{K'_{m2} + [OST1^*]}\Big) \quad [S72]$$

**$(e)$ Rate of change of phosphorylated $SLAC1^*$:**

**$(i)$ $SLAC1^*$ formation:**

From reactions (R7) and (R8)

$$k_3[OST1^* : SLAC1] + d_4[PP2C : SLAC1^*] \quad [S73]$$

**$(ii)$ $SLAC1^*$ depletion:**

From reactions (R7) and (R8)

$$-a_4[PP2C][SLAC1^*] \quad [S74]$$

**$(iii)$ Net time rate of change of $SLAC1^*$:**

$$\frac{d[SLAC1^*]}{dt} = k_3[OST1^* : SLAC1] + d_4[PP2C : SLAC1^*] - a_4[PP2C][SLAC1^*] \quad [S75]$$

Using mass balances from Eqs. S57-S61 and equilibrium equations Eqs. S41-S48 in Eq. S75,

$$\frac{d[SLAC1^*]}{dt} = \frac{k_3[OST1^*]}{K_{m3} + [OST1^*]}\left([SLAC1]_T - \frac{[PP2C]_T[SLAC1^*]}{K_{m4}(1 + \frac{[OST1^*]}{K_{m2}}) + [SLAC1^*]}\right) - \frac{k_4[PP2C]_T}{(1 + \frac{[ABA:PYR]}{K_I})}\left(\frac{[SLAC1^*]}{K_{m4}(1 + \frac{[OST1^*]}{K_{m2}}) + [SLAC1^*]}\right) \quad [S76]$$

Where $[OST1^*]$ can be evaluated from solving ODE in Eq. S72.

Define:

$$V_4^{max} = k_4[PP2C]_T \quad [S77]$$

$$V_4 = \frac{k_4[PP2C]_T}{\left(1 + \frac{[ABA:PYR]}{K_I}\right)} \quad [S78]$$

$$K'_{m4} = K_{m4}\left(1 + \frac{[OST1^*]}{K_{m2}}\right) \quad [S79]$$

Eq. S76 can be re-written as:

$$\frac{d[SLAC1^*]}{dt} = \frac{k_3[OST1^*]}{K_{m3} + [OST1^*]}\left([SLAC1]_T - \frac{[PP2C]_T[SLAC1^*]}{K'_{m4} + [SLAC1^*]}\right) - V_4\left(\frac{[SLAC1^*]}{K'_{m4} + [SLAC1^*]}\right) \quad [S80]$$

The rate processes encoded in Eqs. S72 and S80 represent cascading futile cycles. Futile cycles which consist of opposing enzymatic reactions (e.g., phosphorylation and dephosphorylation), can exhibit first-order or zeroth-order response kinetics depending on the relative concentrations of enzymes and substrates. These responses significantly influence the dynamics and regulatory properties of the cycle.

**Conditions for First-Order Kinetics in Futile Cycles**: The substrate concentration ($[S]$) is low relative to the Michaelis constant of the enzyme ($K_m$). The reaction operates in the linear portion of the enzyme's activity curve, where the enzyme is far from saturation: $[S] << K_m$

**Conditions for Zeroth-Order Kinetics in Futile Cycles**: The substrate concentration ($[S]$) is high relative to the Michaelis constant of the enzyme ($K_m$). The enzyme is saturated, meaning all active sites are occupied, and further increases in substrate do not affect the reaction rate: $[S] >> K_m$

The parameters for futile cycle reactions can be found in Table S7 and can be compared with protein concentrations in Table S9. In our study, reactions (R5), (R7), and (R8) operate with first-order kinetics while reaction (R6) operates with zeroth-order kinetics.

**C. Guard cell membrane electrophysiology.** In this section, we describe the guard cell membrane transport mechanisms incorporated into our model. We adopt a semi-mechanistic approach, coarse-graining the system to include a representation of transporters that are necessary to evaluate the osmotic potential changes driving stomatal closure under ABA regulation via key OST1-gated anion (SLAC1) and voltage-gated cation (GORK and KAT) channels. Specifically, we model $Cl^-$ as the constitutive anion and $K^+$ as the constitutive cation, alongside proton movement driven by $H^+$-ATPase pumps and $2H^+/Cl^-$ symporters[10,54]. We incorporate the specific protein-level regulation details (SLAC1), and voltage-gating characteristics of these transporters (KAT1, GORK, $H^+$-ATPase pump, and $2H^+/Cl^-$ symporter) as established in the literature.

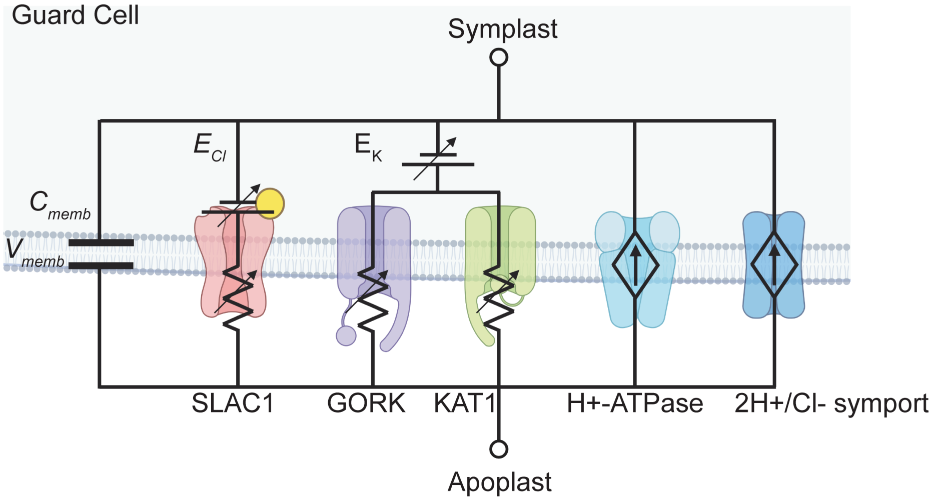

**Fig. S4. Modeling of guard cell electrophysiology**. Circuit representation of membrane features and processes represented explicitly in our model. This representation follows that of Bertil Hille[13]. The guard cell plasma membrane has a capacitance, $C_{memb}$ $[F/m^2]$ and changes in ion concentrations lead to changes in plasma membrane voltage, $V_{memb}$ $[mV]$. SLAC1, GORK, and KAT are voltage-gated ion channels for chloride (SLAC1) and potassium (GORK and KAT1). These channels act as variable resistances for their respective ions (jagged lines with cross-arrow). Their reversal potentials are: $E_{Cl}$ $[V]$ (SLAC1) and $E_K$ $[V]$ (GORK and KAT1). The $H^+$-ATPase pump and 2H/Cl symporter act as current sources that regulate ion fluxes and membrane voltage to maintain guard cell homeostasis. The activation of SLAC1* with elevated ABA facilitates the efflux of $Cl^-$ ions, leading to depolarization of the guard cell plasma membrane. This depolarization opens the GORK cation channel, allowing $K^+$ efflux and resulting in a decrease in guard cells osmotic potential ($\delta\bar{\pi}_g$ $[MPa]$). This change in osmotic potential in guard cells feeds back into change in guard cell turgor pressure and stomatal conductance. The governing equations for this circuit are provided in Section S3C (Eqs. S100-S102).

#### *C.1. Model assumptions and validation.*

($a$) **Ion conservation and electrodiffusion:** We assume no buffering of concentrations for the constitutive ions $Cl^-$ and $K^+$ in the apoplast. This treatment is grounded in the classical Hodgkin-Huxley formalism[55]. Consequently, the concentration of these ions in the apoplast is dictated by a mass balance across the symplast and apoplast (Eq. S96), with transmembrane currents generated from electrodiffusion modeled by the Goldman-Hodgkin-Katz (GHK) equations[13] (Eqs. S82, S84, and S87).

($b$) **Proton dynamics and pH buffering:** While currents from all transporters contribute to the evolution of the membrane potential, we restrict our explicit tracking of concentration dynamics to $K^+$ and $Cl^-$, treating apoplastic and symplastic pH as constant. This simplification is supported by the following considerations:

- *Negligible Osmotic Contribution:* $[H^+]$ is orders of magnitude lower than $[K^+]$ and $[Cl^-]$ (Table S10), rendering its direct contribution to the guard cell osmotic potential negligible.
- *Tight Buffering:* Experimental evidence[56,57] and detailed electrophysiological models[10,11] demonstrate that the guard cell apoplasm and symplasm exhibit tight pH buffering. By fixing pH, we effectively capture this buffering capacity without explicitly modeling the proton dynamics.
- *Physiological Consistency:* We verified that this assumption does not compromise the electrophysiological accuracy. While our instantaneous transient $H^+$ fluxes are higher than steady-state values reported in the OnGuard model[11], potentially due to the different nature of our perturbation, we verified that the resulting intracellular pH change ($\Delta$pH $\leq 0.5$) is quantitatively consistent with observations in Blatt et al.[11] with the $H^+$ buffering rates provided in literature[56,57].

(c) **Validation of transporter kinetics:** We benchmarked the steady-state current-voltage (I-V) profiles of our key transporters against the OnGuard2 model[12]. Our implementation of the GORK channel ($I_{K_{out}}$) reproduces the non-linear voltage dependence and current magnitudes observed in previous computational and experimental studies[12].

***C.2. Description of transport proteins.*** In the following section, we provide descriptions of transport proteins. All transport proteins show a voltage dependence. Hence, we define scaled voltage as:

$$u = \frac{V_{memb}F}{RT} \quad \text{[S81]}$$

The parameters in Eq. S81 are defined as follows and values are provided in Table S10.

- $F$ [C/mol]: Faraday constant
- $V_{memb}$ [mV]: guard cell membrane potential

(a) **Ion channels**

The $SLAC1$ anion channel is the primary transporter of $Cl^-$. The cation channels considered are $GORK$ and $KAT1$ which are responsible for movement of $K^+$ ion outside and inside of the guard cell cytoplasm respectively. For all ion channels, we represent current due to charge movement through them using the Goldman Hodgkin Katz (GHK) representation[13]. This representation is particularly useful when the concentration of ions inside the cell is changing.

(i) **Anion channel - SLAC1:**

$$I_{Cl}^{SLAC1} = P_o^{SLAC1} P_{Cl} z_{Cl}^2 uF \left[ \frac{[Cl^-]_{in} - [Cl^-]_{out} \exp\Big( -z_{Cl}u \Big)}{1 - \exp\Big( -z_{Cl}u \Big)} \right] \quad \text{[S82]}$$

$$P_o^{SLAC1} = \frac{[SLAC1^*]}{[SLAC1]_T} \quad \text{[S83]}$$

The parameters in Eqs. S82 and S83 are defined as follows and values are provided in Table S10.

- $I_{Cl}^{SLAC1}$ [A/m$^2$]: current due to movement of chloride ions
- $P_o^{SLAC1}$ [$-$]: open probability of SLAC1 ion channels
- $P_{Cl}$ [m/s]: permeability of chloride ions through SLAC1 channel
- $z_{Cl}$ [$-$]: valence of chloride ion
- $[Cl^-]_{in}$ [mM]: concentration of chloride ions inside the guard cell in symplast
- $[Cl^-]_{out}$ [mM]: concentration of chloride ions outside the guard cell in apoplast

$GORK$ and $KAT1$ have a voltage dependence[54]. For more negative voltages, $GORK$ inactivates and $KAT1$ is open allowing a passage for influx of $K^+$ ions. The negative potentials generated by the $H^+ - ATPase$ ensure that the hyperpolarized membrane creates enough driving force for $K^+$ uptake even in the case the concentration of $K^+$ in the apoplast is smaller than the symplast.

(ii) **Cation channel - GORK:**

$$I_K^{GORK} = P_o^{GORK} P_K z_K^2 uF \left[ \frac{[K^+]_{in} - [K^+]_{out} \exp\Big( -z_K u \Big)}{1 - \exp\Big( -z_K u \Big)} \right] \quad \text{[S84]}$$

$$P_o^{GORK} = \frac{1}{1 + \exp\Big( - (V_{memb} - V_{1/2}^{GORK}) * \frac{\delta_{GORK} F}{RT}\Big)} \quad \text{[S85]}$$

$$V_{1/2}^{GORK} = \frac{z_K F}{RT} log\Big(\frac{[K^+]_{out}}{K_{GORK}}\Big) \quad \text{[S86]}$$

$(iii)$ **Cation channel - KAT1:**

$$I_K^{KAT1} = P_o^{KAT1} P_K z_K^2 u F \left[\frac{[K^+]_{in} - [K^+]_{out} \exp\Big( - z_K u\Big)}{1 - \exp\Big( - z_K u\Big)}\right] \quad \text{[S87]}$$

$$P_o^{KAT1} = \frac{1}{1 + \exp\Big( - (V_{memb} - V_{1/2}^{KAT1}) * \frac{\delta_{KAT1} F}{RT}\Big)} \quad \text{[S88]}$$

The parameters in Eqs. S84-S88 are defined as follows and values are provided in Table S10.

- $I_K^{GORK,KAT1}$ [A/m$^2$]: current due to movement of potassium ions
- $P_o^{GORK,KAT1}$ [$-$]: open probability of GORK and KAT1 ion channels
- $P_K$ [m/s]: permeability of potassium through GORK and KAT1 channel
- $z_K$ [$-$]: valence of potassium ion
- $[K^+]_{in}$ [mM]: concentration of potassium ions inside the guard cell in symplast
- $[K^+]_{out}$ [mM]: concentration of potassium ions outside the guard cell in apoplast
- $\delta_{GORK,KAT1}$ [$-$]; $V_{1/2}^{GORK,KAT1}$ [mV]; $K_{GORK}$: parameters for gating probabilities of GORK and KAT1 channels

$(b)$ **H-ATPase pump:**

The $H^+$-ATPase pump serves as the primary electrogenic mechanism, driving proton efflux to hyperpolarize the guard cell membrane. Consistent with the buffering principles and scale separation outlined above, we maintain fixed apoplastic and symplastic proton concentrations in the pump's kinetic description. However, to accurately capture the pump's contribution to membrane potential dynamics, we must retain its voltage dependence. Therefore, we adopt the reaction-kinetic formalism of Blatt et al.[10,58,59]. This model captures the non-linear response of pump current ($I_{Hpump}$) to membrane voltage ($u$), while assuming that ATP availability remains sufficient and non-limiting[10,54]. The voltage-current properties are given by:

$$I_{Hpump} = P_o^{Hpump} E_o \left(\frac{k_{+1}k_{+2} - k_{-1}k_{-2}}{k_{+1} + k_{+2} + k_{-1} + k_{-2}}\right) \quad \text{[S89]}$$

$$k_{+1} = k_1 \bar{H}_{in}^{+} \quad \text{[S90]}$$

$$k_{+2} = k_2\Big(\frac{u}{1 - \exp(-u)}\Big) \quad \text{[S91]}$$

$$k_{-1} = k_1 \exp\Big(\frac{\Delta G_{ATP}}{RT}\Big) \quad \text{[S92]}$$

$$k_{-2} = k_2 \bar{H}^{+}_{out}\Big(\frac{u * \exp(-u)}{1 - \exp(-u)}\Big) \quad [S93]$$

The parameters in Eq. S89-S93 are defined as follows and values are provided in Table S10.

- $P_o^{Hpump}$ $[-]$: on probability of H-ATPase pump
- $I_{Hpump}$ [A/m$^2$]: current due to movement of protons out of the cell
- $E_o$ [C/m$^2$]: H-ATPase maximum charge pumping capacity
- $\bar{H}^{+}_{in}$, $\bar{H}^{+}_{out}$[-]: normalized concentration of $H^+$ ions (normalized by 1mM)
- $k_{+1}$, $k_{+2}$, $k_{-1}$, $k_{-2}$, [1/s]: H-ATPase parameters[58,59]
- $\Delta G_{ATP}$ [kJ/mol]: energy from ATP hydrolysis

The proton pump is inhibited by application of ABA[60]. For this inhibition, we assume a first order Hill-type repression function:

$$P_o^{Hpump} = \frac{1}{\Big(1 + \frac{[ABA]}{K_{ABA}}\Big)} \quad [S94]$$

Where, $K_{ABA}$ [nM] is the ABA dependent pumping probability of H-ATPase pump (Table S10).

(*c*) **2H/Cl symport:**

The proton pumps use energy to extrude $H^+$ out of the cell against a gradient of $H^+$ ions across the cell. This gradient of ions from apoplast to symplast is used to carry $Cl^-$ into the cell. The $2H^+/Cl^-$ membrane symporters bring in one $Cl^-$ ion from every two $H^+$ ions that enter the cell along the concentration gradient[61].

$$I_{Cl}^{sym} = P_o^{sym} V_o\left(\frac{[Cl^-]_{in}[H^+]^2_{in} * u - [Cl^-]_{out}[H^+]^2_{out} * u * \exp(-u)}{1 - \exp(-u)}\right) \quad [S95]$$

The parameters in Eq. S95 are defined as follows and values are provided in Table S10.

- $I_{Cl}^{sym}$ [A/m$^2$]: symporter current
- $P_o^{sym}$ $[-]$: 'on' probability of 2H/Cl symporter
- $V_o$ [C/mM$^3$.s.m$^2$]: max symporter current

#### C.3. Governing equations.

(*a*) **Mass balance**

The total moles of ions across the symplast and apoplast remain conserved.

$$Q_X = [X]^0_{in} vol_{in} + [X]^0_{out} vol_{out} = [X]_{in} vol_{in} + [X]_{out} vol_{out} \quad [S96]$$

The parameters in Eq. S96 are defined as follows and values are provided in Table S10.

- $[X]^0_{in}$ [mM]: reference value of symplastic concentration of ion X
- $[X]^0_{out}$ [mM]: reference value of apoplastic concentration of ion X
- $[X]_{in}$ [mM]: instantaneous symplastic concentration of ion X

- $[X]_{out}$ [mM]: instantaneous apoplastic concentration of ion X
- $vol_{in}$ [m$^3$]: Symplast volume
- $vol_{out}$ [m$^3$]: Apoplast volume

The initial conditions define the concentrations of ions in the two compartments. The instantaneous concentration of ion X in the apoplast is given by:

$$[X]_{out} = \frac{[X]_{in}^0 vol_{in} + [X]_{out}^0 vol_{out} - [X]_{in} vol_{in}}{vol_{out}} \qquad \text{[S97]}$$

Eq. S97 requires the volume of guard cell ($vol_{in}$) be expressed in [m$^3$]. In our HP model (see Section S2), we use the volume of guard cell scaled per leaf unit area $V_g^0$ [mmol/m$^2$]. To get $vol_{in}$ [m$^3$] from $V_g^0$ [mmol/m$^2$] using stomatal density ($\phi_g$ [#/m$^2$ leaf area]):

$$vol_{in} = V_g^0 * \bar{\nu}_w / \phi_g \qquad \text{[S98]}$$

The mass balance in Eqs. S101 and S102 is applied to $Cl^-$ and $K^+$ ions respectively. Using the concentrations of $Cl^-$ and $K^+$ ion, the biochemically active change in osmotic potential in guard cell is evaluated as:

$$\delta\bar{\pi}_g = ([Cl^-]_{in} - [Cl^-]_{in}^0 + [K^+]_{in} - [K^+]_{in}^0) * RT \qquad \text{[S99]}$$

(*b*) **Rate of change**
All the currents contribute to change in membrane potential. Hence, in the rate of change of membrane potential, we account for currents from all transporters. For rate of change of ions, we hold $H^+$ ions constant (as noted above), and hence, we write the rate of change of ion concentration equations for $Cl^-$ and $K^+$ ions only.

(*i*) Membrane potential

$$\frac{dV_{memb}}{dt} = \frac{-1}{C_{memb}}(I_{Cl}^{SLAC1} + I_K^{GORK} + I_K^{KAT1} + I_{Hpump} + I_{Cl}^{symp}) \qquad \text{[S100]}$$

In Eq. S100, $C_{memb}$ [F/m$^2$] is the capacitance of the guard cell-membrane (Table S10).

(*ii*) Chloride concentration

$$\frac{d[Cl^-]_{in}}{dt} = \frac{-A_g}{z_{Cl} * F * vol_{in}}(I_{Cl}^{SLAC1} + I_{Cl}^{symp}) \qquad \text{[S101]}$$

(*iii*) Potassium concentration

$$\frac{d[K^+]_{in}}{dt} = \frac{-A_g}{z_K * F * vol_{in}}(I_K^{GORK} + I_K^{KAT1}) \qquad \text{[S102]}$$

In Eq. S102, $A_g$ [m$^2$] is the surface area of the guard cell-membrane (Table S10).

### S4. Coupling hydropassive and hydroactive models

The coupling between the hydropassive and hydroactive models occurs at two key points:

**A. Linking water status to ABA biosynthesis.** Changes in guard cell turgor pressure, driven by water status in the hydropassive model, trigger ABA biosynthesis. This represents water stress stimuli from the hydropassive model informing a biochemical process in the hydroactive model.

$$V_{stress}(\delta P_g) = V^0_{stress}\Big(\frac{\delta P_g^{3.25}}{1.4 + \delta P_g^{3.25}}\Big) \qquad \text{[S103]}$$

The parameters in Eq. S103 are defined as follows:

- $V^0_{stress}$ [nM/s]: Maximum stress dependent biosynthesis of Abscisic Acid in guard cells
- $\delta P_g$ [MPa]: Change in the turgor pressure of guard cells in response to water stress perturbation

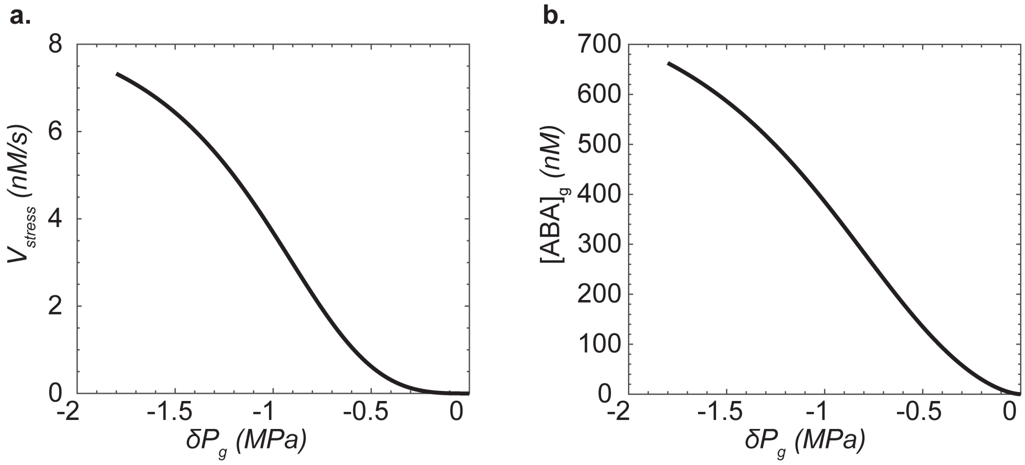

**Fig. S5. Stress dependent biosynthesis of ABA in guard cells. a.** Imposed relationship between ABA biosynthesis rate ($V_{stress}$ $[nM/s]$) and the change in guard cell turgor pressure from its reference value ($\delta P_g$ $[MPa]$) used in the model. The synthesis is described by a sigmoidal function (Eq. S103). **b.** Predicted steady-state guard cell ABA concentration ($[ABA]_g$ $[nM]$) as a function of change $\delta P_g$ based on relationship in (**a**).

The sigmoidal functional form of $V_{stress}$ in Eq. S103 was chosen to ensure the model captures the non-linear dependence of steady-state ABA levels on turgor pressure, as reported in angiosperms[27,28]. These studies demonstrate that steady-state ABA concentrations rise as leaf turgor declines. By defining the synthesis rate as a sigmoidal function of turgor change (Fig. S5a), the model yields steady-state ABA concentrations that reproduce this experimentally observed non-linear accumulation (Fig. S5b).

**B. Feedback on turgor pressure.** Changes in guard cell osmotic potential, governed by the hydroactive model, feedback to regulate turgor pressure. This physiological adjustment affects stomatal conductance, completing the feedback loop between the two models. Mathematically, the coupling occurs as follows: changes in osmotic pressure (Eq. S99), lead to changes in guard cell water potential (Eq. S10), and these changes in water potential drive flows between the guard and subsidiary cells (Eqs. S38 and S39).

### S5. Analyzing the Abscisic Acid autoregulation motif

As discussed in the main text, we identify ABA auto-regulation as the core regulatory motif governing ABA dynamics. Here, we provide further mathematical analysis of these key governing equations to aid in their parameterization and gain insights into the importance of specific component rate processes. We begin by non-dimensionalizing the governing equations, identifying key dimensionless groups that control both the steady-state behavior and transient dynamics of the system. These non-dimensional forms enable us to extract mechanistic insights and place constraints on otherwise uncertain parameters relevant to stomatal conductance regulation in *Arabidopsis thaliana* genotypes. We further show how specific dimensionless ratios shape the features of the system: (i) dynamic features defining the emergence and nature of oscillatory behavior; and (ii) steady-state features defining the bi-stability that leads to hysteresis. The section concludes with a focused examination of how the ABA regulatory motif contributes to the onset and characteristics of such oscillations in the coupled model.

**A. Defining non-dimensional numbers and time-scales.** Let Abscisic Acid (ABA) be denoted by $'A'$ and 8'-hydroxylase enzyme (A8H) be denoted by $'H'$. The rate equations for regulation of ABA are:

$$\frac{d[A]}{dt} = \theta_{syn}\left[V_{stress} + V_{+}\Big(\frac{[A]^n}{K_{+}^n + [A]^n}\Big)\right] - \gamma_A[H]\Big(\frac{[A]}{K_d + [A]}\Big) \qquad \text{[S104]}$$

$$\frac{d[H]}{dt} = V_{-}\Big(\frac{[A]}{K_{-} + [A]}\Big) - \gamma_H[H] \qquad \text{[S105]}$$

Define non-dimensional concentrations:

$$\bar{A} = \frac{[A]}{K_d} \qquad \text{[S106]}$$

$$\bar{H} = \frac{[H]}{\frac{V_{-}}{\gamma_H}} \qquad \text{[S107]}$$

$$\lambda_{+} = \frac{K_{+}}{K_d} \qquad \text{[S108]}$$

$$\lambda_{-} = \frac{K_{-}}{K_d} \qquad \text{[S109]}$$

Non-dimensionalizing the equations for rate of change of $[A]$ and $[H]$:

For $[A]$:

$$\frac{d\bar{A}}{dt} = \theta_{syn}\left[\frac{V_{stress}}{K_d} + \frac{V_{+}}{K_d}\Big(\frac{\bar{A}^n}{\lambda_{+}^n + \bar{A}^n}\Big)\right] - \frac{\gamma_A V_{-}}{K_d\gamma_H}\bar{H}\Big(\frac{\bar{A}}{1+\bar{A}}\Big) \qquad \text{[S110]}$$

For $[H]$:

$$\frac{d\bar{H}}{dt} = \gamma_H\left[\Big(\frac{\bar{A}}{\lambda_{-} + \bar{A}}\Big) - \bar{H}\right] \qquad \text{[S111]}$$

*Note*: Eqs. S110 and S111 are non-dimensional in concentrations but not in time. The choice of time-scales we have are:

$$\tau_c^1 = \frac{K_d}{\theta_{syn}V_{stress}} \qquad \tau_c^2 = \frac{K_d}{\theta_{syn}V_{+}} \qquad \tau_c^3 = \frac{K_d\gamma_H}{\gamma_A V_{-}} \qquad \tau_c^4 = \frac{1}{\gamma_H}$$

Parameters known above are: $\gamma_H$, $\gamma_A$, $K_d$, and $n$. Since, the choice of time-scales other than $\tau_c^4$ have parameters that are unknown, we choose $\tau_c^4$ as the characteristic time.

$$\bar{t} = \frac{[t]}{\frac{1}{\gamma_H}} \quad \text{[S112]}$$

The fully non-dimensional form of Eqs. S110 and S111 is then:

$$\frac{d\bar{A}}{d\bar{t}} = \frac{\theta_{syn} V_{stress}}{\gamma_H K_d} + \frac{\theta_{syn} V_+}{\gamma_H K_d}\Big(\frac{\bar{A}^n}{\lambda_+^n + \bar{A}^n}\Big) - \frac{V_- \gamma_A}{\gamma_H^2 K_d}\bar{H}\Big(\frac{\bar{A}}{1+\bar{A}}\Big) \quad \text{[S113]}$$

$$\frac{d\bar{H}}{d\bar{t}} = \Big(\frac{\bar{A}}{\lambda_- + \bar{A}}\Big) - \bar{H} \quad \text{[S114]}$$

The non-dimensional formulation reveals that the system's transient and steady-state landscapes are shaped by the three terms in the RHS of Eq. S113. Inspection of these terms show that given the parameters $\gamma_A$, $\gamma_H$, $K_d$ are known, variations in system behavior are directly controlled by three specific biological rate processes: the basal expression of ABA ($V_{stress}$), the positive feedback synthesis rate ($V_+$), and the basal expression rate of enzyme A8H ($V_-$). In the coupled system of Eqs. S113 and S114 above:

- The timescale in the transient response is controlled by $V_-$ in $\frac{V_- \gamma_A}{\gamma_H^2 K_d}$.

- At steady state when RHS=0, the parameters controlling the steady states are then $\lambda_+$, $\lambda_-$, $V_-$ in $\frac{V_- \gamma_A}{\gamma_H^2 K_d}$, $V_{stress}$ in $\frac{V_{stress}}{\gamma_H K_d}$, $V_+$ in $\frac{V_+}{\gamma_H K_d}$, and $\theta_{syn}$.

**B. Response landscape of ABA autoregulation motif.** We first examine the steady-state properties of the system. By setting the time derivatives in Eqs. S113 and S114 to zero, we map the equilibrium concentrations of ABA as a function of the stress input. This analysis reveals how the interplay between the positive feedback strength ($V_+$) and the stress-dependent synthesis rate ($V_{stress}$) determines the number and stability of the system's steady states (bifurcation).

In Fig. S6b, we plot the steady-state ABA concentration as a function of $V_{stress}$ for varying strengths of positive feedback ($V_+$), with $V_-$ held fixed. When the positive feedback gain is low (low values of $V_+$; blue curves in Fig. S6b), the concentration of ABA grows monotonically with increasing $V_{stress}$, such that a unique value of $[ABA]_g$ exists for any given stress level. However, as $V_+$ increases (yellow and red curves in Fig. S6b), the system undergoes a bifurcation, giving rise to an S-shaped curve characteristic of bi-stability. In this regime, the system can support three equilibrium solutions over a specific range of stress inputs: two stable states (low and high ABA) separated by an unstable intermediate state (Fig. S6c). This bi-stability implies the potential for hysteresis, where the current state of the system depends on its history.

To comprehensively map the global behavior of the motif, we performed a two-parameter stability analysis in the ($V_+$, $V_{stress}$) plane (Fig. S6d) for a fixed value of $V_-$. This phase diagram classifies the system's dynamic response into four distinct regimes: no oscillations, damped oscillations, sustained oscillations, and unstable state. This steady-state landscape provides critical guidance for parameter estimation with respect to the prediction of damped or sustained oscillations. The black arrows in Figs. S6c and S6d show how the parameter ranges explored using the full model in Fig. 2 and 3 ("Merilo parameters") and in Fig. 4 ("Hysteresis parameters") map onto the predicted dynamical regimes of the governing equations of ABA autoregulation (Eqs. S113 and S114).

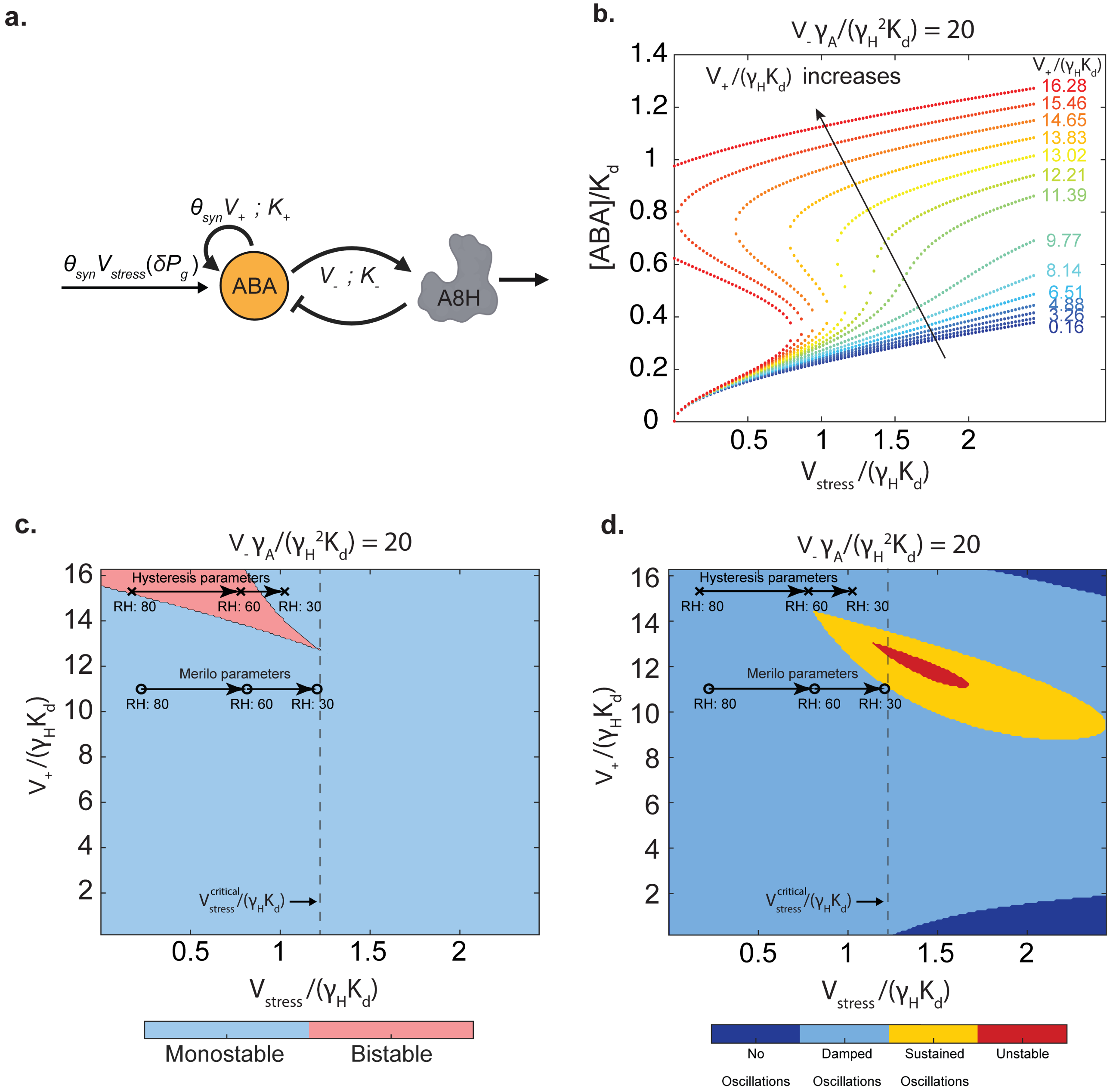

**Fig. S6. Bifurcation and stability analysis of the non-dimensionalized ABA autoregulation model. a.** Schematic of the ABA autoregulation motif governed by Eqs. S113 and S114. **b.** Non-dimensional steady-state ABA concentration ($[ABA]/K_d$) as a function of normalized stress ($V_{stress}/(\gamma_H K_d)$). The curves illustrate the steady-state response to increasing dimensionless feedback gain $V_+/(\gamma_H K_d)$ (arrow indicates increasing values, colored from blue to red). Low positive feedback ($V_+/(\gamma_H K_d)$) yields a single stable state (monotonic curves), while high positive feedback generates S-shaped curves indicative of bi-stability. All curves are plotted with the fixed parameter group $V_-\gamma_A/(\gamma_H^2 K_d) = 20$. **c.** Stability phase diagram in the feedback-stress parameter space. Shaded regions classify steady-state solutions as monostable (blue) or bistable (pink). Overlaid arrows trace the physiological operating points for two distinct parameter sets – "Merilo parameters" and "Hysteresis parameters" as relative humidity (RH) decreases from 80% to 30% in Fig. 2 main text. **d.** Classification of dynamic behaviors based on linear stability analysis. Colored regions indicate parameter combinations yielding no oscillations (dark blue), damped oscillations (light blue), sustained oscillations (yellow), or unstable diverging responses (red). The overlaid trajectories demonstrate that while the range of parameters explored in Fig. 4 with hysteresis ("Hysteresis parameters") remains in the damped oscillation region, the range of parameters explored in Figs. 2 and 3 that match the dynamics reported by Merilo[62] ("Merilo parameters") traverses the oscillatory regime under severe stress (RH 30%). The vertical dashed lines in (c) and (d) are the critical stress values ($V_{stress}^{critical}$) beyond which oscillations are induced in the system defined by "Merilo parameters". The governing equations are provided in Section S5.

**C. Role of $V_{stress}$ in inducing oscillations.** We now numerically analyzed the coupled system of ODEs (Eqs. S113 and S114 - Fig. S7a) as a function of the stress input rate, $V_{stress}$. Fig. S7b shows an increase in $V_{stress}/(\gamma_H K_d)$ with increasing water demand stress (VPD). An increase in VPD (external trigger) acts as an input for increasing $V_{stress}/(\gamma_H K_d)$ (internal trigger) that leads to ABA biosynthesis. Fig. S7c shows the transient dynamics of ABA concentration in response to imposed $V_{stress}$ levels at $t = 0$ (colorbar Fig. S7c), with damped oscillations emerging at lower $V_{stress}/(\gamma_H K_d)$ and sustained oscillations appearing at higher input levels.

The amplitude of oscillations as a function of $V_{stress}/(\gamma_H K_d)$ is presented in Fig. S7d. Sustained oscillations arise when $V_{stress}/(\gamma_H K_d)$ exceeds a critical threshold of approximately $V_{stress}^{critical}/(\gamma_H K_d) \approx 1.22$ (vertical dashed line in Figs. S6e-d). As shown in Fig. S7e, the period of sustained oscillations decreases with increasing $V_{stress}/(\gamma_H K_d)$, indicating a corresponding increase in oscillation frequency.

Oscillatory behavior often reflects the underlying stability characteristics of a dynamical system. To assess system stability, we computed the eigenvalues of the Jacobian matrix evaluated at steady state (Fig. S7f). For $V_{stress} < V_{stress}^{critical}$, the eigenvalues possess negative real parts and complex conjugate imaginary components, indicative of damped oscillations. However, for $V_{stress} > V_{stress}^{critical}$, the real parts become positive, suggesting local instability and a propensity for growing oscillations.

Despite the prediction of unstable dynamics by linear stability analysis, the system exhibits sustained oscillations rather than divergent behavior. This phenomenon is explained by the emergence of a stable limit cycle, as illustrated in the phase portrait (Fig. S7g). For high $V_{stress}/(\gamma_H K_d)$ (e.g., 1.63), trajectories spiral outward from the unstable fixed point but become confined within a trapping zone, forming a closed-loop (blue). In contrast, for lower $V_{stress}/(\gamma_H K_d)$ (e.g., 0.48), during damped oscillations, no such loop forms and the trajectory converges to a stable point (red). Near the equilibrium, positive feedback induces instability and outward spiraling, while further from equilibrium, degradation terms exert dominant negative feedback, redirecting trajectories inward. The dynamic balance between destabilizing feedback and restorative damping results in a self-sustained oscillation, a hallmark of a limit cycle.

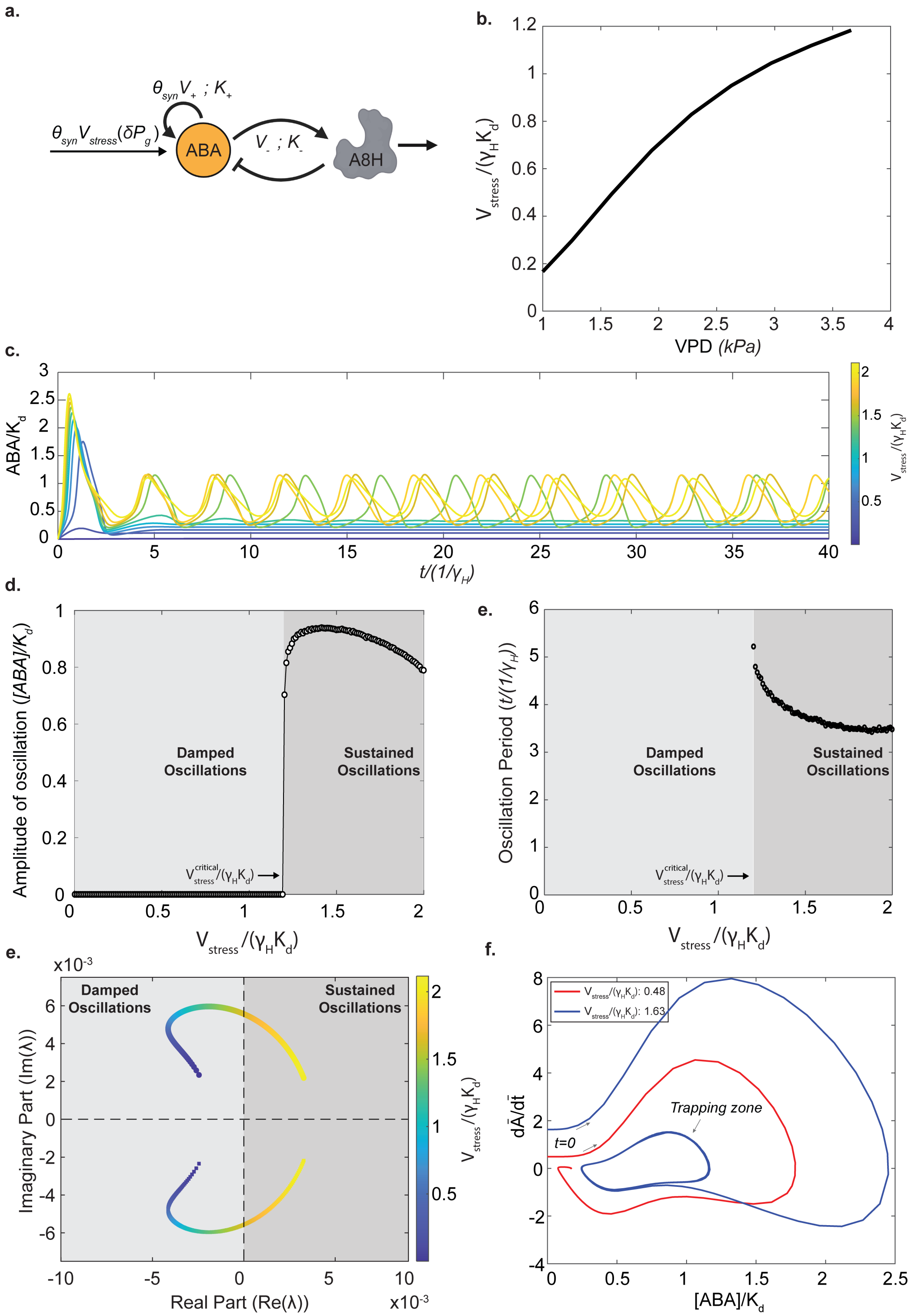

**Fig. S7. Dynamic stability and oscillatory behavior of the non-dimensionalized ABA regulation motif.** This figure characterizes the transition from stability to oscillation as a function of the normalized stress-induced synthesis rate, $V_{stress}/(\gamma_H K_d)$. **a.** Schematic of the ABA autoregulation motif governed by Eqs. S113 and S114 **b.** Relationship between Vapor Pressure Deficit (VPD) and the non-dimensional synthesis rate $V_{stress}/(\gamma_H K_d)$, used to map environmental conditions to model inputs. **c.** Time evolution of normalized guard cell ABA concentration ($[ABA]/K_d$) at imposed stress levels $V_{stress}$ at $t = 0$. Colors correspond to imposed $V_{stress}/(\gamma_H K_d)$ values (see colorbar), response trajectories showing a transition from damped oscillations to sustained oscillations. **d.** Bifurcation diagram showing the oscillation amplitude of $[ABA]/K_d$. The system exhibits a bifurcation at a critical stress threshold ($V^{critical}_{stress}$), marking the onset of sustained oscillations. **e.** Oscillation period ($t \cdot \gamma_H$) as a function of stress intensity, plotted for the region of sustained oscillations. **f.** Trajectory of the Jacobian eigenvalues in the complex plane as stress increases. The crossing of the eigenvalue pair from the left (stable, negative real part) to the right half-plane (unstable, positive real part) confirms the bifurcation mechanism. Colors indicate $V_{stress}$ magnitude. **g.** Phase plane portraits of normalized ABA concentration versus its time derivative ($d\bar{A}/d\bar{t}$). The red trajectory (low stress) spirals into a stable fixed point, while the blue trajectory (high stress) converges to a stable limit cycle (sustained oscillation) surrounding an unstable fixed point. The region containing the limit cycle is marked as a trapping zone. The governing equations are provided in Section S5.

**D. Role of $V_-$ in controlling stomatal oscillations.** A key feature of stomatal response to humidity changes is the transient overshoot in stomatal conductance, characterized by damped oscillations, as observed in Fig. 2 of the main text. Such oscillatory behavior has also been documented in gas exchange measurements during relative humidity step changes in prior studies[12,50,63]. However, Assmann[64] reported a monotonic response in stomatal conductance following a wrong-way response, with no oscillations observed under similar RH step changes. Furthermore, some studies have also shown stomatal oscillations[4,65].

In this section, we explore in the context of the full HP-HA model (Fig. S8a), how the ABA regulatory motif produces distinct dynamic responses depending on the magnitude of the parameter $V_-$, while keeping all other parameters $K_+$, $K_-$, $V_{stress}$, $V_+, \theta_{syn}$ constant (Fig. S8a.). Fig. S8 illustrates the dynamics for VPD step changes from 1.1 kPa to 2.2 kPa (Fig. S8b.–c.) and from 1.1 kPa to 3.9 kPa (Fig. S8d.–e.), using the same parameter values $K_+ = K_- = 1560\ nM$, $V_{stress} = 8.85\ nM/s$, $V_+ = 67.6\ nM/s$ as in Fig. 2 of the main text (Table S1). In Fig. S8b, the stomatal and ABA responses for wildtype ($V_- = 1.5$) show damped oscillations. By contrast, Fig. S8c shows that reducing $V_-$ by 10-fold results in a monotonic transition to the new steady state, with oscillations eliminated. For a stronger VPD perturbation, which induces sustained oscillations, the oscillation frequency is modulated by $V_-$. A decrease in $V_-$ from 2.5 nM/s to 1.5 nM/s leads to a lower oscillation frequency, increasing the period $T$ from 19 minutes (Fig. S8d) to 24 minutes (Fig. S8e). Further, we show in Fig. S8f that the period of oscillation decreases with increasing $V_-$.

$$\Omega = \frac{V_-/K_d}{\gamma_H(\frac{\gamma_H}{\gamma_A})} = \frac{f_{INT}}{f_{SP}} = \frac{\text{Species interaction rate}}{\text{Individual species turnover rate}} \qquad [S115]$$

$\Omega$, a dimensionless ratio (Eq. S115) reflecting the balance between species interaction and individual species turnover rates and hence controlling the timescale of ABA dynamics. Specifically, $V_-/K_d$ quantifies how efficiently enzyme $H$ converts substrate $A$ into product and is a standard metric in enzyme-substrate kinetics. A high $V_-/K_d$ indicates either a high catalytic rate $V_-$ or strong substrate affinity (low $K_d$), implying that enzyme $H$ can effectively catalyze $A$ even at low concentrations. The individual species turnover rate is governed by the system's characteristic timescale for species evolution ($\gamma_H$) and the relative rate of change between $H$ and $A$ ($\gamma_H/\gamma_A$).

When $\Omega$ becomes large, the species interaction frequency ($f_{INT}$) exceeds the species-specific decay rate ($f_{SP}$). In this regime, enzyme $H$ exhibits high affinity for $A$, resulting in strong feedback that leads to overshoots in ABA concentration and producing either damped or sustained oscillations. Conversely, when $\Omega$ becomes small, $f_{INT} < f_{SP}$; enzyme $H$ has weak affinity for $A$, resulting in minimal feedback. In this case, ABA levels rise monotonically to a new steady state without overshoot or oscillations, as $H$ fails to significantly modulate $A$ due to low catalytic efficiency relative to its own degradation.

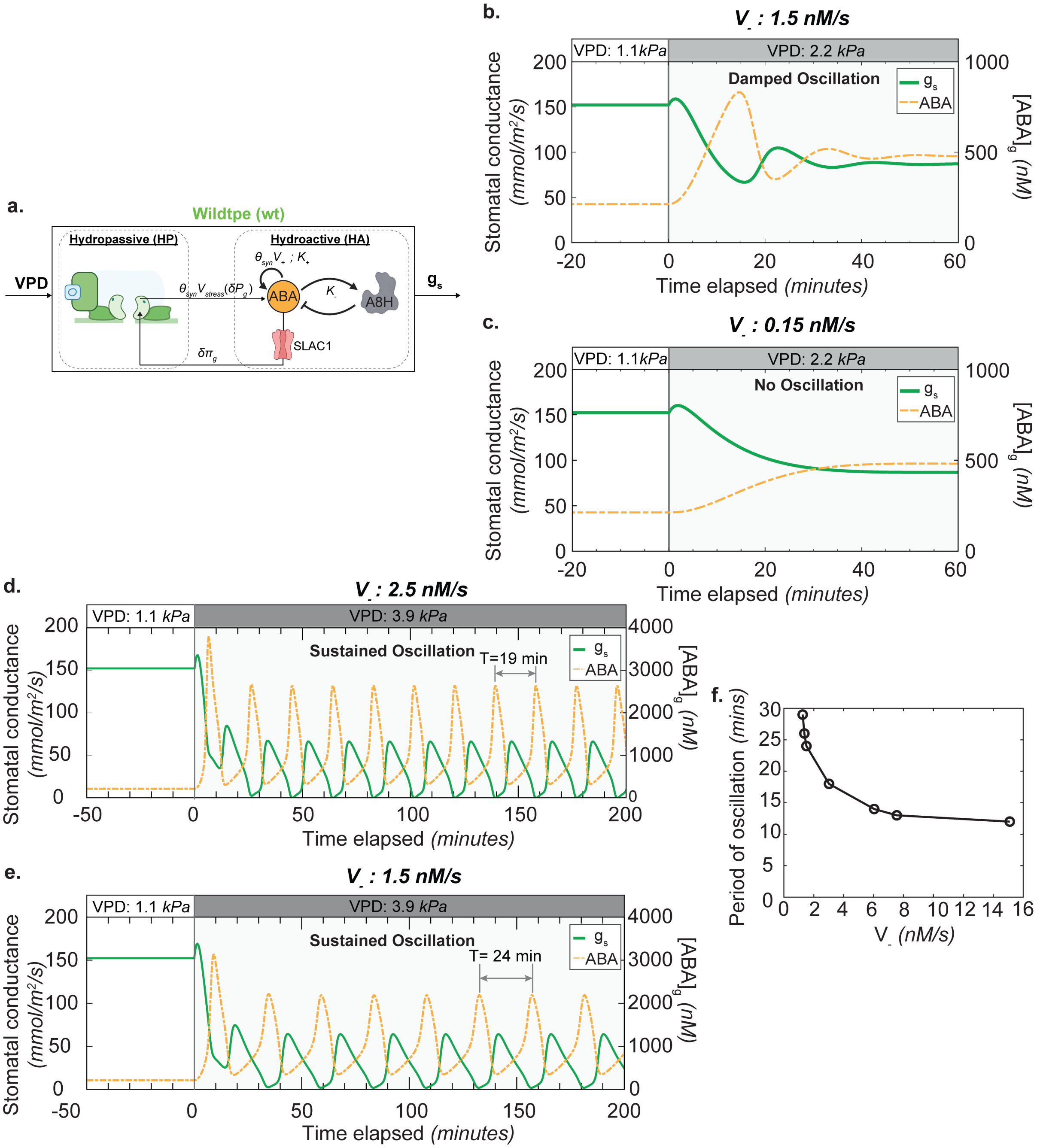

**Fig. S8. Dependence of stomatal oscillation characteristics on upregulation rate of 8'-hydroxylase (A8H), $V_-$. a.** Schematic of the coupled model for wildtype. The schematic shows the hydropassive (HP) and hydroactive (HA) components as defined in Fig. 1 of main text. The HA component features the intact ABA autoregulation motif driving SLAC1 activation that leads to osmotic adjustment and stomatal regulation. Trajectories plotted for parameter values: $K_+ = K_- = 1560\ nM$, $V_{stress} = 8.85\ nM/s$, $V_+ = 67.6\ nM/s$ with $\theta_{syn} = 1$ for wildtype with variable $V_-$. **b.–c.** Stomatal responses to a moderate perturbation in $VPD$ from 1.1 kPa to 2.2 kPa. In both cases, the initial stomatal conductance exhibits a "wrong-way" transient before transitioning to the "right-way" response toward a new steady state. In (**b**), at a higher feedback strength ($V_- = 1.5\ nM/s$), ABA dynamics overshoot due to stronger A8H feedback, leading to damped oscillations in both ABA concentration and stomatal conductance. In (**c**), at a lower feedback strength ($V_- = 0.15\ nM/s$), the system exhibits a smooth monotonic convergence to steady state with no oscillations. This occurs because A8H undergoes rapid enzymatic degradation, weakening its feedback on ABA levels and diminishing the oscillatory behavior. **d.-e.** Stomatal responses to a larger perturbation in $VPD$ from 1.1 kPa to 3.9 kPa. This induced a stronger perturbation in ABA synthesis at $t = 0\ minutes$. In both cases, the "wrong-way" initial response is followed by sustained oscillations in stomatal conductance. In (**d**), ($V_- = 2.5\ nM/s$) yields a sustained oscillatory response with a period of $T = 19\ minutes$. In (**e**), at a lower feedback strength ($V_- = 1.5\ nM/s$), the system exhibits faster oscillations with an increased period of $T = 24\ minutes$, reflecting weaker and slower upregulation of A8H activity. As $V_-$ increases, the frequency of sustained oscillations rises. **f.** Time period of oscillations as a function of $V_-$ feedback strength shows a decline in time period of oscillation with increasing strength of negative feedback. The governing equations for this circuit are provided in Sections S2 and S3.

**E. Parameter fitting for the ABA regulation motif.** The parameters unknown in the ABA regulation motif in Table S6 are: $V_{stress}$, $V_+$, $V_-$, $K_+$, $K_-$ and $\theta_{syn}$.To fully characterize our system, we needed to determine six unknown parameters. We achieved this by matching our model outputs to six corresponding experimental measurements of stomatal conductance change to relative humidity. This established a well-defined system with an equal number of unknowns and constraints. The non-dimensionalized system of ABA governing equations was used to facilitate the analysis of the steady-state and dynamic properties of stomatal conductance. To calibrate our model against experimental data from Merilo[62], we employed a sequential parameter fitting strategy as follows:

1. **Wild-type parameterization**: Initially, we focused on fitting parameters for the wild-type plant, beginning with a fixed value of $\theta_{syn} = 1$.

2. **Characteristic concentration ratio**: We assumed that the ratio of characteristic concentrations for ABA positive feedback ($K_+$) and negative feedback ($K_-$), relative to the ABA degradation Michaelis-Menten constant ($K_d$), is of order one $O(1)$. With $\lambda_+ = \lambda_- = 0.78$, we have $K_+ = K_- = 0.78 * K_d$.

3. **$V_-$ fitting**: Recognizing that the ratios $V_{stress}/V_-$ and $V_+/V_-$ are needed to solve for the steady states, we first constrained $V_-$. We adjusted $V_-$ to achieve the closest possible match to the quantitative and qualitative features of the wild-type response dynamics for relative change in stomatal conductance, specifically targeting the characteristic timescale (valley in stomatal conductance overshoot).

4. **$V_{stress}$ and $V_+$ fitting**: With $V_-$ for dynamics/transients established, we proceeded to constrain the steady-state behavior. This required determining the remaining two parameters, $V_{stress}$ and $V_+$. By solving for the steady states, we generated plots of steady-state ABA levels as a function of ($V_{stress}$, $V_+$). From this plot we identified the combinations that lead to a single stable solution (monostability) and two stable solutions (bi-stability) leading to hysteresis (Fig. S6b and S6c).

   Furthermore, we restricted our zone of interest for parameters that led to damped oscillations as observed in the wildtype data (light blue zone in Fig. S6d). While the bistable region could produce damped oscillations, we could not fit the steady states. Hence, the parameters for wildtype were sampled from the monostable region that led to damped oscillations.

5. **ABA synthesis mutant fitting**: To model the ABA synthesis mutant, we maintained all parameters from above at their wildtype values except for $\theta_{syn}$, which we varied between zero and one to match the experimental steady states.

The dimensional fitted parameters that capture the stomatal responses reported by Merilo[62] and used in Fig. 2 of the main text are:

**Table S1. Fitted ABA regulation motif parameters (dimensional)**

| Parameter | Value | Units |
|---|---|---|
| $K_+$ | 1560 | nM |
| $K_-$ | 1560 | nM |
| $V_{stress}$ | 8.85 | nM/s |
| $V_+$ | 67.6 | nM/s |
| $V_-$ | 1.5 | nM/s |

Fig. S9 presents an additional exploration to demonstrate the appropriateness of the chosen parameters in Table S1 (Fig. S9). Shown are steady-state stomatal response to relative humidity and the speed of stomatal closure defined as the time interval between the peak of the wrong-way response (WWR) and the overshoot valley compared against experimental data from Merilo[62]. This parameter set consistently predict a higher steady-state stomatal conductance ($g_s$) for the *syn* mutant (*aao3-2*) compared to the wildtype (WT) across all tested relative humidity (RH) levels (Fig. S9a). These predictions align well with experimental data[62], correctly capturing the decrease in $g_s$ as RH drops from 80% (low VPD, 1.1 kPa) to 60% (high VPD, 2.2 kPa). Crucially, the model demonstrates that $g_s$ decreases monotonically as guard cell ABA concentration ($[\text{ABA}]_g$) increases (inset, Fig. S9a), confirming that the physiological divergence between the genotypes stems from lower intrinsic ABA production in the mutant. Furthermore, with this parameter set predicts that the speed of stomatal closure is indistinguishable between the WT and the *syn* mutant. This mirrors the experimental finding[62] that the difference in closure speed is not statistically significant (*n.s.*).

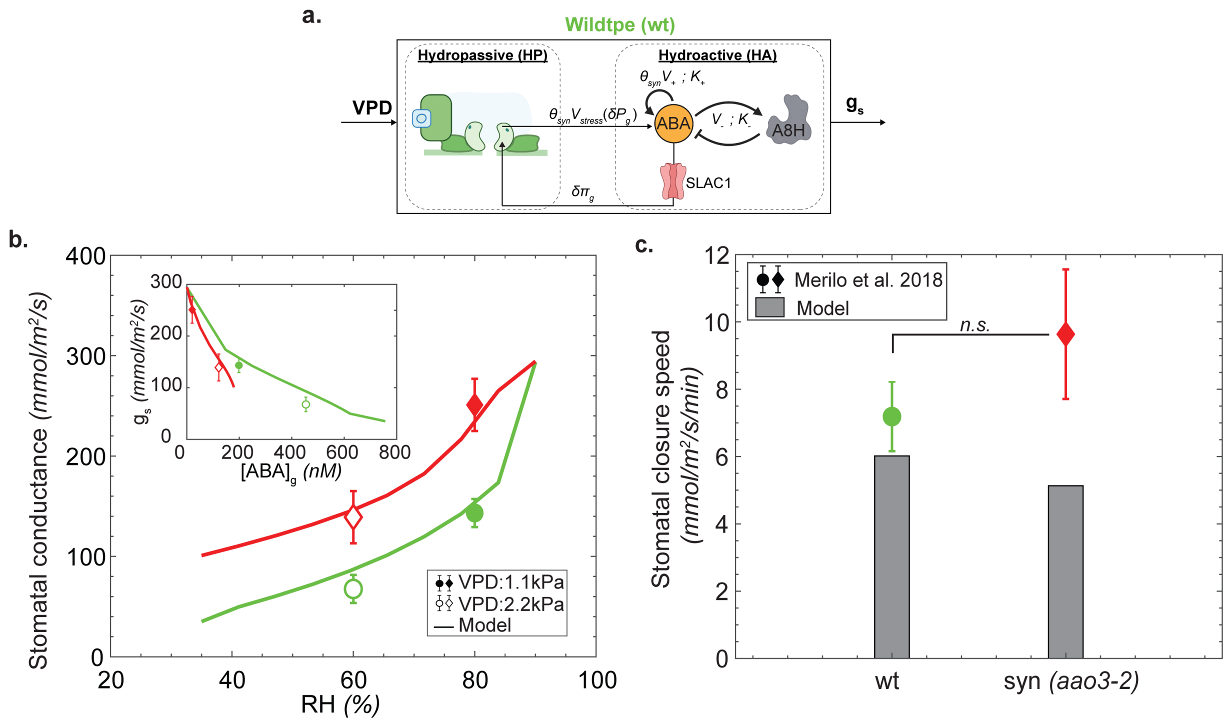

**Fig. S9. Model responses for relative humidity (RH) change - staeady-state and stomatal closure speed: a.** Schematic of the coupled model for wildtype. The schematic shows the hydropassive (HP) and hydroactive (HA) components as defined in Fig. 1 of main text. The HA component features the intact ABA autoregulation motif driving SLAC1 activation that leads to osmotic adjustment and stomatal regulation. **b.** Steady-state stomatal conductance as a function of relative humidity (RH). Solid lines represent the model predictions, while symbols show experimental data from Merilo[62] for wildtype (wt; circles) and the ABA synthesis mutant syn *(aao3-2)* (diamonds). Filled symbols correspond to 80% RH (VPD: 1.1 kPa), and open symbols to 60% RH (VPD: 2.2 kPa). The model successfully captures the higher stomatal conductance observed in syn mutant. (**Inset b.**) Model-predicted relationship steady state stomatal conductance and guard cell ABA concentration ($[ABA]_g$) for wildtype (green line) and the syn mutant (red line). Stomatal conductance decreases monotonically as $[ABA]_g$ increases. **c.** Stomatal closure speed. Grey bars represent experimental data from Merilo while colored diamonds show the model's steady state predictions for wildtype (green) and the syn mutant (red). Error bars represent standard deviation. *n.s.* = not significant. Parameters for ABA regulation motif used are: $K_+ = K_- = 1560\,nM$, $V_{stress} = 8.85\,nM/s$, $V_+ = 67.6\,nM/s$, $V_- = 1.5\,nM/s$ with $\theta_{syn} = 1$ for wildtype and $\theta_{syn} = 0.12$ for synthesis mutant (Table S1)

The parameters capturing the transient stomatal conductance response for wildtype, ABA synthesis mutant, and ABA signaling mutants (Fig. 2) are provided in Table S1 in sub-section S5E. These parameter sets were selected because they capture the transient and steady-state gas exchange responses of different *Arabidopsis thaliana* genotypes, including the emergence of oscillations in stomatal conductance following a large perturbation in relative humidity.

**F. More than one set of parameters can explain the observed stomatal conductance dynamics.** In Fig. S10, we show a different set of parameters corresponding to Table S2 that can also capture the steady-state and transient responses in stomatal conductance to step change in relative humidity from Merilo[62]. However, this set of parameters is not able to capture the emergence of oscillations upon larger step changes in relative humidity.

#### Table S2. Second set of ABA regulation motif parameters (dimensional)

| Parameter | Value | Unit |
|---|---|---|
| $K_+$ | 1000 | nM |
| $K_-$ | 1000 | nM |
| $V_{stress}$ | 9.83 | nM/s |
| $V_+$ | 16.3 | nM/s |
| $V_-$ | 1 | nM/s |

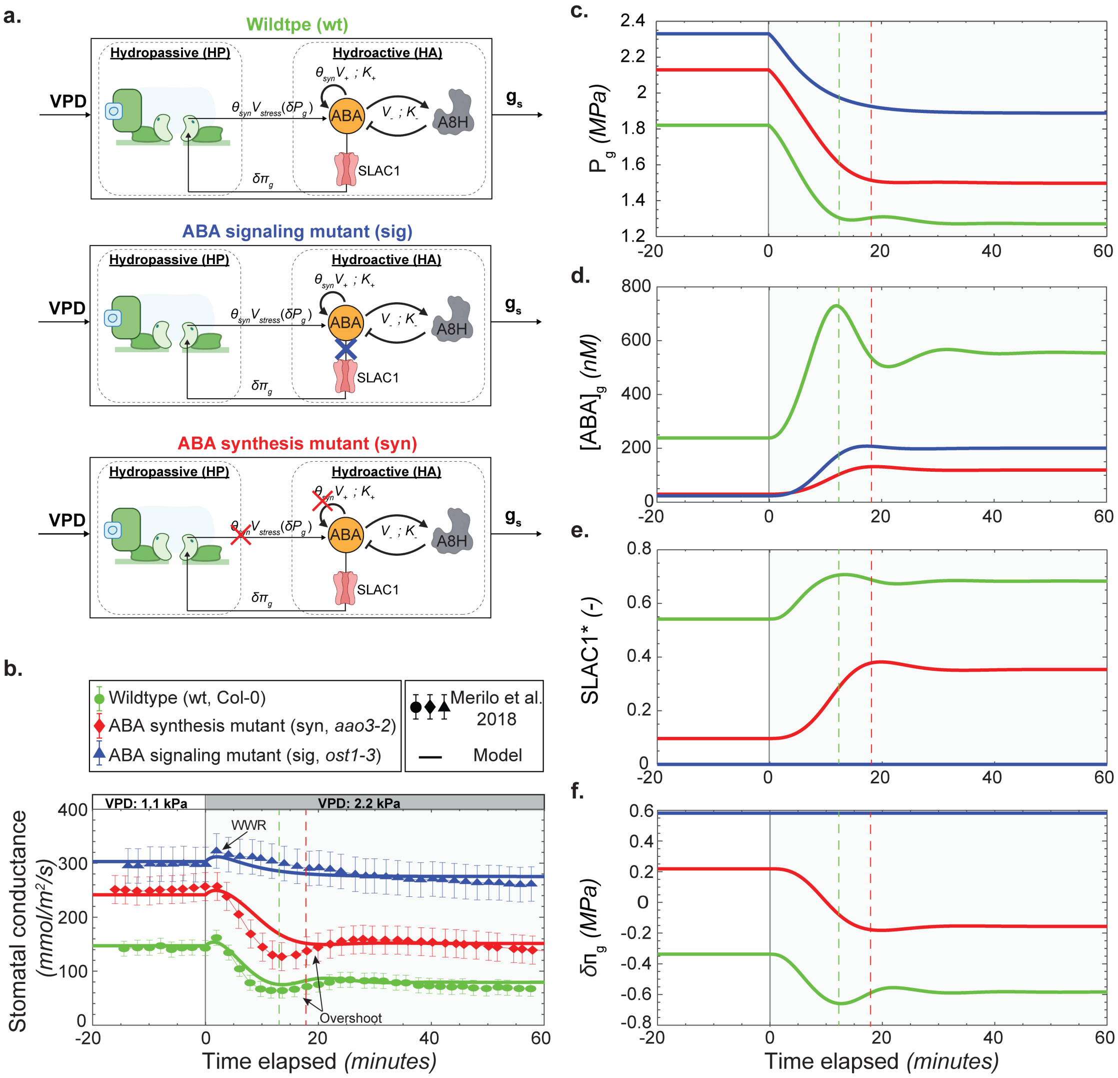

**Fig. S10. Transient stomatal responses to changes in vapor pressure deficit (VPD) across genetic variants for non-unique parameter set. a.** Schematics of the coupled model variants applied to Wildtype (wt), ABA signaling mutant (sig), and ABA synthesis mutant (syn). The schematic shows the hydropassive (HP) and hydroactive (HA) components as defined in Fig. 1. In wt, the HA component features the intact ABA autoregulation motif driving SLAC1 activation. In sig, the pathway is disrupted downstream of ABA autoregulation but upstream of channel activation (blue cross), representing diminished OST1 function. In syn, the ABA production is attenuated (red cross) via a reduction in $\theta_{syn}$. **b.** Predicted and measured transients for a step change in vapor pressure deficit (VPD) from 1.1 kPa to 2.2 kPa for stomatal conductance, $g_s$. The trajectories (solid curves) and data (symbols in (**b**)) are for representations of wildtype (wt, Col-0 – green curves and circles), an ABA synthesis mutant in this same background (syn, *aao3-2* – red curves and diamonds), and an ABA signaling mutant (sig, *ost1-3* – blue curves and triangles). Data are from Merilo [62] for these genetic variants and this same perturbation in VPD. **c.-f.** Predicted transients corresponding to those in (**b**) for: guard cell turgor, $P_g$ (**c**); concentration of ABA in guard cells, $[ABA]_g$ (**d**); fractional activation of anion channel, SLAC1* (**e**); and osmotic potential change in guard cells, $\delta\pi_g$ (**f**). This step in VPD corresponds to a relative humidity (RH) change from 80% to 60%, at 36°C. Parameters for the ABA motif were: $K_+ = K_- = 1000$ nM, $V_{stress} = 9.83$ nM/s, $V_+ = 16.3$ nM/s, $V_- = 1$ nM/s with $\theta_{syn} = 1$ for wildtype and $\theta_{syn} = 0.12$ for synthesis mutant (Table S2). See main text and Section S3 for details on model and representations of genetic variants.

**G. Understanding the emergence of oscillations in coupled HP-HA model.** To investigate the origin of oscillations in stomatal dynamics, we performed a combined theoretical and numerical analysis of feedback mechanisms in our multiscale model.

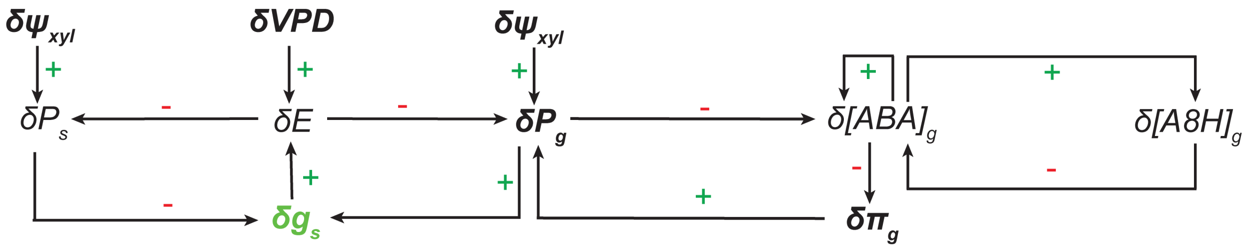

**Fig. S11. Feedback loops in coupled hydropassive-hydroactive (HP-HA) model.** The turgor pressure, $P_g$ [MPa], osmotic potential, $\pi_g$ [MPa], and ABA concentrations, $[ABA]_g$ [nM] regulate stomatal aperture through a set of coupled feedback loops. HP processes couple changes in water availability ($\delta\psi_{xyl}$ [MPa]) and demand ($\delta VPD$ [kPa]) to changes in turgor pressure ($\delta P_g$). These changes trigger the HA pathway with changes in the concentration of ABA ($\delta[ABA]_g$) and downstream changes in osmotic potential ($\delta\pi_g$) that feed back to change guard cell turgor and stomatal conductance ($\delta P_g$). The HA process includes two internal feedback loops that capture the Autoregulatory (AR) dynamics of ABA: positive feedback of ABA on its own synthesis (above $\delta[ABA]_g$) and negative feedback through its degradation, mediated by A8H (right $\delta[ABA]_g$).

A positive feedback occurs when "more of A leads to more of B" or "less of A leads to less of B", while a negative feedback occurs when "more of A leads to less of B" or "less of A leads to more of B". A summary of the feedback motifs represented in the circuit diagram (Fig. S11 in the main text; also shown in Fig. S12a, S13a) is outlined below:

**Table S3. Feedbacks in multiscale model**

| Parameter | Description | Value |
|---|---|---|
| **Hydropassive framework** | | |
| More of $VPD$ leads to more of $E$ | Positive feedback | $+$ |
| More of $E$ leads to less of $P_g$ | Negative feedback | $-$ |
| Less of $P_g$ leads to less of $g_s$ | Positive feedback | $+$ |
| Less of $g_s$ leads to less of $E$ | Positive feedback | $+$ |
| More of $E$ leads to less of $P_s$ | Negative feedback | $-$ |
| Less of $P_s$ leads to more of $g_s$ | Negative feedback | $-$ |
| **Hydroactive framework** | | |
| Less of $P_g$ leads to more of $[ABA]_g$ | Negative feedback | $-$ |
| More of $[ABA]_g$ leads to less of $\pi_g$ | Negative feedback | $-$ |
| Less of $\pi_g$ leads to less of $P_g$ | Positive feedback | $+$ |
| Less of $P_g$ leads to less of $g_s$ | Positive feedback | $+$ |
| **ABA regulation motif** | | |
| More of $[ABA]_g$ leads to more of $[ABA]_g$ | Positive feedback | $+$ |
| More of $[ABA]_g$ leads to more of $[A8H]_g$ | Positive feedback | $+$ |
| More of $[A8H]_g$ leads to less of $[ABA]_g$ | Negative feedback | $-$ |

To quantify the influence of these feedback loops, we evaluated their effects on system dynamics using different configurations shown in Fig. S12 and S13. Each figure contains four feedback scenarios (panels a–d), with (i) depicting the input signal and (ii–iv) showing outputs for guard cell ABA concentration, guard cell osmotic potential change, guard cell and subsidiary cell turgor pressures, and stomatal conductance, respectively. In Fig. S12:

- (a) represents the full wild-type system subjected to a step drop in VPD from 1.1 kPa to 2.2 kPa. The inset in panel (a-i) shows the resulting time profile of stress-induced ABA synthesis $V_{stress}$.

- (b) simulates a hydropassive-only system by removing ABA's effect on osmotic potential, analogous to an *ost1* or *slac1* mutant, while keeping the same VPD step change as in (a).
- (c) VPD constant at 1.1 kPa and a step change in osmotic potential, equivalent in magnitude to the wild-type response in (a-iii).
- (d) this case removes hydromechanical triggering of ABA synthesis and directly imposes a step change in $V_{stress}$ while holding VPD constant to trigger a change in $[ABA]_g$.

Fig. S13 follows an identical configurations as Fig. S12, except the VPD step change in panels (a) and (b) is from 1.1 kPa to 3.9 kPa, and in (c) and (d), VPD is pinned at 3.9 kPa with step changes scaled accordingly to match panel (a). Key findings are as follows:

- In the full wildtype system (panel a), we observe damped oscillations in Fig. S12 and sustained oscillations in Fig. S13 across all state variables, reflecting the influence of magnitude of VPD perturbation (same as main text Fig. 3).
- In the hydropassive-only case (b), ABA exhibits an overshoot and relaxes to a steady state, with a larger overshoot in Fig. S13 due to the greater VPD drop. However, this change does not propagate to osmotic potential change, turgor pressure, or stomatal conductance. This indicates that hydraulic responses alone, without ABA-induced osmotic feedback, do not produce oscillations.
- In panel (c), direct perturbation of osmotic potential causes marginal ABA accumulation and step-like changes in turgor and conductance, again failing to trigger oscillations.
- In panel (d), where upstream feedback from turgor to ABA is removed but downstream ABA feedback remains intact, we recover the full dynamic range of ABA. Both Fig. S12 and Fig. S13 exhibit oscillations—damped and sustained, respectively, mirroring the wildtype. This isolates the hydroactive contribution from ABA regulation and reveals that oscillations are driven by the ABA regulatory loop, arising from the nonlinear nature of ABA autoregulation motif.

Taken together, these results clearly demonstrate that oscillatory behavior in stomatal dynamics depends on the ABA feedback circuitry, particularly the nonlinear interactions within the ABA regulatory motif, as described by Eq. S104 and S105.

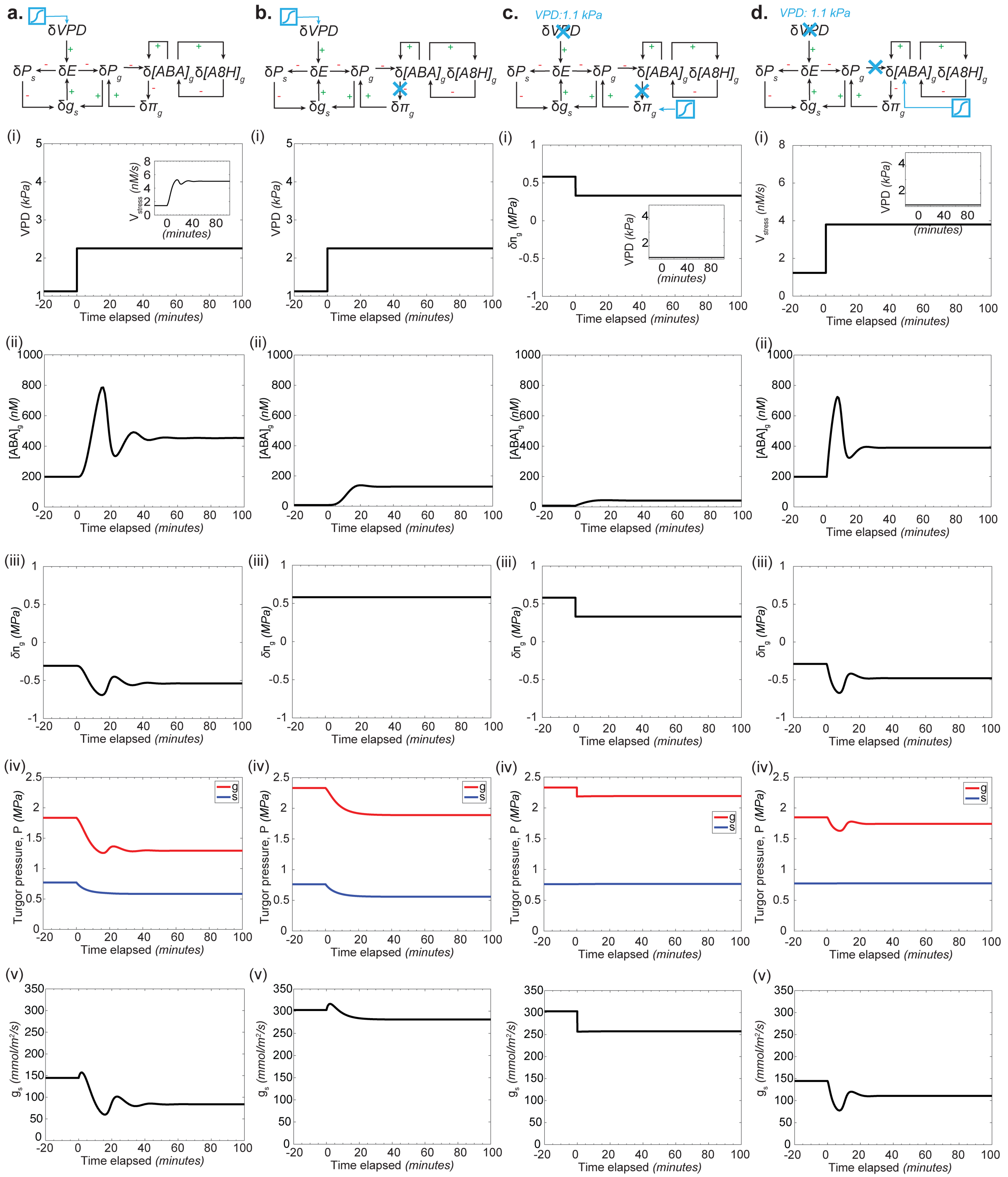

**Fig. S12. VPD, ABA, osmotic pressure, turgor, and stomatal conductance responses under different feedback configurations.** Panels **a.–d.** show model responses under four distinct feedback motifs. For each case, we present: (i) imposed variable change: RH for (**a**) and (**b**) and $\delta\pi_g$ and $V_{stress}$ for (**c**) and (**d**) respectively, (ii) guard cell ABA concentration $[ABA]g$, (iii) guard cell osmotic potential change $(\delta\pi_g)$ , (iv) turgor pressure, and (v) stomatal conductance over time. In panels (**a**) and (**b**), the system is perturbed by a step change in RH from 80% to 60%. In panels (**c**) and (**d**), RH is held constant at 60%, and equivalent perturbations are applied instead to $\delta\pi_g$ (panel **c**) and $V_{stress}$ (panel **d**), mirroring the effects of RH change in panel (**a**).

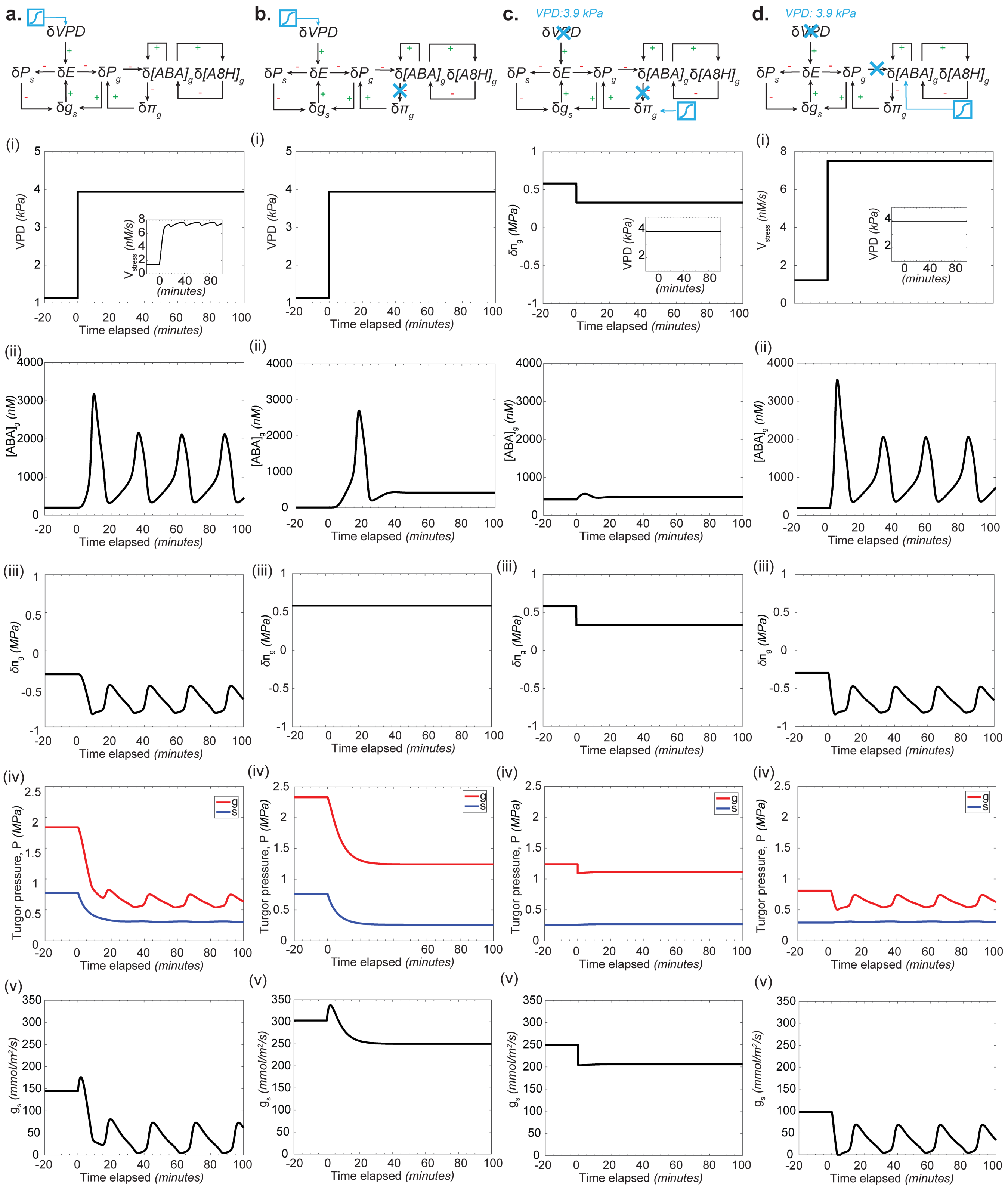

**Fig. S13. VPD, ABA, osmotic pressure, turgor, and stomatal conductance responses under different feedback configurations.** Panels **a.–d.** show model responses under four distinct feedback motifs. For each case, we present: (i) imposed variable change: RH for (**a**) and (**b**) and $\delta\pi_g$ and $V_{stress}$ for (**c**) and (**d**) respectively, (ii) guard cell ABA concentration $[ABA]g$, (iii) guard cell osmotic potential change $(\delta\pi_g)$ , (iv) turgor pressure, and (v) stomatal conductance over time. In panels (**a**) and (**b**), the system is perturbed by a step change in RH from 80% to 30%. In panels (**c**) and (**d**), RH is held constant at 30%, and equivalent perturbations are applied instead to $\delta\pi_g$ (panel **c**) and $V_{stress}$ (panel **d**), mirroring the effects of RH change in panel (**a**).

### S6. Comparing model results with experimental data of oscillations and hysteresis

In this section, we present comparison of experiments with the predictions from our model.

**A. Comparing model and experiments on oscillations.** In their seminal 1974 study, Farquhar and Cowan[66] conducted gas exchange experiments on cotton and observed both damped and growing oscillations in stomatal conductance and evaporation. Our model captures these dynamic features. As shown in Fig. S14, following an initial step decrease in relative humidity (Fig. S14c), Farquhar and Cowan reported damped oscillations in stomatal conductance (Fig. S14a) and evaporation (Fig. S14b). Upon a second RH perturbation, they observed oscillations that became unstable and increased in magnitude over time.

Our model reproduces these responses: a first RH step elicits damped oscillations, while a subsequent step induces growing oscillations in both stomatal conductance and evaporation (Fig. S14d–f). Notably, while Farquhar and Cowan documented the onset of instability, the experimental time window may have been insufficient to capture the full development of a limit cycle. Our model supports this interpretation, predicting a transition from instability to sustained oscillations with continued stimulation.

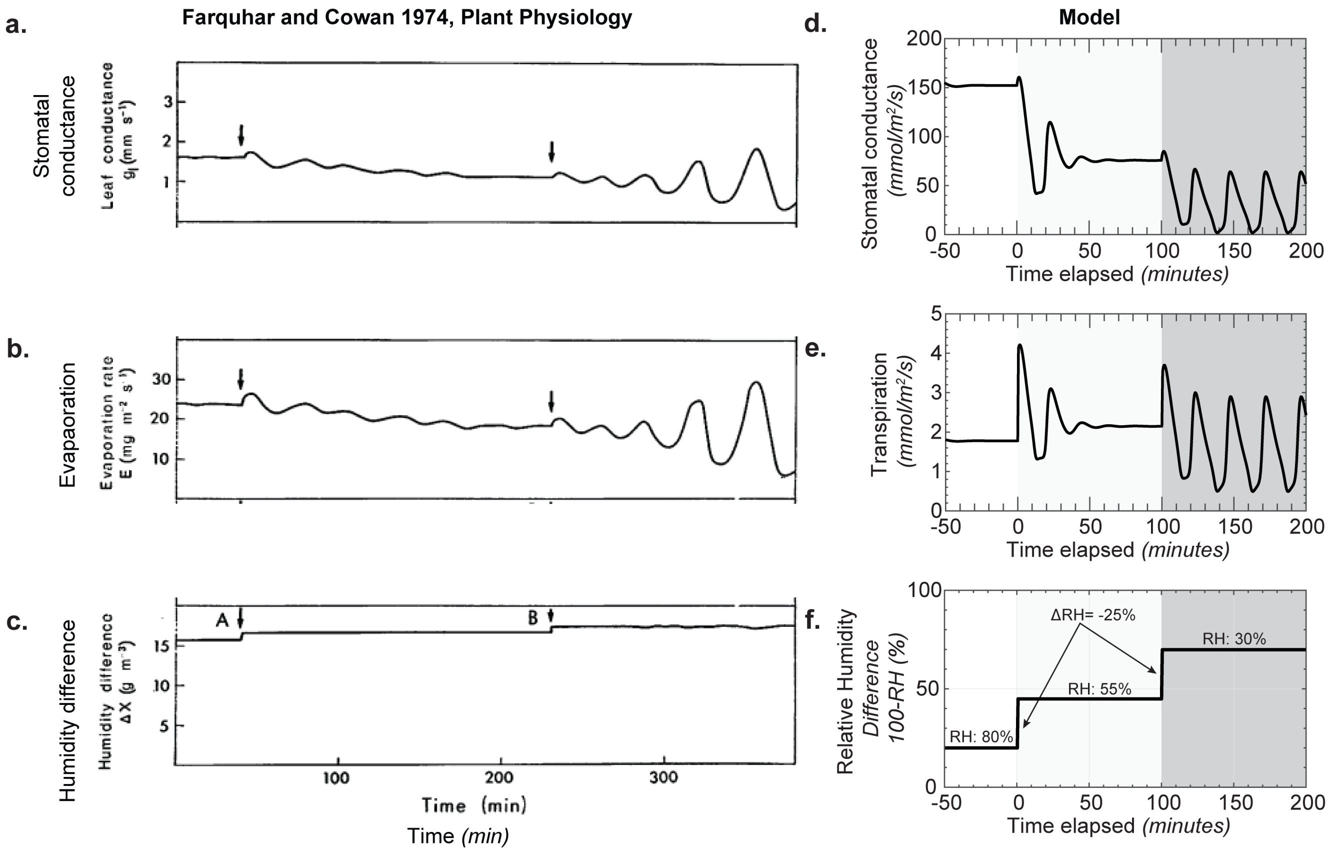

**Fig. S14. Emergence of oscillations in response to step changes in relative humidity: comparison with experimental data.** Panels **a.–c.** are adapted from Farquhar and Cowan[66]. **a.** Stomatal conductance shows a damped oscillatory response to the first step change in relative humidity, followed by increasing oscillatory behavior after a second step change. **b.** Transpiration rate corresponding to the stomatal conductance in (**a**), with panel **c.** showing the imposed relative humidity changes at time points A and B. Panels **d.–f.** are adapted from our model. **d.** Model predictions for two sequential RH perturbations of equal magnitude (25%): first from 80% to 55% at t=0, and then from 55% to 30% at t=100 minutes. **e.** Simulated stomatal conductance exhibits damped oscillations following the first RH drop at t=0, and sustained oscillations following the second drop at t=100. **f.** Corresponding simulated transpiration rate for the stomatal conductance shown in (**e**), capturing the dynamic response to RH perturbations.

**B. Comparing model and experiments on hysteresis.** The dynamic coupled model successfully simulates the transient wild-type stomatal closure following a step increase in VPD from 1.1 kPa to 2.2 kPa (Fig. S15b), validating that the parameter set (Fig. S15a) used for hysteresis is also consistent with experimental transients[62], although showing a slightly higher drop in $g_s$ compared to the parameter set used in Fig. 2, which stems from a larger $V_+/V_{stress}$ ratio. The underlying mechanism involves a drop in guard cell turgor ($P_g$, Fig. S15c), coupled with a rise in guard cell ABA concentration ($[ABA]_g$, Fig. S15d). Elevated $[ABA]_g$ activates the anion channel SLAC1* (Fig. S15e), driving the efflux of solutes. This solute loss results in a decrease in osmotic potential ($\delta\pi_g$, Fig. S15f), leading to stomatal closure.

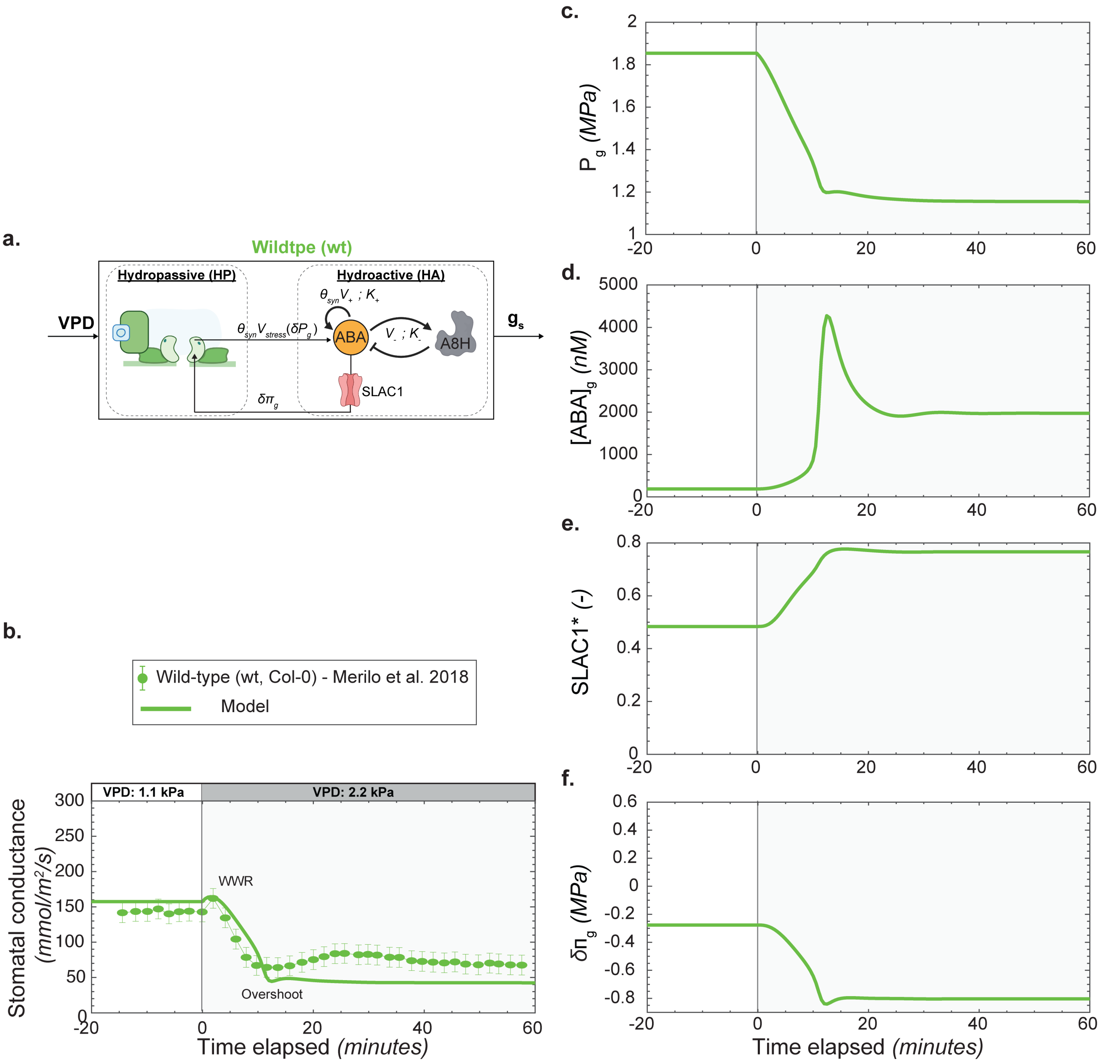

**Fig. S15. Transient stomatal responses in wild type to changes in vapor pressure deficit (VPD) for the hysteresis parameter set used in Fig. 4. a.** Schematic of the full coupled model, highlighting the hydropassive component and the core ABA autoregulation motif. **b.** Predicted and measured transient responses of stomatal conductance $g_s$ following a step increase in VPD from 1.1 to 2.2 kPa. Solid curves show model predictions, and symbols denote experimental data for wild type. Data are from Merilo[62] for this same perturbation in VPD. **c–f.** Model-predicted transients corresponding to panel (b) for: guard cell turgor pressure $P_g$ (c), guard cell ABA concentration $[ABA]_g$ (d), fractional activation of the anion channel SLAC1$^*$ (e), and osmotic potential change in guard cells $\delta\pi_g$ (f). The VPD step corresponds to a change in relative humidity from 80% to 60% at $36^\circ$C. Parameters for the ABA motif were $K_+ = K_- = 1560$ nM, $V_{\text{stress}} = 7.37$ nM/s, $V_+ = 94$ nM/s, and $V_- = 1.5$ nM/s with $\theta_{\text{syn}} = 1$ for wild type. See main text and Section S3 for details on model.

Our model predicts that hysteresis in stomatal conductance arises from bi-stability in ABA dynamics, with two stable ABA concentrations emerging from the nonlinear coupled ODEs that define the ABA regulatory motif (Fig. S6, S12, S13).

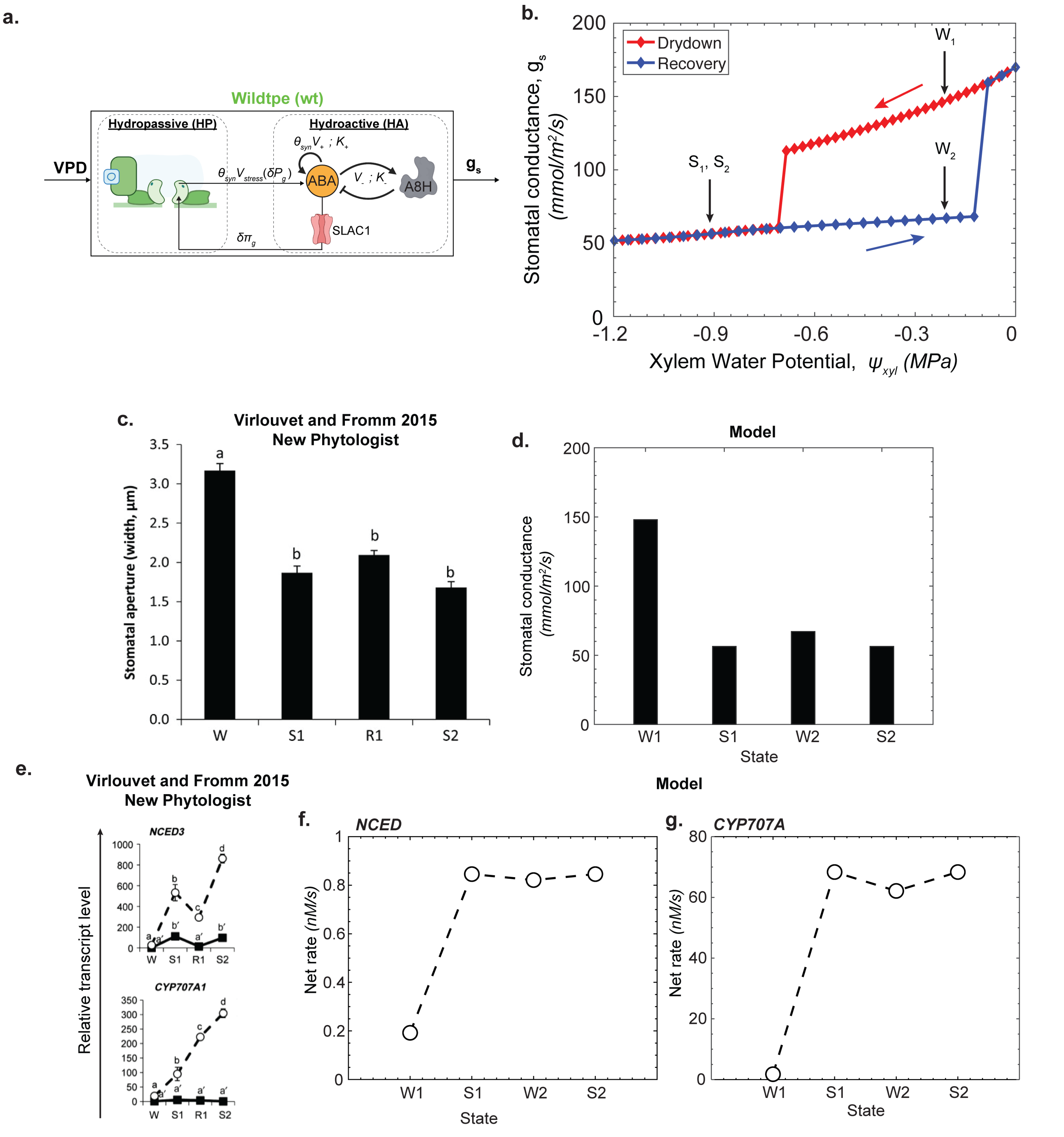

**Fig. S16. Emergence of hysteresis in stomatal response and comparison with experimental data. a.** Trajectories plotted for the parameter set $K_+ = K_- = 1560$ nM, $V_{\text{stress}} = 7.37$ nM/s, and $V_+ = 94$ nM/s, with $\theta_{\text{syn}} = 1$ for wild type. **b.** Steady-state trajectories of stomatal conductance during a sequential drydown (decreasing xylem water potential, $\psi_{\text{xyl}}$; red points) and recovery (increasing $\psi_{\text{xyl}}$; blue points) under constant environmental conditions (RH = 80% at $36^\circ$C; VPD = 1.1 kPa). This panel is reproduced from Fig. 4b of the main text. Water status is categorized as W1–W2 (well-watered) and S1–S2 (water-stressed), with transitions following the sequence W1 $\rightarrow$ S1 $\rightarrow$ W2 $\rightarrow$ S2. **c.** Experimental measurements of stomatal aperture across well-watered states (W, R1) and water-stressed states (S1, S2), adapted from Virlouvet[67]. **d.** Model-predicted stomatal conductance for the same sequence of W1–W2 and S1–S2 transitions. **e.** Relative transcript levels of *NCED3*, *AAO3*, *CYP707A1*, and *CYP707A3* across W, R1, S1, and S2 states, adapted from Virlouvet[67]. Circles denote expression in guard cells; filled squares represent transcript levels in the remainder of the leaf. **f.** Model-predicted rate of ABA synthesis via the rate-limiting enzyme NCED. **g.** Model-predicted rate of ABA catabolism via A8H (CYP707A) during sequential drydown and recovery. Notably, the W1–W2 and S1–S2 state pairs share the same xylem water potential yet exhibit distinct fluxes, highlighting the hysteresis inherent in the system.

To confront our model with experimental results found in literature, in Fig. S16a, we reproduce the model-predicted steady-state trajectories shown in the main text Fig. 4, highlighting well-watered states (W1, W2) and water-stressed states (S1, S2) that share the same xylem water potential.

To investigate the molecular basis of this hysteresis, Virlouvet and Fromm[67] subjected *Arabidopsis thaliana* plants to repeated cycles of dehydration and rehydration by air-drying and rewatering roots. This experimental sequence — $W \rightarrow S1 \rightarrow R1 \rightarrow S2$ produced alternating well-watered (W, R1) and water-stressed (S1, S2) states. However, because xylem water potential was not directly measured in the experiments, the precise water status during stress and recovery phases may not correspond exactly, particularly between W, R1 and S1, S2.

Their data reveal a non-symmetric change in stomatal aperture (Fig. S16c), consistent with our model predictions (Fig. S16d). In parallel, they conducted transcriptomic analysis of key ABA-related genes, including *NCED3*, and *CYP707A1* (Fig. S16e). Gene expression data from isolated guard cells (circles) and whole-leaf tissue (filled squares) showed that *NCED3*, a key enzyme in ABA biosynthesis, exhibits a hysteretic response across dehydration and rehydration cycles. This pattern mirrors the model-predicted hysteresis in ABA flux derived from rate limiting NCED-mediated synthesis. Gene *CYP707A1* expression increased under stress, consistent with our model's assumption that ABA catabolism via *CYP707A1* scales with ABA levels (Fig. S16f.-g.).

Our modeling analysis indicates that the "naïve" stomatal response operates in a monostable regime (Fig. 2 and 3), whereas the "stress-memorized" state exhibiting hysteresis requires a transition to a bi-stable regime. This shift is driven by stronger positive feedback in the ABA autoregulatory loop (Fig. S6). We propose that this requirement for distinct parameterizations is a mathematical recapitulation of transcriptional priming, a phase transition where the system is physically "hard-wired" for a heightened response[68,69].

Biologically, this transition is evidenced by the "super-induction" of the rate-limiting ABA biosynthetic gene, *NCED*. Virlouvet and Fromm[67] demonstrated that guard cells possess a specific transcriptional memory distinct from mesophyll tissue. Upon re-exposure to dehydration, *NCED* transcript levels are significantly higher than during the initial stress[67]. This biological observation—that the machinery for ABA synthesis becomes more responsive after training—provides the mechanistic justification for the increased synthesis gain ($V_+/V_{stress}$) utilized in our hysteresis parameterization.

Mechanistically, this primed state is maintained during recovery by epigenetic remodeling. Genes retain molecular memory marks, specifically elevated H3K4me3 histone modifications and the stalling of RNA Polymerase II (Ser5P Pol II) at promoter regions[68–70]. Consequently, the increased gain $V_+$ in our model is not an arbitrary fit, but the quantitative equivalent of this poised transcriptional machinery. By linking the ABA positive feedback loop to these epigenetic "locking" mechanisms, our model aligns plant water stress memory with the concept of inherent stomatal hysteresis.

Taken together, these experimental observations provide molecular support for our model hypothesis that ABA provides a mechanistic basis for hysteresis in stomatal conductance arising from bi-stability in ABA autoregulation.

### S7. Table of parameters and solution strategy

**A. Table of parameters for hydropassive model.** Table (S4) lists the parameters used for stomatal water relations. The value for pressure-conductance scaling factor is tuned to allow for best match between experimental data and model predictions. The value for parameters are similar to those presented in literature[3,5].

**Table S4. Stomatal biomechanics**

| Parameter | Description | Value | Units | Ref |
|---|---|---|---|---|
| $\chi$ | Pressure-conductance scaling factor | 295 | mmol/MPa/m$^2$/s | 3 |
| $m$ | Mechanical advantage | 1.71 | – | 3 |
| $\phi_g$ | Stomatal density | $150 * 10^6$ | #/m$^2$ leaf area | 71 |

**Table S5. Physiological parameters in hydropassive model**

| Parameter | Description | Value | Units | Ref |
|---|---|---|---|---|
| **Volume per leaf area at reference state** | | | | |
| $V_m^0$ | Mesophyll cell | 1978 | mmol/m$^2$ | 6 |
| $V_s^0$ | Subsidiary cell | 100 | mmol/m$^2$ | 6 |
| $V_g^0$ | Guard cell | 100 | mmol/m$^2$ | 6 |
| **Turgor Pressure at reference state** | | | | |
| $P_m^0$ | Mesophyll cell | 2.2 | MPa | 72 |
| $P_s^0$ | Subsidiary cell | 1 | MPa | 72 |
| $P_g^0$ | Guard cell | 2.5 | MPa | 73 |
| **Osmotic Potential at reference state** | | | | |
| $\pi_m^0$ | Mesophyll cell | 2.3 | MPa | 72 |
| $\pi_s^0$ | Subsidiary cell | 1.1 | MPa | 3 |
| $\pi_g^0$ | Guard cell | 2.6 | MPa | 73 |
| **Modulus of elasticity** | | | | |
| $\varepsilon_m^0$ | Mesophyll cell | 4 | MPa | 72 |
| $\varepsilon_s^0$ | Subsidiary cell | 0.5 | MPa | 72 |
| $\varepsilon_g^0$ | Guard cell | 3.6 | MPa | 73 |
| **Hydraulic Resistances and conductances** | | | | |
| $R_{xm}$ | Xylem - mesophyll | 0.018116 | MPa.m$^2$.s/mmol | 3 |
| $R_{ms}$ | Mesophyll - subsidiary | 5 | MPa.m$^2$.s/mmol | fitted |
| $R_{sg}$ | Subsidiary - guard | 10 | MPa.m$^2$.s/mmol | fitted |
| $g_g^1$ | Guard to leaf air-space | 2.9 | mmol/.m$^2$.s | fitted |
| $g_e^1$ | Subsidiary to leaf air-space | 2.9 | mmol/.m$^2$.s | fitted |
| $g_g^2$ | Guard to atmosphere | 1 | mmol/.m$^2$.s | fitted |
| $g_e^2$ | Subsidiary to atmosphere | 10 | mmol/.m$^2$.s | fitted |
| $g_{oxz}$ | Mesophyll to sub-stomatal cavity | Eq.(S16) | mmol/.m$^2$.s | 18 |

The physiologically possible range for $R_{ms}$, $R_{sg}$ is from $[0.0327 - 32.78]$ $(m^2.s.MPa)/mmol$[74]. The physiologically possible range for $g_g^2$, $g_e^2$ is from $[0 - 10]$ $(mmol/m^2/s)$[75]. We use the values of hydraulic conductances and resistances that best fit the sig (*ost1-3*) mutant stomatal response to VPD change.

### B. Table of parameters for hydroactive model.

1. **Abscisic Acid regulation motif.** See section S3.E and Fig.S6 for details on fitting procedure for parameters in Table S6.

**Table S6. Abscisic acid regulation motif parameters**

| Parameter | Description | Value | Units | Ref |
|---|---|---|---|---|
| $V_{stress}$ | Basal expression of ABA | $8.85$ | nM/s | fitted |
| $V_{+}$ | Saturating expression rate of ABA positive feedback | $67.58$ | nM/s | fitted |
| $V_{-}$ | Basal expression of H | $1.51$ | nM/s | fitted |
| $K_{+}$ | ABA concentration for half-maximal ABA positive feedback | $1560$ | nM | 31 |
| $K_{d}$ | ABA concentration for Michaelis-Menten degradation | $2000$ | nM | fitted |
| $K_{-}$ | ABA concentration for half-maximal H synthesis | $1560$ | nM | fitted |
| $\gamma_A$ | Degradation rate for ABA-H reaction | $0.25$ | 1/s | 31 |
| $\gamma_H$ | Degradation rate for H | $0.0031$ | 1/s | 76 |
| $n$ | Number of self-auto-regulatory genes in ABA biosynthesis pathway | $3$ | – | 29,30 |

2. **Bicyclic signaling cascade.** All kinetic parameters listed in Table S7 were sourced from established literature, with the exception of the SLAC1 phosphorylation rate constant $k_3$ [1/s]. Specifically, $k_3$ dictates the basal open probability of SLAC1 ($P_{open}$), which in turn controls the magnitude of anion efflux. Experimental values for this specific rate were unavailable. We fitted $k_3$ such that this SLAC1-mediated efflux counterbalances the anion uptake driven by the $2H^+/Cl^-$ symporter, thereby establishing a stable baseline for intracellular chloride prior to ABA stimulation.

**Table S7. Futile cycle parameters**

| Parameter | Description | Value | Units | Ref |
|---|---|---|---|---|
| $K_{m1}$ | Michaelis Menten reaction constant for OST1 phosphorylation | 2.5 | $\mu$M | 77 |
| $k_1$ | Reaction rate for OST1 phosphorylation | 0.03 | 1/s | 77 |
| $K_{m2}$ | Michaelis Menten reaction constant for OST1 dephosphorylation | 0.097 | $\mu$M | 78 |
| $k_2$ | Reaction rate for OST1 dephosphorylation | 0.032772 | 1/s | 78 |
| $K_{m3}$ | Michaelis Menten reaction constant for SLAC1 phosphorylation | 23.93 | $\mu$M | 48 |
| $k_3$ | Reaction rate for SLAC1 phosphorylation | 3.75 | 1/s | fitted |
| $K_{m4}$ | Michaelis Menten reaction constant for SLAC1 phosphorylation | 0.97 | $\mu$M | 78 |
| $k_4$ | Reaction rate for SLAC1 dephosphorylation | 0.1386 | 1/s | 78 |

3. **Non-competitive inhibition of PP2C**

**Table S8. Non-competitive inhibition**

| Parameter | Description | Value | Units | Ref |
|---|---|---|---|---|
| $K_D$ | Disassociation constant for ABA with PYR | 1500 | nM | 44,52 |
| $K_I$ | Inhibition disassociation constant for PP2C with ABA:PYR complex | 30 | nM | 44,52 |

4. **Concentration of proteins**

**Table S9. Concentration of proteins**

| Parameter | Description | Value | Units | Ref |
|---|---|---|---|---|
| $[MAP3K]_T$ | Total concentration of MAP3K protein | 1 | $\mu$M | 52,79 |
| $[OST1]_T$ | Total concentration of OST1 protein | 1 | $\mu$M | 52,79 |
| $[PP2C]_T$ | Total concentration of PP2C protein | 1 | $\mu$M | 52,79 |
| $[SLAC1]_T$ | Total concentration of SLAC1 protein | 1 | $\mu$M | 52,79 |
| $[PYR]_T$ | Total concentration of PYR protein | 1 | $\mu$M | 52,79 |

### 5. Guard cell electrophysiology

#### Table S10. Guard cell electrophysiology parameters

| Parameter | Description | Value | Units | Ref |
|---|---|---|---|---|
| **Guard cell electrophysical parameters** | | | | |
| $C_{memb}$ | Membrane capacitance | $0.01$ | F/m$^2$ | 10 |
| $A_g$ | Guard cell surface area | $20 * 10^{-10}$ | m$^2$ | 10 |
| $vol_{in}$ | Guard cell symplast volume | $12$ | pL | 10 |
| $vol_{out}$ | Guard cell apoplast volume | $65$ | pL | 10 |
| **Ion channels** | | | | |
| $\delta_{GORK}$ | Parameter for GORK gating probability | $2$ | – | 10 |
| $K_{GORK}$ | Parameter for GORK gating probability | $150$ | mM | 10 |
| $\delta_{KAT1}$ | Parameter for KAT1 gating probability | $1.6$ | – | 10 |
| $V_{1/2}^{KAT1}$ | Parameter for KAT1 gating probability | $150$ | mV | 10 |
| **Ion concentrations** | | | | |
| $[Cl^-]_{in}^0$ | Concentration of chloride ions in symplast | $200$ | mM | 10 |
| $[Cl^-]_{out}^0$ | Concentration of chloride ions in apoplast | $20$ | mM | 10 |
| $[K^+]_{in}^0$ | Concentration of potassium ions in symplast | $200$ | mM | 10 |
| $[K^+]_{out}^0$ | Concentration of potassium ions in apoplast | $20$ | mM | 10 |
| $pH_{in}^0$ | pH of hydrogen ions in symplast | $7.2$ | – | 10 |
| $pH_{out}^0$ | pH of hydrogen ions in apoplast | $6.5$ | – | 10 |
| **Biophysical parameters for ions** | | | | |
| $z_{Cl}$ | Valence for chloride ion | $-1$ | – | 10 |
| $z_K$ | Valence for potassium ion | $1$ | – | 10 |
| $z_H$ | Valence for hydrogen ion | $1$ | – | 10 |
| $P_{Cl}$ | Permeability of chloride ions through SLAC1 | $1 * 10^{-7}$ | m/s | 10 |
| $P_K$ | Permeability of potassium ions through KAT1 and GORK | $5 * 10^{-6}$ | m/s | 10 |
| **H-ATPase pump** | | | | |
| $E_o$ | H-ATPase maximum charge pumping capacity | $1.1 * 10^6$ | C/m$^2$ | 10 |
| $k_1$ | Parameter for H-ATPase pump | $0.55$ | 1/s | 10 |
| $k_2$ | Parameter for H-ATPase pump | $2.58 * 10^{-4}$ | 1/s | 10 |
| $K_{ABA}$ | ABA dependent pumping probability of H-ATPase pump | $50$ | nM | 60 |
| **2H/Cl symport** | | | | |
| $V_o$ | 2H/Cl symport maximum flux | $9.25 * 10^4$ | C/mM$^3$.s.m$^2$ | 10 |
| **Thermodynamic Parameters** | | | | |
| $\Delta G_{ATP}$ | Free energy from ATP hydrolysis | $-26$ | kJ/mol | 80 |
| $R$ | Universal gas constant | $8.314$ | J/mol.K | 80 |
| $T$ | Temperature | $308$ | K | chosen |
| $F$ | Faraday constant | $96485.3329$ | C/mol | 80 |
| $\bar{\nu}_w$ | Molar concentration of water | $1.8 * 10^{-5}$ | m$^3$/mol | 80 |

### C. Solution strategy.

***C.1. Summary of unknown variables, number of unknowns and governing equations.*** In our model, the variables xylem water potential $\psi_{xyl}$ and atmospheric air partial pressure $w_a$ are imposed independent variables. Below we provide tables associating the unknown variables in the model and the associated governing equations that allow to solve for them.

#### a. Hydraulics

**Table S11. Associating variables and governing equations for hydraulic model**

| Count | Unknown Variable | Governing Equation |
|---|---|---|
| **Water potential** | | |
| 1 | $\psi_m$ | [S37] |
| 2 | $\psi_s$ | [S38] |
| 3 | $\psi_g$ | [S39] |
| 4 | $\psi_{ssc}$ | [S36] |
| **Cell volume** | | |
| 5 | $V_m$ | [S23] written for mesophyll cell |
| 6 | $V_s$ | [S23] written for subsidiary cell |
| 7 | $V_g$ | [S23] written for guard cell |
| **Stomatal conductance** | | |
| 8 | $g_s$ | [S11] |
| **Cell turgor pressure** | | |
| 9 | $P_g$ | [S10] written for guard cell |
| 10 | $P_s$ | [S10] written for subsidiary cell |
| **Vapor pressure** | | |
| 11 | $w_m$ | [S14] written for mesophyll cell |
| 12 | $w_s$ | [S14] written for subsidiary cell |
| 13 | $w_g$ | [S14] written for guard cell |
| 14 | $w_{ssc}$ | [S14] written for SSC airspace |
| **Apoplast water flux** | | |
| 15 | $J_1$ | [S17] |
| 16 | $J_2$ | [S18] |
| 17 | $J_3$ | [S19] |
| **Symplast water flux** | | |
| 18 | $J_m$ | [S24] |
| 19 | $J_s$ | [S25] |
| 20 | $J_g$ | [S26] |
| **Transpiration flux** | | |
| 21 | $E_m$ | [S27] |
| 22 | $E_s^1$ | [S28] |
| 23 | $E_g^1$ | [S29] |
| 24 | $E_s^2$ | [S30] |
| 25 | $E_g^2$ | [S31] |
| 26 | $E_{tot}$ | [S32] |
| **Biochemically active guard cell osmotic adjustment** | | |
| 27 | $\delta\bar{\pi}_g$ | [S99] |
| **Mesophyll conductance to transpiration** | | |
| 28 | $g_{oxz}$ | [S16] |

Note: Eqs. S37-S39 are written using Eqs. S33-S35, hence, we do not count Eqs. S33-S35 in our tally of governing equations.

#### b. ABA synthesis and signaling

**Table S12. Associating variables and governing equations for ABA synthesis and signaling**

| Count | Unknown Variable | Governing Equation |
|---|---|---|
| **ABA synthesis** | | |
| 1 | $[ABA]$ | [S53] |
| 2 | $[H]$ | [S56] |
| 3 | $[ABA:H]$ | [S40] |
| **Futile cycle - MAP3K-OST1-OST1*** | | |
| 4 | $[MAP3K]$ | [S58] |
| 5 | $[OST1]$ | [S60] |
| 6 | $[MAP3K:OST1]$ | [S41] |
| **Futile cycle - PP2C-OST1*-OST1** | | |
| 7 | $[PP2C]$ | [S59] |
| 8 | $[OST1^*]$ | [S72] |
| 9 | $[PP2C:OST1^*]$ | [S42] |
| **Futile cycle - OST1*-SLAC1-SLAC1*** | | |
| 10 | $[SLAC1]$ | [S61] |
| 11 | $[OST1^*:SLAC1]$ | [S43] |
| **Futile cycle - PP2C-SLAC1*-SLAC1*** | | |
| 12 | $[SLAC1^*]$ | [S80] |
| 13 | $[PP2C:SLAC1^*]$ | [S44] |
| **Non-competitive inhibition** | | |
| 14 | $[PYR]$ | [S57] |
| 15 | $[ABA:PYR]$ | [S45] |
| 16 | $[PP2C:ABA:PYR]$ | [S46] |
| 17 | $[PP2C:OST1^*:ABA:PYR]$ | [S47] |
| 18 | $[PP2C:SLAC1^*:ABA:PYR]$ | [S48] |

#### c. Guard cell electrophysiology

**Table S13. Associating variables and governing equations for guard cell physiology**

| Count | Unknown Variable | Governing Equation |
|---|---|---|
| **Membrane voltage** | | |
| 1 | $V_{memb}$ | [S100] |
| **Ion concentration** | | |
| 2 | $[Cl^-]_{in}$ | [S101] |
| 3 | $[K^+]_{in}$ | [S102] |
| 4 | $[Cl^-]_{out}$ | [S97] written for $Cl^-$ |
| 5 | $[K^+]_{out}$ | [S97] written for $K^+$ |
| **Current** | | |
| 6 | $I_{Cl}^{SLAC1}$ | [S82] |
| 7 | $I_K^{GORK}$ | [S84] |
| 8 | $I_K^{KAT1}$ | [S87] |
| 9 | $I_{Hpump}$ | [S89] |
| 10 | $I_{Cl}^{sym}$ | [S95] |

***C.2. Code Availability.*** All MATLAB code used to generate the results in this study is permanently archived in a Zenodo repository at: `https://doi.org/10.5281/zenodo.17888362`. The most current version of the code is also available on GitHub at: `https://github.com/desai-sahil/sys-bio-gs.git`.

The repository is organized into three directories corresponding to the main text figures: `Figure 2 Transients and Steady State Merilo 2018`, `Figure 3 Damped and Sustained Oscillations`, and `Figure 4 Steady State Drydown ABA Hysteresis`. Each directory contains a `README_*.m` file detailing steps to reproduce main text figures. To generate the figures, execute the `main_*.m` script located within each folder.

- **For VPD change**: The code is run for change in VPD with a fixed value of $\psi_{xyl} = -0.1\ MPa$. This fixed value of $\psi_{xyl}$ over a change in relative humidity fixes the value of $g_{oxz}$ in eq. (S16).
- **For hysteresis**: The code is run for change in relative humidity with a fixed value of $RH =\ 80\%$ at $T = 36°C$ and varying the xylem water potential ($\psi_{xyl}$) from 0 $MPa$ to $-1.2\ MPa$. The maximum value of $g_{oxz}$ in Eq. (S16) was used to avoid confounding the effects of non-stomatal regulation from stomatal regulation.

**File inventory**: The provided code archive contains the following key files:

**`main_*.m`**
The primary execution script. Run this file to perform dynamic/steady-state simulations and generate plots for Fig. 2, Fig. 3 and Fig. 4 in the main text.

**`run_fullModel.m`**
A wrapper script that handles the ODE solver calls and post-processing steps (Not required for Figure 4).

**`fullModel_DeltaPg_ode.m`**
The core function defining the system of ordinary differential equations (ODEs) for water potentials, ABA autoregulation, and ion channel dynamics.

**`read_parameters_wt_Merilo_2018.m`**
Script containing the parameter definitions for the Wildtype (WT) genotype used in Fig. 2 and Fig. 3 of main text.

**`read_parameters_wt_hysteresis.m`**
Script containing the parameter definitions for the Wildtype (WT) genotype used in Fig. 4.

**`evaluate_output.m`**
Calculates derived metrics (e.g., stomatal conductance) from the raw ODE solver output.

**`f_*.m`**
Helper functions to do intermediate calculations.

To ensure proper execution, users should verify the following environment settings:

- **Software Version:** MATLAB R2021a or later is required.
- **Path Configuration:** Ensure that all functions and files are added to the MATLAB search path before running the main scripts.
- **Toolboxes:** Standard MATLAB installation is sufficient; no specialized toolboxes are required for the core ODE solver.
